# A Comprehensive Atlas of AAV Tropism in the Mouse

**DOI:** 10.1101/2024.09.10.612279

**Authors:** Christopher J. Walkey, Kathy J. Snow, Jote Bulcha, Aaron R. Cox, Alexa E. Martinez, M. Cecilia Ljungberg, Denise G. Lanza, Marco De Giorgi, Marcel A. Chuecos, Michele Alves-Bezerra, Carlos Flores Suarez, Sean M. Hartig, Susan G. Hilsenbeck, Chih-Wei Hsu, Ethan Saville, Yaned Gaitan, Jeff Duryea, Seth Hannigan, Mary E. Dickinson, Oleg Mirochnitchenko, Dan Wang, Cathleen M. Lutz, Jason D. Heaney, Guangping Gao, Stephen A. Murray, William R. Lagor

**Affiliations:** Department of Integrative Physiology, Baylor College of Medicine, Houston, TX 77030, USA; The Jackson Laboratory, 600 Main Street, Bar Harbor, ME 04609 USA; Horae Gene Therapy Center and Department of Microbiology and Physiological Systems, University of Massachusetts Medical School, Worcester, MA 01605, USA; Li Weibo Institute for Rare Diseases Research, University of Massachusetts Medical School, Worcester, MA 01605, USA; Section of Diabetes, Endocrinology, and Metabolism, Department of Medicine, Baylor College of Medicine, Houston, TX 77030, USA; Department of Pediatrics, Baylor College of Medicine, Houston, TX 77030, USA; Duncan Neurological Research Institute, Texas Children’s Hospital, Houston, TX 77030, USA; Department of Molecular and Human Genetics, Baylor College of Medicine, Houston, Texas, USA; Department of Molecular and Cellular Biology, Baylor College of Medicine, Houston, TX 77030, USA; Dan L. Duncan Comprehensive Cancer Center, Baylor College of Medicine, Houston, Texas, USA; Office of Research Infrastructure Programs, Division of Program Coordination, Planning, and Strategic Initiatives, Office of the Director, National Institutes of Health, Bethesda, MD USA; RNA Therapeutics Institute, University of Massachusetts Medical School, Worcester, MA 01605, USA; Current address: 4D Molecular Therapeutics, Emeryville, CA 94608, USA; Current address: Institute of Molecular Medicine, University of Texas Health Science Center at Houston, Houston, TX 77030, USA; Current address: Department of Neuroscience, Baylor College of Medicine, Houston, TX 77030, USA; Current address: Department of Biomedicine, Biotechnology and Public Health, Biomedical Research and Innovation Institute of Cadiz (INiBICA), University of Cadiz, 11002 Cadiz, Spain; Current address: Department of Experimental Therapeutics, MD Anderson Cancer Center, Houston, TX 77030, USA; Current address: The Jackson Laboratory, 600 Main Street, Bar Harbor, ME, 04609 USA

## Abstract

Gene therapy with Adeno-Associated Viral (AAV) vectors requires knowledge of their tropism within the body. Here we analyze the tropism of ten naturally occurring AAV serotypes (AAV3B, AAV4, AAV5, AAV6, AAV7, AAV8, AAV9, AAVrh8, AAVrh10 and AAVrh74) following systemic delivery into male and female mice. A transgene expressing ZsGreen and Cre recombinase was used to identify transduction in a cell-dependent manner based on fluorescence. Cre-driven activation of tdTomato fluorescence offered superior sensitivity for transduced cells. All serotypes except AAV3B and AAV4 had high liver tropism. Fluorescence activation revealed transduction of unexpected tissues, including adrenals, testes and ovaries. Rare transduced cells within tissues were also readily visualized. Biodistribution of AAV genomes correlated with fluorescence, except in immune tissues. AAV4 was found to have a pan-endothelial tropism while also targeting pancreatic beta cells. This public resource enables selection of the best AAV serotypes for basic science and preclinical applications in mice.

## Introduction

Gene therapy requires efficient and specific delivery of the nucleic acid cargo to the tissue and cell type of interest. Over the past three decades, adeno-associated viruses (AAVs) have emerged as the leading viral vector for *in vivo* gene therapy (Pupo et al., 2022). AAV vectors have been tested in more than 250 clinical trials, culminating in six FDA approved products at the time of this report. AAV are non-cytopathic and can transduce both dividing and non-dividing cells (Wu et al., 2006). AAV vectors elicit modest and generally manageable immune responses relative to other viral vectors (Rabinowitz et al., 2019; Zinn and Vandenberghe, 2014). Additionally, the AAV genome persists in cells mostly as non-integrating circular episomes, which partially mitigates the risk of chromosomal insertions and genotoxicity (Sabatino et al., 2022). Despite this progress, our knowledge of AAV biology remains incomplete. An important goal is to improve the therapeutic utility of AAVs by broadening the range and specificity of tissues and cell types that can be targeted for therapy.

AAVs, members of the *Parvoviridae* family, consist of a 4.7 kb single-stranded DNA genome (Srivastava et al., 1983), surrounded by a 25 nm diameter non-enveloped protein capsid (Atchison et al., 1965). AAVs are dependoviruses that require a helper virus, such as adenovirus or a herpes virus, for replication (Geoffroy and Salvetti, 2005). The wild-type AAV genome contains two genes *rep* and *cap,* flanked on either side by ∼145 nucleotide hairpin like structures called Inverted terminal repeats (ITRs) (Srivastava et al., 1983). The *rep* gene encodes four proteins involved in replication, encapsidation, and integration of the viral genome. The *cap* gene produces three different capsid proteins, VP1, VP2 and VP3 (Rose et al., 1971). The VP capsid proteins assemble into an icosahedral structure of sixty total subunits, in a 1:1:10 ratio(Snijder et al., 2014). Three accessory proteins-Assembly Activating Protein (AAP) (Sonntag et al., 2010), Membrane Associated Activating Protein (MAAP) (Ogden et al., 2019), and protein X (Hermonat et al., 1999), are also encoded in an alternate open reading frame within *Cap*.

Recombinant AAV (rAAV) are generated by replacing the *rep* and *cap* genes with a transgene expression cassette, flanked by the ITRs. In principle any transgene cassette up to ∼4.9 kb can be packaged into rAAV, enabling treatment of many, but not all, gene therapy targets.

The process of AAV transduction involves multiple sequential events including: *1)* trafficking to the target tissue, *2)* initial interaction with glycans at the cell surface, *3)* binding to an internalization receptor, *4)* retrograde transport from plasma membrane to the trans Golgi network, *5)* escape from the endosome, *6)* avoidance of degradation in the cytoplasm, *7)* nuclear import, *8)* uncoating to release the single-stranded DNA genome, *9)* second strand synthesis and formation of circular DNA episomes, *10)* transcription and translation of the transgene (Riyad and Weber, 2021). All steps of this complex itinerary are directed by interactions between the capsid and host factors on the cell surface or inside the target cell. Accordingly, the amino acid sequence of the *Cap* gene is the single most important determinant of AAV tropism.

At least 13 distinct AAV serotypes encompassing hundreds of naturally occurring genomic variants have been isolated, from multiple vertebrate species (Gao et al., 2004; Loeb et al., 2024). AAV serotypes differ in their *cap* sequence, with the greatest diversity on surface-exposed variable loop regions that participate in receptor binding and antigenicity. AAV’s first interaction with target cells involves glycans present on numerous plasma membrane proteins. Glycan binding preferences are known for several serotypes including-heparan sulfate proteoglycan for AAV2 (Kern et al., 2003), N-linked galactose for AAV9 (Bell et al., 2012), α2–3 or α2-6 N-linked sialic acid for AAV5 (Kaludov et al., 2001), and α2–3 O-linked sialic acid for AAV4 (Kaludov et al., 2001). For some serotypes, specific proteins have also been identified as co-receptors such as αVβ5 integrin (Summerford et al., 1999) for AAV2, PDGFR (Di Pasquale et al., 2003) for AAV5, and laminin receptor (LamR) (Akache et al., 2006) for AAVs2, 3 and 9. Most naturally occurring AAV serotypes, excluding AAV4 and AAVrh32.33 (Dudek et al., 2018), require a common receptor called AAV Receptor (AAVR) to enter cells (Pillay et al., 2016). AAVR binds to conserved regions in the AAV capsid through its IgG-like polycystic kidney disease (PKD) domains and facilitates endocytosis (Silveria et al., 2020). Most AAV serotypes also require interactions with GPR108 to escape from the trans Golgi network, with AAV5 as a notable exception(Dudek et al., 2020).

The tropism of recombinant AAV vectors *in vivo* is determined by a complex interplay of many factors including: sex, age, dose, route of administration, residence time in the circulation, relative abundance of glycan and internalization receptors, competition between tissues, interaction with the innate and adaptive immune systems, and cell type-dependent pathways. It should be noted that AAV transduction in tissue culture is inefficient for most serotypes, and does not predict *in vivo* tropism. Tropism can also vary greatly across species. For example, AAV3B is exceptionally poor at transduction of mouse liver, but one of the best serotypes for human hepatocytes (Biswas et al., 2020; Wang et al., 2015). Likewise, the engineered AAV9 peptide insertion variants AAV-PHP.B which crosses the blood brain barrier following systemic delivery, are effective only in mouse strains expressing the Ly6a receptor (Huang et al., 2019). Despite important species differences, common themes do emerge, which have greatly improved our understanding of AAV biology and vectorology.

The mouse remains the primary model for initial preclinical testing of AAV gene therapies. The vast majority of studies have focused on a limited subset of commonly used human and primate AAV isolates due to their likelihood for effective translation to humans. In a seminal paper, Zincarelli et al. performed the most extensive analysis of AAV tropism in mice to date (Zincarelli et al., 2008). The authors profiled the biodistribution and expression of 9 AAV serotypes (AAV1-9) across 9 tissues and 6 serial timepoints following tail vein injection of a luciferase reporter vector. This study revealed a strong tropism for the liver for almost all serotypes, with varying kinetics of expression. The value of such well controlled comparative studies cannot be understated. However, the study design did not allow for a detailed examination of transduction at the level of individual cells, and lacked a broad survey of organs and tissue types. It also focused only on male mice.

In the current work, we performed a comprehensive survey of AAV tropism in male and female mice following systemic administration using fluorescent reporter activation. We define the biodistribution and functional transduction of ten naturally occurring AAV serotypes across 22 different tissues, including reproductive organs, and hematopoietic stem cells. By using fluorescent protein activation as a readout of transduction, we were able to detect rare events, and in some cases identify specific target cell types. We have identified novel tropisms for multiple serotypes, most notably AAV4 which broadly delivers to endothelial cells in most tissues as well as beta cells in the pancreas. Our results have generated a comprehensive atlas of murine AAV tropism that is freely available to researchers interested in using AAVs as gene delivery vehicles *in vivo*. Data and images associated with these studies are publicly available at https://scge.mcw.edu/toolkit. This resource will help in the selection of the best serotype for preclinical gene therapy studies, and open new doors for previously untargetable cell types.

## Results

### Vector design and approach

To date, most surveys of AAV tropism in mice have examined only a few serotypes focusing on small subset of possible tissues, using readouts that lack cell type specific resolution. To comprehensively evaluate AAV tropism in mice, we designed an AAV vector with a bi-directional chicken beta actin (CB) promoter with a CMV enhancer element expressing both ZsGreen and Cre recombinase (**Figure 1A**) (Lahey et al., 2020). Recent work indicates that Cre-mediated activation of a genomically encoded fluorophore is a more sensitive method of detecting AAV transduction than AAV-mediated delivery (Lang et al., 2019). We chose to employ the Ai9 reporter mouse strain, which harbors a CAG-driven tdTomato bright red fluorescent reporter gene at the *Rosa26* locus(Madisen et al., 2010). The Ai9 allele contains a lox-STOP-lox cassette which can be removed by Cre, permanently activating tdTomato expression. We reasoned that delivery of this bidirectional reporter into Ai9 mice would allow us to simultaneously track cells with active transgene expression (ZsGreen), and those cells with a history of Cre expression (tdTomato) (**Figure 1B**). ZsGreen levels are expected to vary as a function of AAV copy number per cell, while tdTomato provides a binary readout of transduction with a particularly bright and uniform signal, also marking daughter cells. Additionally, biodistribution of the AAV genome can be measured for a quantitative readout of tissue uptake in relation to transgene expression.

**Figure 1.**
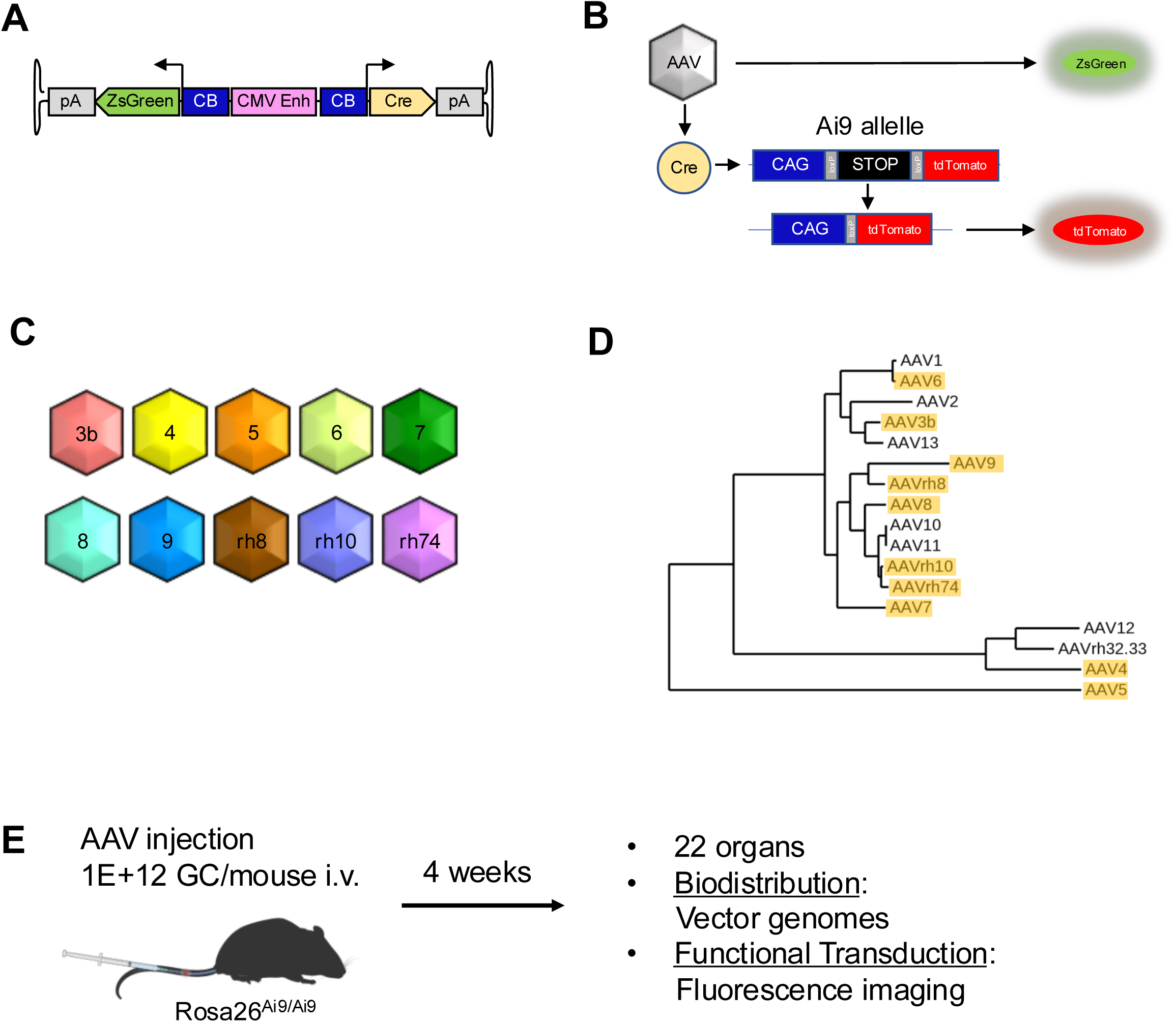
Experimental design for the analysis of AAV tropism survey in mice. **A)** AAV vector design-CMV enhancer element/bidirectional CB promoter allows for expression of both Cre and ZsGreen from the same AAV vector. **B)** Detection -Cells transduced with AAV will express both the zsGreen fluorescent reporter and Cre recombinase to activate the Ai9 allele. ZsGreen fluorescence reports levels of active AAV transgene expression, while TdTomato fluorescence permanently marks cells with a history of transduction. **C)** Serotypes - The transgene cassette was packaged into the indicated capsid serotypes. **D)** Phylogenetic comparison of AAV serotypes – VP1 capsid protein sequences relationships between different natural AAV serotypes. Serotypes used in this study are highlighted in orange. **E)** Experimental workflow - For each serotype, two cohorts of 4 to 6 male and 4 to 6 female mice were injected intravenously and dissected four weeks later. From one cohort, genomic DNA was purified from a panel of organs for quantitative PCR analysis. From the second cohort, the same organs were prepared and imaged by fluorescent microscopy.

AAV serotypes were chosen to reflect a diversity of capsid sequences (**Figure 1C**). As secondary criteria, we restricted our choices to natural serotypes of highest preclinical and/or clinical relevance. Our final list included: AAV3b, AAV4, AAV5, AAV6, AAV7, AAV8, AAV9, AAVrh8, AAVrh10 and AAVrh74 (**Figure 1C**) (Chiorini et al., 1999; Chiorini et al., 1997; Gao et al., 2003; Gao et al., 2004; Gao et al., 2002; Rutledge et al., 1998). **Figure 1D and Supplemental Table 1** illustrate the relationship between capsid amino acid sequences. Our experimental workflow is outlined in **Figure 1E**. Five-week-old mice, homozygous for the Ai9 allele at the *Rosa26* locus, were injected intravenously through the tail vein. This route of administration was chosen to allow even distribution throughout the body. The AAV dosage was 1E12 genome copies (GC) per mouse, equivalent to ∼5E13GC/kg, intended to maximize ability to detect extra-hepatic transduction. To reveal sexual dimorphism with respect to AAV tropism, and provide sufficient power for statistical analysis, four to six mice of each sex were injected. Four weeks after injection, a panel of twenty-two organs was harvested and prepared for either DNA isolation or fluorescent imaging **(Supplemental Table 2)**.

### Biodistribution

Total DNA was prepared from each organ sample and assayed for the viral genome copy number by quantitative PCR. Heat map summaries of the results are presented in **Figure 2A**, while the same data for a subset of individual serotypes are shown in **Figure 2B** and **Supplemental Tables 3 and 4.** As expected for an intravenous injection, most serotypes (AAV5,6,7,8,9,rh8,rh10 and rh74) displayed the strongest tropism for liver in both male and female mice, with values exceeding 10^6^ GC/μg of genomic DNA. Significant levels of AAV vector genomes were also observed in samples from adrenal glands, gastrocnemius muscle, diaphragm, heart, and skin. Two serotypes were notable exceptions to this pattern. AAV3B was found to have low transduction overall, with a total combined delivery to all organs of ∼ 10^5^ GC/μg DNA and did not target the liver. AAV4 displayed a significant tropism for the lung (1.3 x 10^6^ GC/μg DNA for males and 2.3 x 10^6^ GC/μg DNA for females) and did not appear to transduce the liver effectively. These results closely match those found previously (Zincarelli et al., 2008).

**Figure 2.**
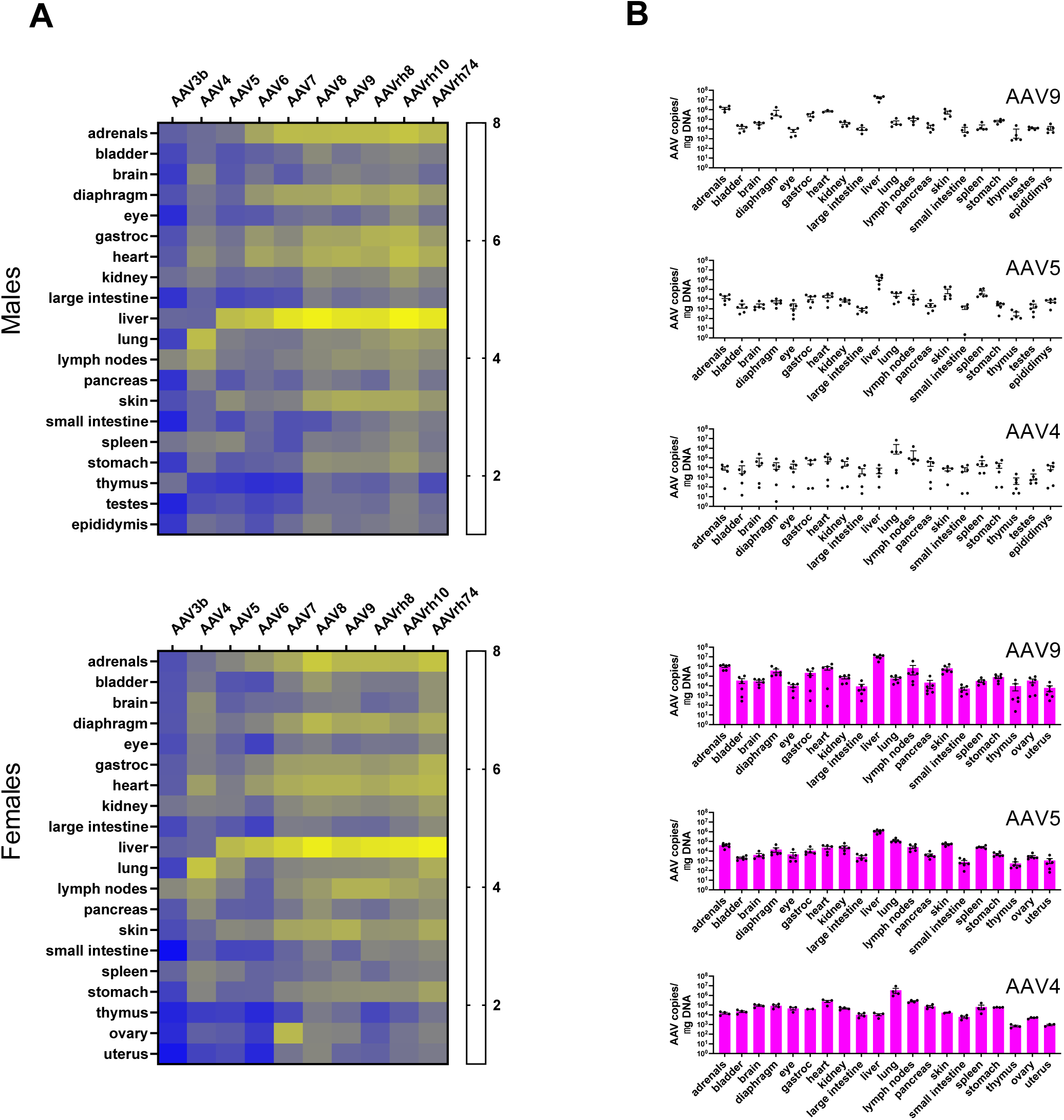
Biodistribution of 10 AAV serotypes across 22 different tissues. **A)** Viral biodistribution summary - Total DNA was isolated from each organ in individual mice. Absolute copy numbers of viral genomes were measured by quantitative PCR, and calculated as AAV genome copies (GC) per microgram of input DNA and shown on a log_10_ scale. N= 4 to 6 for each serotype and each sex. **B)** Sample tissue comparison - qPCR results are plotted to show all organs assayed for a particular serotype. Results from male (blue) and female (pink) mice injected with AAV9, AAV5 and AAV4 are shown, with somatic organs listed in alphabetical order, and reproductive organs at the end. Error bars represent +/- SEM.

This comprehensive survey of serotypes allows for quantitative comparisons of delivery within a single organ and across sexes. **Supplemental Figure 1** and **Supplemental Table 5** identify the serotypes that produced the highest vector genome copy number in each organ. For example, in lung, AAV4 produced the highest viral genome copy number in both male and female mice. In a similar manner, AAV8 was the most effective serotype for transducing bladder. Interestingly, there was some discordance between male and female mice with respect to the best performing serotypes. In male mice, AAVrh10 produced the highest combined viral genome copy number (5.1 x 10^7^ GC/μg DNA), as well as being the top scorer for fourteen of twenty organs. In female mice, AAVrh74 produced the highest combined viral load (5.1 x 10^7^ GC/μg DNA) and was the top scorer in nine of twenty organs. Overall, some differences were observed between male and female mice **(Supplemental Figure 2)**, including more efficient transduction of liver in males versus female mice, as previously noted (Davidoff et al., 2003; Pañeda et al., 2009).

### Functional transduction

The ability of an AAV to mediate functional transduction resulting in fluorescent protein expression was assessed by imaging. As mentioned above, we employed two simultaneous strategies to produce fluorescent reporter molecules: direct fluorescence by expression of ZsGreen from the vector, and Cre-mediated activation of tdTomato from the Ai9 allele. Initial examination of the images revealed that significantly more cells expressed Cre-activated tdTomato than ZsGreen. This phenomenon was consistent through all serotypes and all tissues. A sample of these images comparing ZsGreen and tdTomato expression in tissues transduced with AAV4 and AAV9 is shown in **Supplemental Figure 3**. Given the dramatically improved sensitivity of tdTomato following Cre activation, we restricted subsequent analyses to this reporter.

Representative images from organs with an average high (liver), medium (adrenal glands), and low (thymus) transduction efficiency across serotypes are shown for all ten serotypes in **Figure 3**. Images for all ten AAV serotypes across the 22 different organs/tissues are shown in **Supplemental Figure 4**. As expected from the biodistribution data, a strong tdTomato signal was observed in the liver for most serotypes with the exceptions of AAV3B and AAV4 (**Figure 3A**). This result provides an initial demonstration of the concordance between viral genome copy number and functional transduction measured by a fluorescent reporter. Likewise, tdTomato fluorescence images from the adrenal glands matched the biodistribution data. In this case, fluorescent imaging revealed a consistent pattern of tdTomato expression in the cortex, especially towards the center of the organ, while the medulla was relatively untransduced (**Figure 3B**). This result demonstrates the advantage of using fluorescent imaging to detect transduction: the ability to identify a particular region or cell type with a different AAV tropism within the intact organ. Finally, despite the low viral genome copy in several tissues for a variety of serotypes, Cre-mediated tdTomato expression was observed in a handful of cells, for example in the thymus with AAV6 and AAV7 (**Figure 3C**). Likewise, although fluorescent imaging revealed transduction in both testes and ovaries, the activated cells appeared somatic based on morphology and location (**Figure 4**). In testes, tdTomato-positive cells for all serotypes appeared exclusively outside the seminiferous tubules (i.e. within the interstitial space), as defined by DAPI staining (**Figure 4A**). In the ovary, tdTomato-positive cells appeared solely outside the follicles (i.e. within the medulla) (**Figure 4B**). Therefore, our imaging did not identify transduction of germ cells by any serotype. Hence, our technique allows detection of rare transduction events in single cells which may nonetheless be biologically relevant, particularly for immune cells and the germ line.

**Figure 3.**
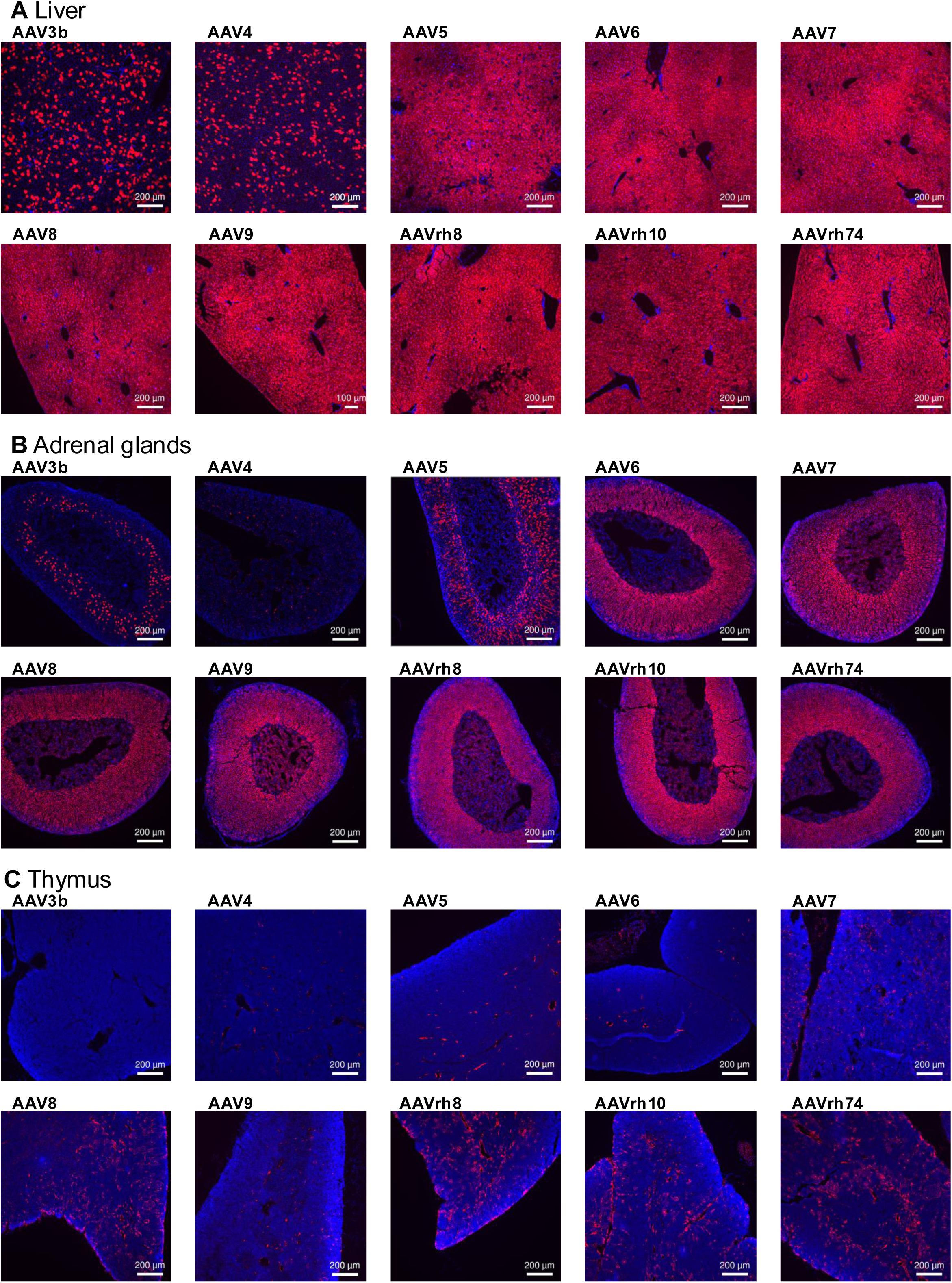
Functional transduction detected by fluorescent imaging: Organ samples were prepared and imaged for fluorescent tdTomato (red) indicative of functional transduction. Nuclei were stained for DAPI (blue). Representative images from **(A)** liver (high transduction), **(B)** adrenal glands (medium transduction) and (**C)** thymus (low transduction) for each serotype are shown.

**Figure 4.**
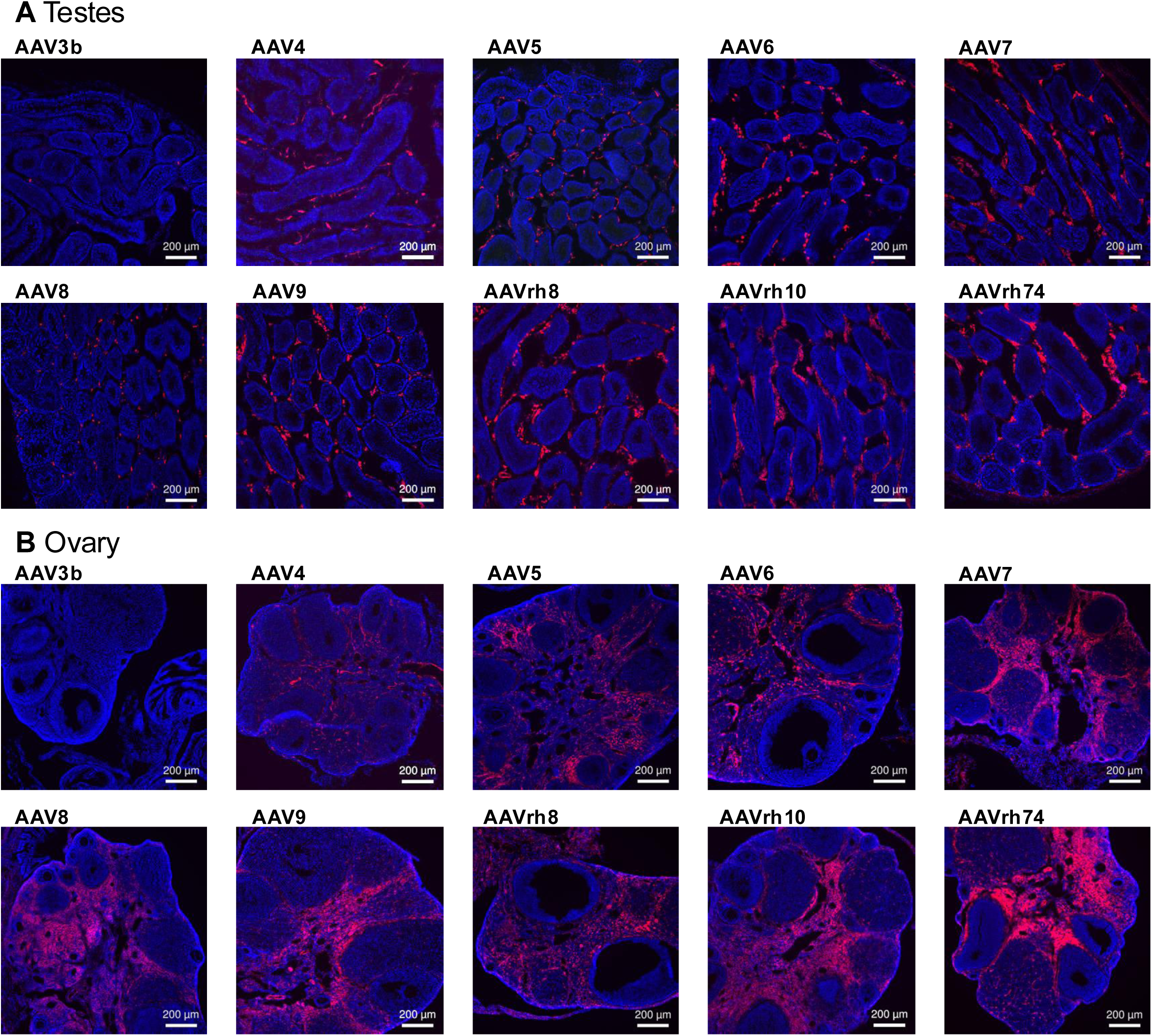
Functional transduction of testes and ovary. tdTomato fluorescence (red) and DAPI nuclei (blue) for tissues from mice transduced with the indicated serotypes in **A)** testes and **B)** ovary.

To quantify functional transduction, we developed a novel image scanning technique that quantifies tdTomato+ red fluorescence per total area. Our results are summarized in **Figure 5** and **Supplemental Tables 6 and 7**. Notably, these results appear to closely match the viral biodistribution data from **Figure 2**. For both males and females, liver is the most heavily transduced organ for serotypes for AAV5, 6, 7, 8, 9,r h8, rh10 and rh74-reaching the maximum value of 100%. Adrenal gland, heart, and muscle (diaphragm and gastrocnemius) also produced strong signal for these serotypes. For AAV4, lung had the highest percentage of tdTomato-positive cells in males, as expected from our biodistribution data. For females transduced with AAV4, heart and diaphragm showed a slightly higher percentage of tdTomato-positive cell and fiber area than lung (**Figure 5**, **Supplemental Figure 5)**.

**Figure 5.**
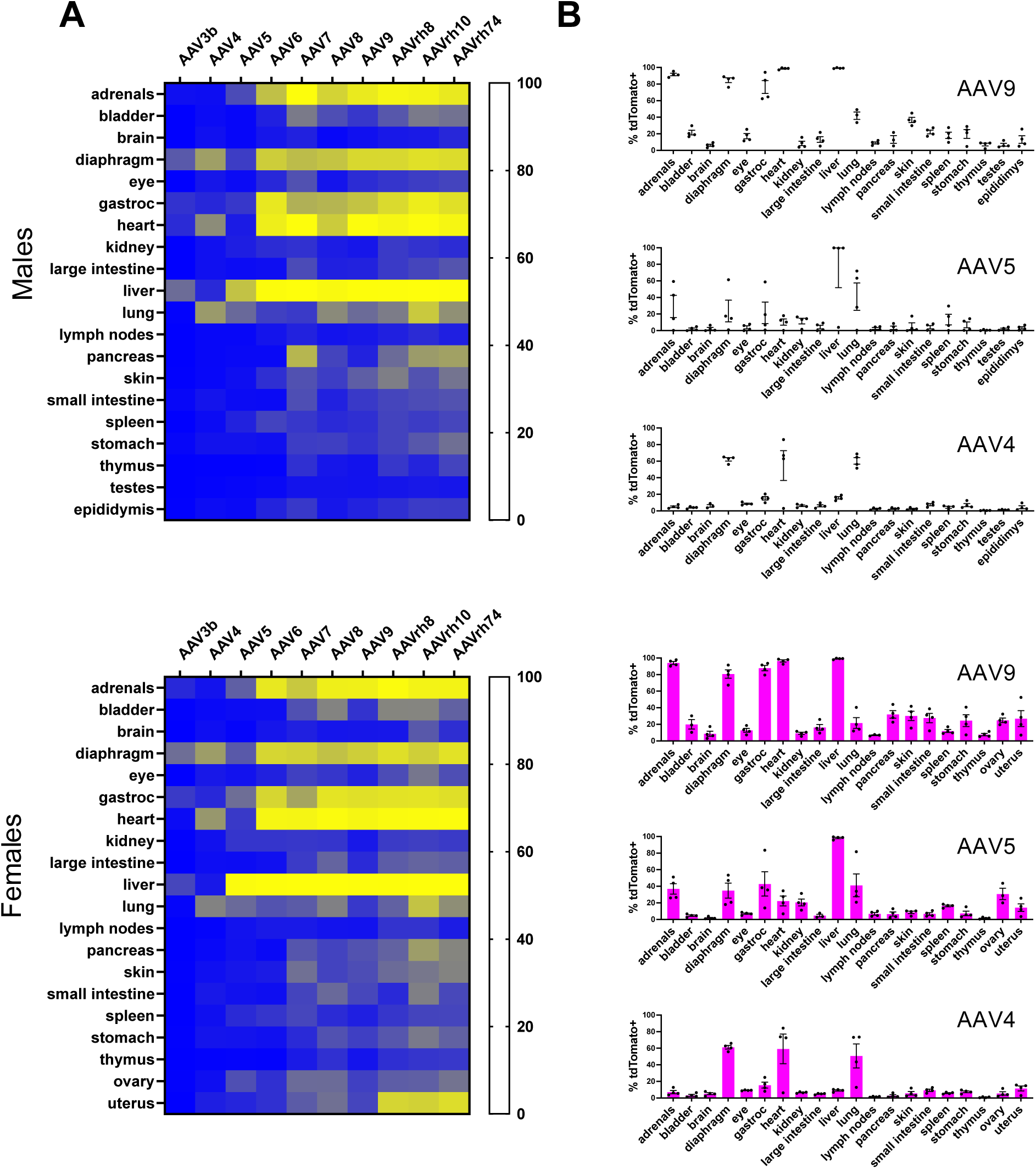
Quantification of functional transduction across tissues. **A)** Image quantification -tdTomato signal in each image was measured to quantify transduction efficiency from four images for each organ, serotype and sex, and expressed as a percentage of maximum coverage (N=4) Results are shown as a heat map. **B)** Sample tissue comparison - Image quantification results are plotted to show all organs assayed for a particular serotype. Results from male (blue) and female (pink) mice injected with AAV9, AAV5 and AAV4 are shown, with somatic organs listed in alphabetical order, and reproductive organs at the end. Error bars represent +/- SEM.

Nonetheless, some differences between the viral biodistribution data and functional transduction were observed. For example, AAVrh8, rh10 and rh74 produced a significant number of tdTomato-positive cells in the uterus (**Figure 5A**, **Supplemental Figures 4 and 5)**, despite a relatively low viral genome copy number (**Figure 2A**). Therefore, we sought to establish an overall correlation for all tissues and serotypes between viral genome copy number and tdTomato-positive cells **(Supplemental Figure 6)**. The results indicate a strong overall correlation between viral genome copy number and tdTomato+ cells. Interestingly, the data produced sigmoidal curves. At the low end, this result suggests that a certain threshold of viral particles can enter a tissue before tdTomato allele activation by Cre occurs. At the high end, the curve flattens once tdTomato-positive cells reach 100%, and further transduction cannot be registered. Graphs with the same correlation data for individual organs are shown in **Supplemental Figure 7**. For most tissues, a direct relationship between viral genome copies and tdTomato positive cells is evident. However, three organs: spleen, thymus, and lymph nodes, appear to show reasonably high vector genome copy number without a corresponding amount of tdTomato-positive cells. This apparent non-functional transduction (DNA delivery without transgene expression) may be related to the immune function of these organs, or reflect promoter-specific silencing in these tissues.

### AAV4 has a pan-endothelial tropism and transduces pancreatic islets

The tropism of AAV4 differed significantly from all the other serotypes tested, with poor liver transduction and a very strong signal in lung. This less studied AAV serotype is one of the most distinct in terms of capsid sequence, and has a unique preference for α2–3 O-linked sialic acid (Kaludov et al., 2001). AAV4 was reported to exhibit a cardiopulmonary tropism based on vector genomes and luciferase imaging(Shen et al., 2013), but specific cell types were not identified. To further interrogate cell type specificity, we generated AAV4 vectors expressing Cre recombinase, driven by either the chicken beta actin promoter with CMV enhancer, or the EF1 alpha promoter. These vectors did not express ZsGreen, allowing a broader range of fluorescent markers to be used for analysis. Intravenous delivery of AAV4-Cre to Ai9 mice resulted in substantial tdTomato-positive signal in lung outside the airways (**Figure 6A**). Flow cytometry revealed that the majority of tdTomato signal originated from endothelial cells based on CD31+/CD45- expression. Remarkably, >85% of lung endothelial cells were transduced by AAV4 in both male and female mice. In contrast, only ∼20% of non-immune, non-endothelial cells in the lung appeared to be transduced by AAV4 (**Figure 6B**).

**Figure 6.**
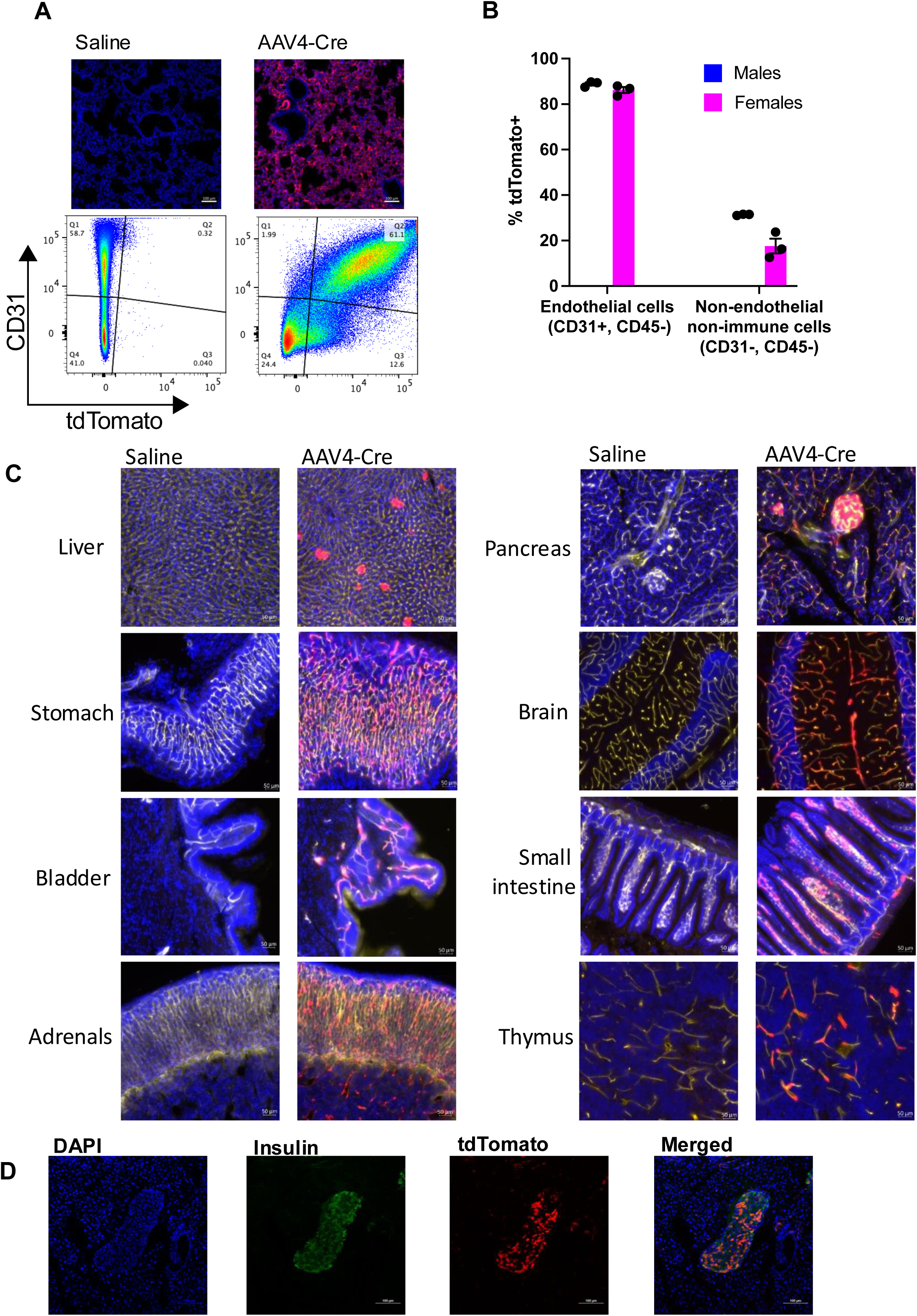
AAV4 exhibits a unique tropism for endothelial cells and pancreatic beta cells. **A)** Transduction of pulmonary endothelial cells by AAV4 - Fluorescent images and flow cytometry plots showing induction of tdTomato expression (upper right quadrant) in endothelial cells (CD31+, CD45-) following injection with AAV4-Cre. Three male and three female mice were examined four weeks after receiving a dose of 5E11 GC/mouse. One male and one female mouse were injected with saline as negative controls. **B)** Quantification of pulmonary endothelial cell transduction flow cytometry results. Error bars represent +/- SEM. **C)** Transduction of endothelial cells in multiple tissues - Tissue sections from negative control and AAV4-Cre injected mice are shown, with DAPI staining for nuclei (blue), DyLight649-lectin staining for endothelial cells (yellow) and tdTomato (red). Images show coincident tdTomato expression in lectin-stained endothelial cells. **D)** Pancreas sections were stained with a fluorescently-labelled insulin antibody and imaged by confocal microscopy. Coincident insulin staining (green) and tdTomato fluorescence (red) confirm AAV4 transduction of beta cells within pancreatic islets.

These striking results prompted us to ask whether AAV4 might also transduce endothelial cells in other vascular beds, a historically neglected cell type for gene therapy applications. The experiment was repeated using an AAV4 vector expressing Cre driven by the EF1alpha promoter. To label the lumenal endothelial surface of the vasculature, DyLight649-conjugated tomato lectin was injected intravenously prior to dissection. Forty micron sections from multiple organs were prepared and imaged for the fluorescent lectin and tdTomato. As expected, liver was poorly transduced with AAV4 showing only sparse TdTomato hepatocytes and possible signal in isolated liver sinusoidal endothelial cells. In stark contrast, impressive transduction of endothelial cells was seen in the stomach, bladder, adrenals, pancreas, brain, small intestine, and thymus (**Figure 6C**). Clear and distinct endothelial signal was also observed in the large intestine, muscle, lymph nodes, skin, ovary, and uterus **(Supplemental Figure 8)**. In separate experiments, we also confirmed strong endothelial delivery to the retina **(Supplemental Figure 9A)**. AAV4 also mediated transduction of the largest vessel in the body, hitting 26% of endothelial cells in the thoracic aorta **(Supplemental Figure 9B)**.

In addition to its apparent pan-endothelial tropism, we also noted strong signal in pancreatic islets with AAV4 **(Supplemental Figure 3A)**. To confirm this result, we performed a second experiment with a Cre-only AAV4 construct. Co-staining with an antibody to insulin confirmed efficient transduction of beta cells with AAV4 (**Figure 6D**). We re-examined other serotypes for transduction of pancreatic islets in our initial experiments. Although AAV8 appeared to have some tropism for islets, it was much less specific, with strong tdTomato signal from surrounding acinar cells following intravenous delivery **(Supplemental Figure 10)**. Strong transduction of pancreatic acinar cells was seen for AAV7,9,rh8,rh10, and rh74. AAV5 also appeared to also transduce islets, although much less efficiently than AAV4.

### Hematopoietic and immune cell transduction

We also analyzed transduction efficiency of bone marrow-derived hematopoietic stem and progenitor cells, as well as circulating leukocytes. Cells isolated from bone marrow were stained with the appropriate antibodies to identify functional transduction by Cre-induced tdTomato expression by flow cytometry (**Figure 7**, **Supplemental Figure 11, Supplemental Tables 8 and 9)**. Strong transduction of short-and long-term hematopoietic stem cells (ST-HSC and LT-HSC) and multipotent progenitor cells (MPP1, MPP2 and MPP3) was observed for multiple serotypes. For AAVrh8, rh10 and rh74 transduction efficiency was greater than 50% in both males and females (**Figure 7A,B**, **Supplemental Figure 11A)**. Interestingly, AAV7 appeared to transduce hematopoietic stem cells more efficiently in males than in females (66% vs 53%, p=0.03), while for AAV8, females showed a higher transduction efficiency in these cells compared to males (67% vs 48%, p=0.02). Interestingly, AAV9 which exhibited broad biodistribution and functional transduction across multiple organs, was relatively inefficient in bone marrow-derived cells.

**Figure 7.**
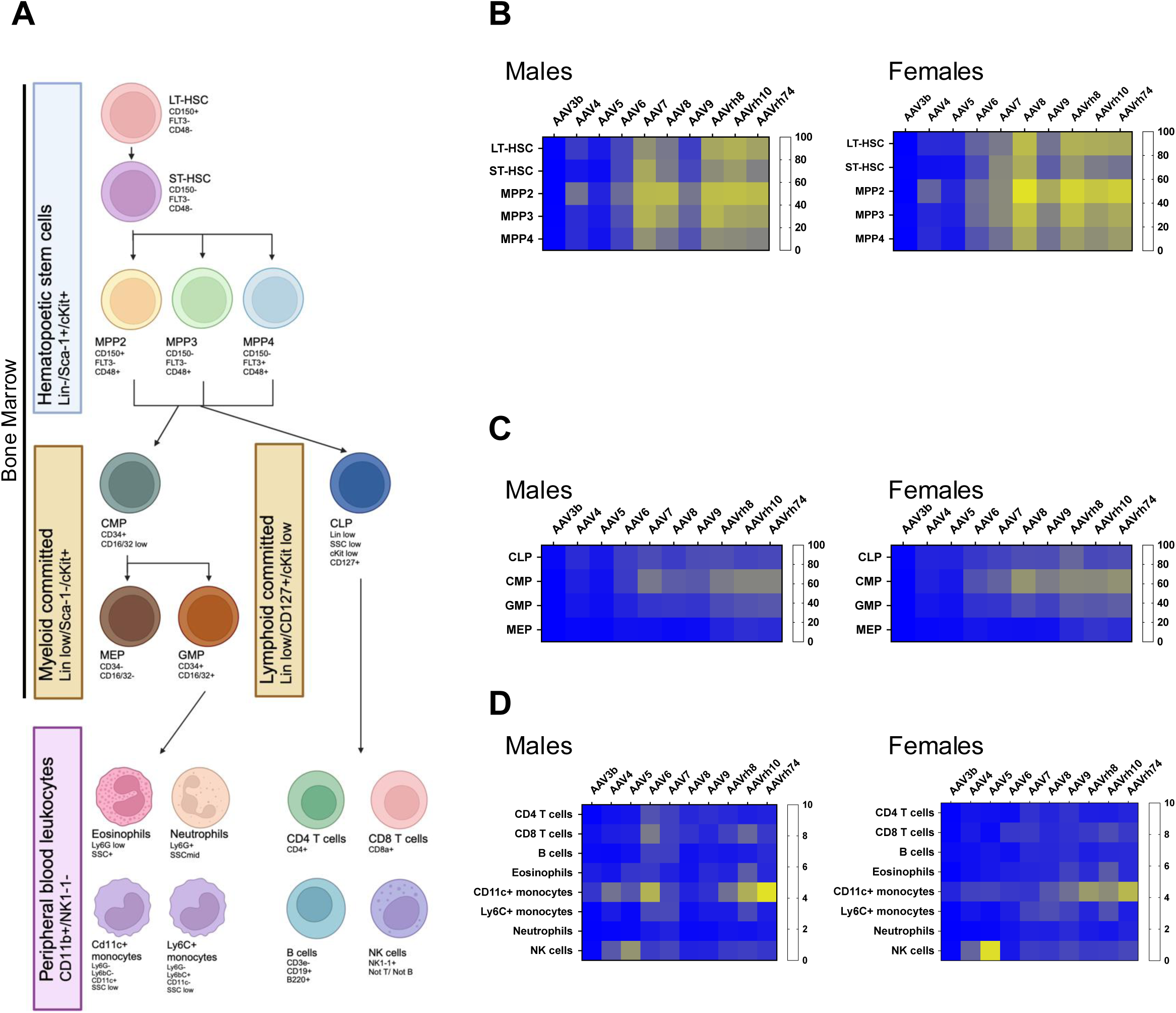
Transduction of hematopoietic stem cells, progenitor cells, and leukocytes. **A)** Hematopoetic stem cell and leukocyte lineages tested. **B)** Transduction of hematopoietic stem cells by AAV - Cells were isolated from the bone marrow of AAV-injected mice and stained for relevant cell types: long-term (LT)-HSC (CD150+, Flt3-, CD48-), short-term (ST)-HSC (CD150-, Flt3-, CD48-), MPP2 (CD150+, Flt3-, CD48+), MPP3 (CD150, Flt3-, CD48+) and MPP4 (CD150-, Flt3+, CD48+). Transduction was scored by activation of tdTomato fluorescence, detected by flow cytometry. Colors on heat maps indicate % tdTomato-positive cells in the population. **C)** Transduction of lineage-committed cells from bone marrow - As above, cells were isolated from bone marrow and stained for relevant cell types: CMP (CD34+, CD16/32 low), MEP (CD34-, CD16/32-), GMP (CD34+, CD16/32+). **D)** Transduction of circulating leukocytes - Peripheral blood mononucleated cells (PBMCs) were collected from AAV-injected mice, and stained for relevant cell types: CD4 T cells (CD4+, CD3e+), CD8 T cells (CD8a+, CD3e+), B cells (CD19+, B220+, CD3e-), eosinophils (Ly6G low, SSC+), CD11c+ monocytes (Ly6G-, Lyb6C-, CD11c+, SSC low), Ly6C+ monocytes (Ly6G-, Lyb6C+, CD11c-, SSC low), neutrophils (Ly6G+, SSC mid) and NK cells (NK1-1+, Not T/NotB).

This analysis was extended to lineage-committed cell types within the bone-marrow. Committed progenitor cells were generally less transduced than stem cells and multipotent progenitors (**Figure 7C**, **Supplemental Figure 11B)**. Common myeloid progenitors were more strongly transduced than common lymphoid progenitors in both males and females by multiple AAV serotypes. However, the more differentiated myeloid progenitor cell types, granulocyte-macrophage progenitors and megakaryocyte-erythrocyte progenitors were less transduced. Circulating leukocytes were also assayed for transduction, following isolation of peripheral blood mononucleated cells. Notably, overall transduction efficiency was significantly lower for these cells compared to bone marrow derived cells. For all cell types and serotypes, mean transduction efficiencies were less than 10% (**Figure 7D**, **Supplemental Figure 11C)**. Nonetheless, some differences were observed. CD11c+ monocytes, which includes dendritic cells, were more highly transduced than other cell types, especially by AAVrh74. In contrast, NK cells showed a stronger transduction by AAV5 than other serotypes. Our results have produced the most complete survey of AAV tropism in the mouse hematopoetic system to date.

## Discussion

Currently, the most cited survey of AAV tropism in mice comes from the work Zincarelli and colleagues (Zincarelli et al., 2008). This study explored the viral biodistribution of nine AAV serotypes (AAV1-9) across eight organs (brain, kidney, heart, liver, lung, skeletal muscle, and testes). Systemic intravenous injection was used because it provides broad tissue distribution and is broadly therapeutically relevant. The AAVs used in the experiments carried luciferase expression cassettes under a ubiquitous promoter, allowing serial measurement of functional transduction through bioluminescent imaging. Their results identified AAV9 as having the widest organ distribution and highest transgene expression. However, this study was limited to male mice, and did not allow examination of transduction at the level of individual cells. More recent surveys have focused on assessing AAV tropism among cell types within a specific organ, including brain (Aschauer et al., 2013; Brown et al., 2021), and eye (Aschauer et al., 2013; Brown et al., 2021), or from a single serotype (Zhao et al., 2020).

Here, we performed a comprehensive evaluation of AAV tropism in mice, the most widely used preclinical model organism for gene therapy. Our experiments expand on prior work in several important ways. We examined ten serotypes originally isolated from humans (AAV3B, AAV5, AAV6 and AAV9) and non-human primates (AAV4, AAV7, AAV8, AAVrh8, AAVrh10 and AAVrh74), with a diversity of capsid structures. Our organ panel consisted of twenty-two separate tissues, as well hematopoietic stem and progenitor cells, and circulating immune cells. A dose of 1E12 GC (10-fold higher than Zincarelli et al.) was chosen to ensure extrahepatic transduction was effectively captured. We also included both male and female mice in our analyses. Perhaps most importantly, we determined functional transduction by fluorescent imaging of individual organ sections. This technique allowed a detailed examination of functional transduction at the level of individual cells, rather than the broad organ-level detection of transgene expression from bioluminescence or beta-gal staining that is traditionally employed. Additionally, by testing serotypes individually we avoided competition for common receptors or variable representation which can confound pooled library approaches.

Results from our evaluation of viral biodistribution correlate well with the previous findings. In general, our values for viral genome copies detected, calculated as genome copies (GC)/μg total genomic DNA, were concordant with previously published values (Zincarelli et al., 2008). As expected, most serotypes (AAV6,7,8,9,rh8,rh10 and rh74) preferentially transduced the liver. In addition, muscle (gastrocnemius and diaphragm), as well as heart, was strongly transduced by these serotypes. Notably, these serotypes are closely related evolutionarily, reinforcing the role of capsid structure in determining viral tropism (Gao et al., 2004). AAV5, which is the phylogenetically most distant from other serotypes, also transduced liver but failed to effectively transduce muscle and heart.

By examining both male and female mice, we were able to uncover evidence of sexual dimorphism with respect to AAV tropism across the natural serotypes tested. As expected, stronger liver transduction was seen in males than females for most serotypes (Davidoff et al., 2003). These sex differences would likely be greater at doses lower than 1E12 GC. The notable exception was AAVrh74, which produced a pronounced preference for transduction in female mice, not only in liver, but across all organs. In contrast, AAVrh10 showed a transduction preference for males for all organs except lymph nodes, which were very weakly transduced in both sexes. AAV6 also demonstrated a bias towards transduction in males across multiple organs. For most other organs and serotypes, sex differences were relatively minor. Nonetheless, sexual dimorphism remains an important consideration for AAV-mediated gene therapy, given the potential impact on efficacy (Piechnik et al., 2022). The extent and nature of sexual dimorphism in human AAV gene therapy remains an open question.

To generate data revealing functional transduction, we employed a dual fluorescent reporter strategy. The transgene contained within our AAVs expressed both the ZsGreen fluorescent reporter and Cre recombinase. In the Ai9 reporter strain used in the studies, Cre expression drives activation of tdTomato fluorescent reporter expression. This event permanently marks the targeted cell and its daughter cells after transduction. During our initial examination of our fluorescent images, we noted that Cre-driven tdTomato expression was visible in significantly more cells than ZsGreen, irrespective of the tissue or serotype. tdTomato activation also had an excellent signal-to-noise ratio allowing for more accurate discrimination of positive cells from background fluorescence. Hence, we found that this technique was the most sensitive mechanism for detecting transduction, corroborating recent findings (Lang et al., 2019).

Our results strongly reinforce the utility of fluorescent imaging for the detection of functional AAV transduction. We detected rare transduction events in individual cells from multiple tissues where the biodistribution data suggested a low tropism for a particular serotype. From a clinical point of view, rare transduction events by a therapeutic AAV may be consequential. Therefore, our results demonstrate the need for careful analysis of transduction of non-target tissues. In particular, we focused on sex organs for evidence of non-target transduction, given concerns over the possible germline transmission of genome alterations.

We noted significant transduction of both testes and ovaries. However, in testes only somatic cells located outside the tubules were transduced. Likewise in ovaries, the transduced cells appeared outside the follicles, also indicating that they were not part of the germ line. Schuettrumpf et al. found that vector genomes are detectable in semen from rabbits receiving an intravenous injection of AAV2 vectors(Schuettrumpf et al., 2006). In follow up work, Favaro et al. analyzed the risk of germline transmission following intravenous delivery of AAV2 and AAV8 vectors expressing Factor IX to rabbits(Favaro et al., 2009). Although recombinant AAV genomes were shed in rabbit semen, these were thought to be derived primarily from epithelial cells from the genitourinary tract. Studies in mice confirm that AAV genomes are shed in semen, but consistently show a lack of transmission to offspring(Ferla et al., 2017; Fonck et al., 2023; Jakob et al., 2005), even when mouse spermatozoa are directly exposed to high doses of AAV2 prior to in vitro fertilization(Couto et al., 2004). The risk of germline transmission could theoretically be greater with AAV-based genome editing approaches. Nonetheless, we did not find evidence of germ cell delivery in either male or female mice using a highly sensitive reporter which permanently marks transduced cells.

Strong physical and functional transduction by AAV6,7,8,9, plus rh8, rh10 and rh74 serotypes was also observed in adrenal glands, an organ not included in previous comparative surveys of AAV tropism, but which has previously been found to be transduced by AAV9 (Zhao et al., 2020). This result validates our decision to include smaller, less frequently examined organs in our survey. Imaging revealed that the transduction occurred primarily in the cortex, rather than the medulla. Adrenals should hence be considered not only as a tissue potentially targetable with AAV delivery, but also one to consider in terms of side effects originating from non-target tissues. Further work is needed to determine to what degree adrenals may also be transduced in non-human primates.

Two serotypes differed in the liver-dominant tropism most frequently observed, with a viral genome copy number 100-fold lower in this organ compared to the other serotypes. Interestingly, these two serotypes AAV3B and AAV4, differed in their tropism in non-hepatic tissues. AAV3B did not appear to transduce other organs in the place of liver. Instead, it did not appear to have an affinity for any specific organ. This effect may be specific for mice, since previous studies have shown that while this serotype has a poor tropism for mouse liver, it can strongly transduce hepatocytes from humans and non-human primates (Wang et al., 2015). The highest physical transduction for AAV3B was observed in lymph nodes and spleen, although fluorescent imaging did not indicate functional transduction of these organs by this serotype. It is possible, given the role of these organs in mediating immune responses, that in these tissues, virus particles are taken up, but processed to generate an immune response, rather than for gene delivery. This biodistribution data provides useful information about potential latent reservoirs of AAV that impact adaptive immunity and long-term efficacy.

A major finding of this work is the pan-endothelial tropism of AAV4, a lesser studied AAV serotype that is difficult to produce at high titer. AAV4 transduced the liver relatively poorly, but strong transduction was observed in the lung as previously reported (Shen et al., 2013; Zincarelli et al., 2008). We found that pulmonary endothelial cells are the primary cell type targeted in the lung with AAV4 by the intravenous route. We also noted substantial and highly specific endothelial cell transduction by AAV4 in many other organs (>10 different tissues), opening the door for endothelial-directed gene therapies. AAV4 differs from other serotypes not only in capsid structure, but also in its unique affinity for α2–3 O-linked sialic acid (Kaludov et al., 2001; Shen et al., 2013). Interestingly, AAV4 is the only serotype we tested which does not require AAVR for cell entry, potentially explaining its poor liver tropism. It is also not clear why AAV4 is so effective for endothelial cells, although other serotypes (i.e. AAV8, AAV9) are presumably transcytosed through the endothelial barrier to enter parenchymal cells. Further work is needed to identify the endocytosis receptor for AAV4, and to define its intracellular trafficking itinerary, which results in endothelial transduction rather than transcytosis. In terms of pancreatic islet delivery, AAV8 has long been known to transduce beta cells following intravenous or intraperitoneal injection in mice (Cheng et al., 2007; Gaddy et al., 2010; Johnson et al., 2013; Rehman et al., 2008; Wang et al., 2006). Previously, AAV8 was used to transduce acinar cells with 3 key beta cell transcription factors, attempting to convert acinar cells to restore depleted beta cells in diabetes (Xiao et al., 2018). However, AAV4 appears to have the greatest selectivity for beta cells over the acinar cells in pancreas with systemic intravenous delivery. Thus, our observations may be useful for the design of delivery vehicles for gene therapies that could directly target expansion of beta cells in diabetes.

There is growing interest in targeting hematopoietic stem and progenitor cells for *in vivo* gene therapy and genome editing. Previous data suggests significant transduction of bone marrow-derived HSCs by AAV6, AAV8 and AAV9 (Goldstein et al., 2019). Our data corroborates and extends these results. AAV based gene editing of such cells within the bone marrow niche would allow for life changing gene therapies. For example, *ex vivo* gene editing therapies have just been approved for sickle cell disease and beta thalassemia. However, the high cost of such therapies combined with the difficult logistics of bone marrow conditioning, means that they are unlikely to be fully translated on a global scale to the populations who would most benefit. A single dose AAV gene therapy that can target the bone marrow following systemic administration would be very powerful. It is also noteworthy, that some immune cell populations were transduced in peripheral blood, primarily CD11c+ monocytes, NK cells, and CD8 T-cells, albeit with much lower efficiency. These cell types are potential targets for cancer immunotherapies (i.e. CAR-T, CAR-NK) as well as gene editing to eradicate residual HIV infection. Effectiveness of such therapies will depend both on the mode of action, as well as efficiency of delivery. This also suggests that unintended transduction of immune cell populations may play an underappreciated role in the safety and efficacy of AAV gene therapies in general.

We believe this resource will be immediately useful for other researchers targeting specific cell types or tissues in mouse models, as well as for avoiding unintended delivery to non-target tissues. These data will inform the community of the most effective vectors for preclinical gene therapy in mice, and more broadly improve our understanding of AAV biology. Moreover, the detailed documentation of targeted tissues and cell types, including the germline, is critical to the ongoing development and deployment of genome editor-based therapeutics. To this end, all data are provided through the Somatic Cell Genome Editing Consortium Toolkit (https://scge.mcw.edu/toolkit) for further exploration and analysis. It remains to be determined how translatable these tropism data are to humans. Although there are many examples where tropism does not translate well across species, many common themes exist. For example, most approved AAV gene therapy products involve AAV2, AAV5, AAV8 or AAV9 capsids. These vectors were tested extensively in mouse, primate, and/or canine models before progressing to human clinical trials. Vectors capable of transducing liver, skeletal muscle, CNS, and retina, are generally effective across multiple species-albeit with varying efficiency. An important endeavor will be comparing tropism data from mice reported here, with that obtained in non-human primates, as well as with clinical outcomes. Ideally, cross-species compatible vectors can be developed for specific tissues using these data as a starting point, thus streamlining the development of gene therapies from initial proof-of-concept in mice to ultimate application in humans.

## Materials and Methods

### Recombinant AAV Production

AAV vectors were produced by standard triple-transfection in HEK293 cells. Three days post transfection, cells and culture media were harvested, and crude lysates were prepared. Downstream purification was carried out by two rounds of CsCl sedimentation followed by dialysis. Purified AAV vectors were sterilized using a 0.22-micron filter and formulated in phosphate buffered saline (PBS) with 5% sorbitol and 0.001% of F-68. AAV vector genome titers were determined by droplet digital PCR. AAV vector purity was assessed by gel electrophoresis followed by silver staining.

### AAV Transduction in mice

Mice used in experiments were from the Ai9 strain, B6.Cg-*Gt(ROSA)26Sor^tm9(CAG-tdTomato)Hze^*/J (Madisen et al. 2010, JAX# 007909). At five weeks of age, male and female mice were injected intravenously through the tail vein with recombinant AAV. Injections were carried out in parallel at BCM and JAX using the same viral batch. Four to six mice of each sex were injected with AAVs of each serotype. Dosage was 1E12 GC/mouse, in a total volume of 0.2 ml phosphate-buffered saline. Following injection, mice were housed under standard conditions, and fed *ad libitum*, while monitoring for weight loss or other health issues. After four weeks, the mice were humanely euthanized, and dissected for biodistribution and functional transduction analyses. In experiments involving labeling of endothelial cells, 100 mg of DyLight649-labelled *Lycopersicon Esculentum* (tomato) lectin (Vector Laboratories, DL-1178-1) was injected intravenously through the tail vein in a volume of 0.1 ml just prior to euthanasia and dissection. Tissue samples from each organ were prepared for imaging as described below. In addition, small samples (∼50 mg) were flash frozen in liquid nitrogen and stored at -80°C for subsequent genomic DNA preparation. For brain samples, one hemisphere cut sagittally was used for DNA preparation, to account for possible differences in AAV transduction between regions. All animal work was performed in accordance with the guidelines for animal care at Baylor College of Medicine (BCM) and the Jackson Laboratory (Jax), under approved Institutional Animal Care and Use Committee protocols.

### DNA Isolation and Quantitative PCR

Total DNA (viral and mouse genomic) isolation from frozen samples was performed using DNeasy Blood & Tissue kits (Qiagen) according to the manufacturer’s instructions. Tissue disruption steps were repeated twice on muscle and heart samples to ensure quantitative recovery of DNA. The DNA concentration in each sample was assayed by fluorescence using a Quant-iT Broad Range kit (Invitrogen). Absolute viral genome copy numbers in each sample were determined by SYBR-based quantitative PCR using a standard curve with the AAV transgene plasmid and validated primers specific to the Cre recombinase transgene (Forward: 5ʹ-TGA CGG TGG GAG AAT GTT AAT C-3ʹ, Reverse: 5ʹ-GCT ACA CCA GAG ACG GAA ATC-3ʹ) (Lang et al., 2019). Results were calculated as AAV genome copies per μg input total DNA.

### Fluorescent Imaging

Tissue preparation and imaging at JAX were carried out as follows: Twelve mice per AAV serotype treatment group were euthanized via CO_2_ asphyxiation. For collection of blood cells and bone marrow for flow cytometry analysis, immediately following confirmation of expiration, peripheral blood (250-500 uL) was collected via cardiac puncture into K2-EDTA microtainer tubes (BD #365974) and leg bones (either 2 femurs or 1 femur and 1 tibia) were collected into cold 1XPBS i 12 well plates for isolation of bone marrow. The remaining tissues: brain (sagittal bisection), eyes, inguinal lymph node, skin, bladder, ovary, uterus, epididymis, testes, large intestine, gastrocnemius muscle, diaphragm, heart, thymus, lung, spleen, kidney, liver, adrenal gland, pancreas, small intestine, and stomach were collected into 7 vials and immersion fixed in 25ml, cold 4% PFA in 1xPBS per vial at 4°C on a shaking incubator for 4-5 hours. Tissues were washed in cold 1X PBS twice and then transferred to into cold 30% sucrose/1XPBS solution on a shaking incubator overnight to equilibrate. Excess connective tissue was trimmed and tissues were combined into 7 groups/blocks to minimize the number of blocks per mouse, and then immersed in Tissue-Tek O.C.T in cryomolds, frozen on dry ice, and stored at 80°C. Tissues were cryosectioned at 15μm and sections mounted on Superfrost Plus slides (Fisher Scientific), allowed to air dry for ∼10 min at room temp, and washed with 1xPBS twice for 5 minutes. Slides were then stained with 1:10,000 Hoechst in 1xPBS for 5 min, washed 2x with 1xPBS and slides mounted using ProLong Diamond Antifade Mountant, cured overnight, sealed with clear nail polish. Tissues were imaged using a widefield DMi8 microscope (Leica) using a 10X or 20X objective. Representative sections from each tissue were analyzed and at least 3 animals per sex per treatment were assessed, with most including 4 mice per sex per treatment. Quantification of tdTomato (RFP) signal and thus viral transduction was performed as follows: images were opened in QuPath (Bankhead et al., 2017), and cells were detected using the built-in watershed cell detection function based on DAPI/nuclear signal. All images from the same tissue type were processed with the same cell detection settings, with only slight changes in parameters used for different tissues to account for changing cell and nuclear diameters (i.e. adjusting sigma microns (range 0.5-0.9) and cell expansion microns). Background signal for RFP expression was determined empirically from uninjected control tissues from Ai9 mice imaged at defined increasing exposure times and used to set detection threshold minimums for each image. RFP signal above background threshold was scored per cell (co-localization). Data recorded per image includes the total number of DAPI nuclei (cells) and the number of nuclei which had RFP signal above threshold (transduced cells) and we report the calculated % of transduced RFP+ cells out of the total number of cells per image (RFP+ cells/total cells x 100%).

Tissue preparation and imaging at BCM were carried out as follows: Following euthanasia, animals were perfused briefly with phosphate-buffered saline through a cardiac puncture, then organs were dissected. Tissue samples from each organ were placed in 8 ml of freshly prepared 4% buffered paraformaldehyde and fixed overnight at 4°C with mild shaking. The next day, fixed samples were transferred to a 15% buffered sucrose solution, then three hours later, transferred to a 30% buffered sucrose solution for equilibration overnight at 4°C. Next, the fixed and cryopreserved samples were frozen in OCT, sectioned at 14 μm, DAPI-stained for nuclei, and imaged on an Axioscan Z.1 slide scanner (Zeiss) under a 20X objective, with the appropriate fluorescence settings, at the RNA *In Situ* Hybridization Core at Baylor College of Medicine. Sectioning was performed to maximize the imaged tissue surface area for each organ. Three non-consecutive sections from each sample were imaged. For experiments involving lectin labeling of the vasculature, 40 μm sections were prepared to generate a wider depth of field.

Aortas were prepared for imaging by cutting the left and right carotid arteries and the left subclavian artery to release the aortic arch. Then, the aorta was cut at the level of the diaphragm. The heart and aortic arch were removed, and the remaining thoracic aorta was *en face* opened and pinned on a dissecting dish with the luminal side facing up. The thoracic aorta was fixed overnight with 4% PFA (in PBS), then incubated overnight with 30% Sucrose (in PBS) and stored in PBS at 4°C afterwards. A ∼4 mm piece of aorta was cut and stained with DAPI (1:5000 dilution in PBS, D3571, Invitrogen) for 5 minutes, followed by a wash in PBS for 3 minutes. Then, the sample was placed flat on a glass slide with the lumen faced up and covered with a coverslip with glycerol. Direct tdTomato, DyLight649-lectin and DAPI fluorescence were detected using a Nikon A1-Rs confocal microscope using a 60x objective at the Integrated Microscopy Core (Baylor College of Medicine). Confocal sections were acquired every 2 µm from the lumenal side to the first layer of smooth muscle cells.

For immunofluorescence images of pancreatic islets, PFA-fixed and frozen pancreas sections were brought to room temperature for 5 minutes and then placed 0.1% Triton-X-100 in PBS for 5 minutes. A one-hour block was performed with 5% donkey serum in PBS, followed by a two-hour incubation with primary antibodies to insulin (1:1500; rabbit; Cell Signaling #3014s; RRID:AB_2126503. Slides were washed twice in 0.01% Triton-X-100 in PBS for 5 minutes each. Slides were then incubated with secondary antibodies for donkey anti-rabbit (1:1000; Jackson ImmunoResearch #711-175-152, RRID:AB_2340607) and DAPI (1:2000; Sigma #32670) followed by two washes in 0.01% Triton-X-100 in PBS for 5 minutes each and coverslip mounting with Prolong Gold Antifade (Invitrogen #P36930). Omission of the primary antibody was used a control. Images were captured on a Nikon A1R confocal microscope.

### Flow cytometry

To assess pulmonary endothelial cell transduction, Ai9 mice were injected intravenously through the tail vein with 5E+11 GC/mouse of AAV4-CB6-PI-Cre. Mice were humanely euthanized two weeks post-injection, and lungs were excised and processed for flow cytometry using a mouse lung dissociation kit according to the manufacturer’s instructions (Miltenyi Biotec). The resulting single cell suspension was resuspended in PBS/0.5%BSA/2mM EDTA, and non-specific antibody binding was blocked with anti-mouse CD16/CD32 (BD Pharmingen #553142) at 1:100 dilution for 15 minutes at 4 °C. Cells were stained with fixable Live/Dead Blue to exclude dead cells, and antibodies specific for the following markers: CD31 (BV605, clone 390, BioLegend #102427), CD45.2 (FITC, clone 104, BioLegend #109806), CD11b (PerCP-eFluor710, clone M1/70, Invitrogen #46-0112-80) at a 1:100 dilution for 45 min at 4 °C. Flow cytometry was performed on an LSRII (BD Biosciences) instrument, and data analyzed using FloJo software (v10, BD Biosciences).

To assess cell transduction in hematopoietic lineages, whole blood and bone marrow were collected from Ai9 mice injected with 1E+12 GC/mouse. Leg bones were kept cold in 1XPBS in 12 well plates on ice. Bones were crushed with mortar and pestle and rinsed with AutoMACS buffer (Miltenyi Cat# 130-091-222) with BSA (Miltenyi Cat#130-091-376) (final concentration of 0.5% BSA). Liquid was filtered through Falcon 32235 5ml tubes with cell strainer. Cells were centrifuged at 500xg for 5 minutes and liquid decanted. Cells were resuspended at approximately 2x10^7 per ml and 1x10^6 cells were stained with either antibody cocktail BM-A or BM-B for 30 minutes at 4 degrees C. Samples were washed with PBS with 5mM EDTA and 0.5% BSA, stained with 10ul of 5ug/ml DAPI (BD Biosciences, Cat.#564907) per 200ul of sampleand analyzed on BD FACSymphony A5 with 500K live cells set as stopping gate. Whole blood for white cells (peripheral blood lymphocytes-PBL) was treated with 3ml 1X Leinco RBC Lysis buffer for 5 minutes at RT, centrifuged for 5 min at 500xg, decanted, resuspended with 3ml 1X RBC Lysis Buffer for 5 minutes at RT, centrifuged for 5 min at 500xg, decanted, washed with 4ml PBS with 5mM EDTA and 0.5% BSA, centrifuged for 5 min at 500xg, decanted, blotted on paper towel, and resuspended in a total volume of 50ul for staining. 10ul of PBL staining cocktail was added for 30 minutes at 4°C, washed with 2ml of PBS with 5mM EDTA and 0.5% BSA, centrifuged for 5 min at 500xg, decanted, resuspended in 200ul of PBS with 5mM EDTA and 0.5% BSA, stained with 10ul of 5ug/ml DAPI, and run on the cytometer. Samples were analyzed on BD FACSymphony A5 and data analyzed using FloJo (v10, BD Biosciences).

Antibody staining panels for the various hematopoietic lineages are listed below: hematopoietic stem cells (HSC, BM-A), Committed Lymphoid and Myeloid Progenitors (CLP/CMP, BM-B), and Peripheral Blood Lymphocytes (PBL). Antibody, fluorophore, clone, supplier, catalog number, volume used in staining cocktail mix (μL) for each panel.

| Staining Panel | Cell Surface Marker | Fluorophore | Clone | Supplier | Catalog number | volume (ul) |
| --- | --- | --- | --- | --- | --- | --- |
| BM-A (HSC) | CD150 | PE-Cy7 | Clone TC15-12F12.2 | Biolegend | 115904 | 1.5 |
|  | c-kit | APC | Clone 2B8 | BD Biosciences | 553356 | 2 |
|  | Flt3 | BV421 | Clone A2F10.1 | BD Biosciences | 562898 | 15 |
|  | Sca-1 | BV510 | Clone D7 | BD Biosciences | 565507 | 4 |
|  | CD48 | BV786 | Clone HM48.1 | Biolegend | 103449 | 15 |
|  | Fc Block (anti-CD16/CD32) | ~ | Clone 2.4G2 | Leinco Technologies | C247 | 1 |
|  | Lineage Biotin mix | ~ |  |  |  | 16.5 |
| **BM-B (CLP/CMP)** | CD127 | PE-Cy7 | Clone A7R34 | Biolegend | 135014 | 7.5 |
|  | CD34 | AlexaFluor 647 | Clone RAM34 | BD Biosciences | 560230 | 15 |
|  | CD16/CD32 | AlexaFluor 700 | Clone 93 | eBioscience | 56-0161-82 | 7.5 |
|  | c-kit | BV421 | Clone 2B8 | BD Biosciences | 562609 | 2 |
|  | Sca-1 | BV510 | Clone D7 | BD Biosciences | 565507 | 4 |
|  | Lineage Biotin mix | ~ |  |  |  | 16.5 |
| **PBL** | CD19 | Brilliant Blue 700 | Clone 1D3 | BD Biosciences | 566411 | 3 |
|  | CD11c | PE-Cy7 | Clone N418 | TONBO Biosciences | 60-0114-U100 | 3 |
|  | CD3e | APC | Clone 145-2C11 | Biolegend | 100312 | 7.5 |
|  | CD8a | APC-R700 | Clone 53-6.7 | BD Biosciences | 564983 | 1.5 |
|  | Ly6G | BV421 | Clone 1A8 | BD Biosciences | 554063562737 | 3 |
|  | Ly6C | BV570 | Clone HK1.4 | Biolegend | 128029 | 3 |
|  | NK1.1 | BV711 | Clone PK136 | Biolegend | 108745 | 7.5 |
|  | B220 | BUV496 | Clone RA3-6B2 | BD Biosciences | 564662 | 3.75 |
|  | CD11b | BUV661 | Clone M1/70 | BD Biosciences | 612977 | 1.5 |
|  | CD4 | BUV805 | Clone GK1.5 | BD Biosciences | 564922 | 3 |
|  | Fc Block (anti-CD16/CD32) | ~ | Clone 2.4G2 | Leinco Technologies | C247 | 1 |
| **Lineage Biotin mix:** | CD3e | biotin | Clone 145-2C11 | TONBO Biosciences | 30-0031-U500 |  |
|  | CD4 | biotin | Clone GK1.5 | Biolegend | 100404 |  |
|  | CD8 | biotin | Clone 53-6.7 | eBioscience | 13-0081-86 |  |
|  | B220 | biotin | Clone RA3-6B2 | BD Biosciences | 553086 |  |
|  | Gr-1 | biotin | Clone RB6-8C5 | eBioscience | 13-5931-86 |  |
|  | CD19 | biotin | Clone 1D3 | BD Biosciences | 553784 |  |
|  | NK1.1 | biotin | Clone PK136 | eBioscience | 13-5941-85 |  |
|  | Ter119 | biotin | Clone TER119 | Biolegend | 116204 |  |
|  | CD11b | biotin | Clone M1/70 | eBioscience | 13-0112-86 |  |

### Statistical analysis

All possible pairwise comparisons (k=45) of AAV serotypes, within sex and organ (cell type), for each type of data (biodistribution, function transduction, functional transduction by flow cytometry) were performed using the pairwise.wilcox.test() function in package: stats of R version 4.3.2. P-values were calculated by the exact method unless there were ties, in which case, the function used the Chi-square approximation with continuity correction. Within each organ and sex, p-values were adjusted using the false discovery method (Benjamini and Hochberg, 2018).

## Data Availability

All data and images associated with these studies are publicly available at https://scge.mcw.edu/toolkit, as part of the Somatic Cell Genome Editing initiative.

## Acknowledgments

This work was made possible by NIH grants U42 OD026645 to J.D.H., M.E.D., and W.R.L.; U42 OD035581 to J.D.H., W.R.L. and C.J.W; U42 OD026635 to S.A.M. and C.M.L.; and UG3 HL147367 to G.G. Additional support came from grants UG3 HL151545, R01 HL132840, and R01 DK124477 to W.R.L., grants R01DK136694 and R56DK128098 to A.R.C., and grant K01DK128226 to M.A.-B.. The Jackson Laboratory Imaging Sciences and Flow Cytometry Cores receive support from the JAX Cancer Center (P30CA034196). The BCM RNA In Situ Hybridization Core supported these studies and receives funding from the Baylor College of Medicine Intellectual and Developmental Disabilities Center (IDDRC) (P50HD103555). This project was also supported by the Cytometry and Cell Sorting Core at Baylor College of Medicine with funding from the CPRIT Core Facility Support Award (CPRIT-RP180672), the NIH (CA125123 and RR024574) and the assistance of Joel M. Sederstrom. Additional support was provided by the Quantitative Science Shared Resource at BCM (P30CA125123). We also thank Kazuhiro Oka and Austin Seal at the BCM Gene Vector Core for assistance with packaging AAV4-Cre vectors.

## Author Contributions

The studies were conceived by W.R.L., J.D.H., C.J.W., D.W., G.G., C.M.L., K.J.S., and S.A.M. in consultation with O.M. and other members of the NIH Somatic Cell Genome Editing Consortium, and NIH program officials. Funding to support the work was provided by grants awarded to C.J.W., S.M.H., M.E.D., C.M.L., M.A.-B., J.D.H., S.A.M., G.G. and W.R.L. AAV vector design and production was carried out by J.B., D.W. and G.G. Studies of biodistribution of AAV genomes were conducted by C.J.W., A.E.M., M.D.G. and M.A.C. Studies of functional transduction by fluorescence were carried out by K.J.S., E.S., J.D., Y.G., S.H., S.A.M and C.M.L. Additional sectioning and imaging was performed by M.C.L. Analysis of endothelial cell transduction was carried out by C.J.W., M.D.G., M.A.-B., C.F.S. and M.C.L.. Analysis of pancreatic islet transduction was performed by A.R.C. and M.C.L. with support from S.M.H. Expertise on confocal microscopy imaging provided by C.W.H. Generation of experimental reporter mice was achieved by D.G.L. and J.D.H. Statistical analyses of transduction data was performed by C.J.W. and S.G.H. The manuscript was written by C.J.W., W.R.L., J.D.H., K.J.S., S.A.M., D.W., and G.G. with input and approval from all authors. The content is that of the authors and is not intended to represent the official views of the National Institutes of Health.

## Supplemental Information

Supplemental Tables S1-S9

Supplemental Figures S1-S11

Document S1: Supplemental Figure Legends

