## Supplemental Figures and Tables for "A Comprehensive Atlas of AAV Tropism in the Mouse"

#### Supplementary Figure Legends

##### Supplementary Figure 1

**Viral biodistribution analysis – Serotype comparison**: qPCR results are plotted to show all serotypes assayed from the specified organ. Results from male (blue) and female (pink) mice are shown with somatic organs listed in alphabetical order, and reproductive organs at the end. Error bars represent +/- SEM.

##### Supplementary Figure 2

**Sex differences in biodistribution.** Viral genome copy number data from Figure 2 was compared between sexes and mapped as relative differences (male/female). Blue indicates a bias in transduction efficiency towards males, while pink indicates a bias towards females.

##### Supplementary Figure 3

(a) **Comparison of zsGreen and tdTomato fluorescence - AAV4:** A panel of organs from AAV4-treated mice was imaged for zsGreen and tdTomato. In each case, tdTomato produced a a stronger signal, indicating the superiority of Cre-activated tdTomato as a maker of viral transduction. White bars indicate 100 μm.

(b) **Comparison of zsGreen and tdTomato fluorescence - AAV9:** As in (a), but with AAV9. Again, tdTomato proved to be a superior marker of viral transduction.

##### Supplementary Figure 4

**Functional transduction detected by fluorescent imaging:** Following dissection, organ samples were prepared and imaged for fluorescent tdTomato (red). Nuclei were stained for DAPI (blue). Sample images from all serotypes and organs are shown.

##### Supplementary Figure 5

**Functional transduction by individual organ:** The data described in Figure 5 is plotted by organ, serotype, and sex (blue bars for males, pink bars for females). Error bars represent +/- SEM.

##### Supplementary Figure 6

**Correlation between viral biodistribution and functional transduction - combined:** Average viral genome copy numbers are plotted versus corresponding tdTomato-positive cell values from image quantification for each serotype. Males and females are plotted separately.

**Supplementary Figure 7**

**Correlation between viral biodistribution and functional transduction by individual organ:** Average viral genome copy numbers are plotted versus corresponding tdTomato-positive cell values from image quantification for each serotype. Males and females are plotted separately. Spearman's rank correlation coefficients (r) and corresponding p-values are shown.

**Supplementary Figure 8**

**Transduction of endothelial cells in a panel of tissues:** As in Figure 7c, tissue sections from negative control and AAV4-Cre injected mice are shown, with DAPI staining for nuclei (blue), DyLight649-lectin staining for endothelial cells (yellow) and tdTomato (red). Images show coincident tdTomato expression in lectin-stained endothelial cells.

**Supplementary Figure 9**

(a) **Transduction of endothelial cells in retina:** As above, retinas were dissected and imaged for nuclei (DAPI, blue), DyLight649-lectin (yellow) and tdTomato (red). tdTomato-positive cells were coincident with lectin-stained endothelial cells.
(b) **Transduction of endothelial cells in aorta:** As in Figure 7d, sections from the aortic root and aortic arch were imaged for nuclei (DAPI, blue), DyLight649-lectin (yellow) and tdTomato (red). tdTomato-positive cells, indicating AAV4-Cre transduction, were observed on the surface of the aorta, coincident with lectin-stained endothelial cells.

**Supplementary Figure 10**

**Transduction of pancreatic islets by multiple serotypes:** Pancreas sections from AAV injected animals from all ten serotypes were examined for tdTomato expression in apparent islets. As expected, AAV4 produced a strong tdTomato signal restricted to an islet. AAV5 also transduced an islet, but to a lower extent than AA4. AAV8 transduced multiple cell types in the pancreas including islets. Other serotypes were much less effective at transducing islets.

**Supplementary Figure 11**

(a) **Transduction of hematopoietic stem cells by AAV:** Flow cytometry data described in Figure 9a is plotted by cell type, serotype, and sex (blue bars for males, pink bars for females). Error bars represent +/- SEM.

(b) **Transduction of lineage-committed cells from bone marrow:** Flow cytometry data described in Figure 9b is plotted by cell type, serotype, and sex. Error bars represent +/- SEM.

(c) **Transduction of circulating leukocytes:** Flow cytometry data described in Figure 9c is plotted by cell type, serotype, and sex. Error bars represent +/- SEM.

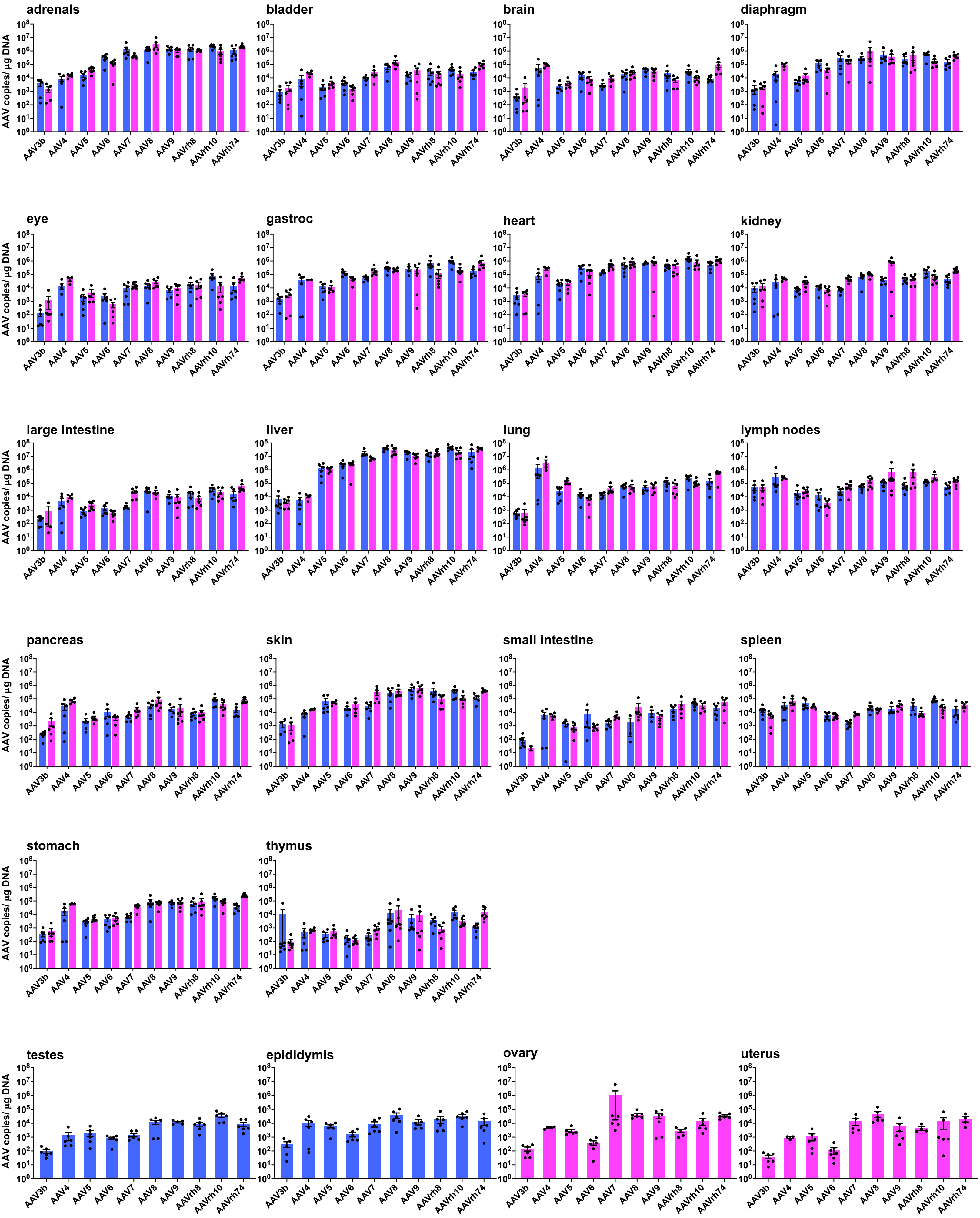


Supplementary Figure 1 - Biodistribution by individual organs

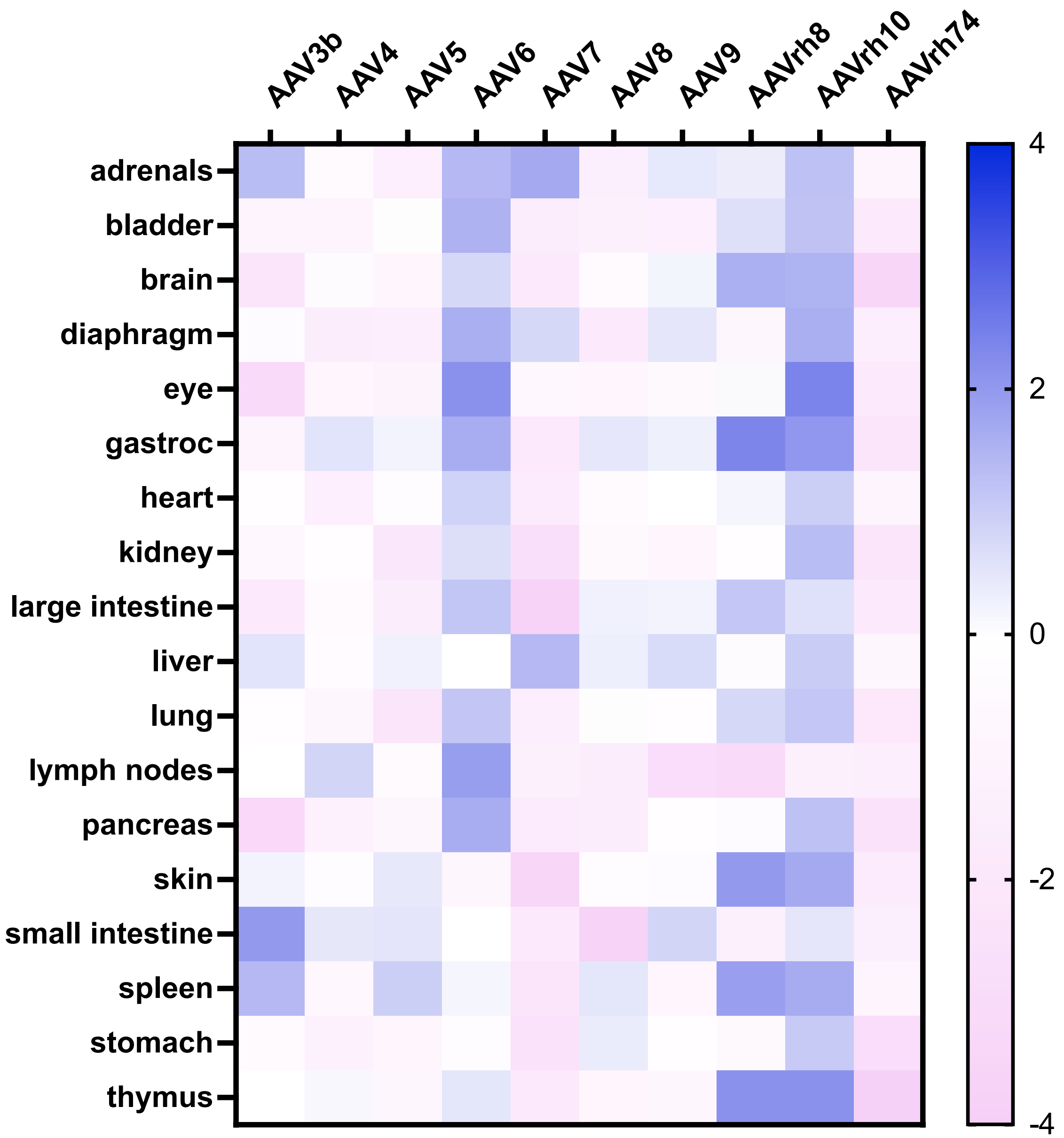


Supplementary Figure 2 – Sex differences

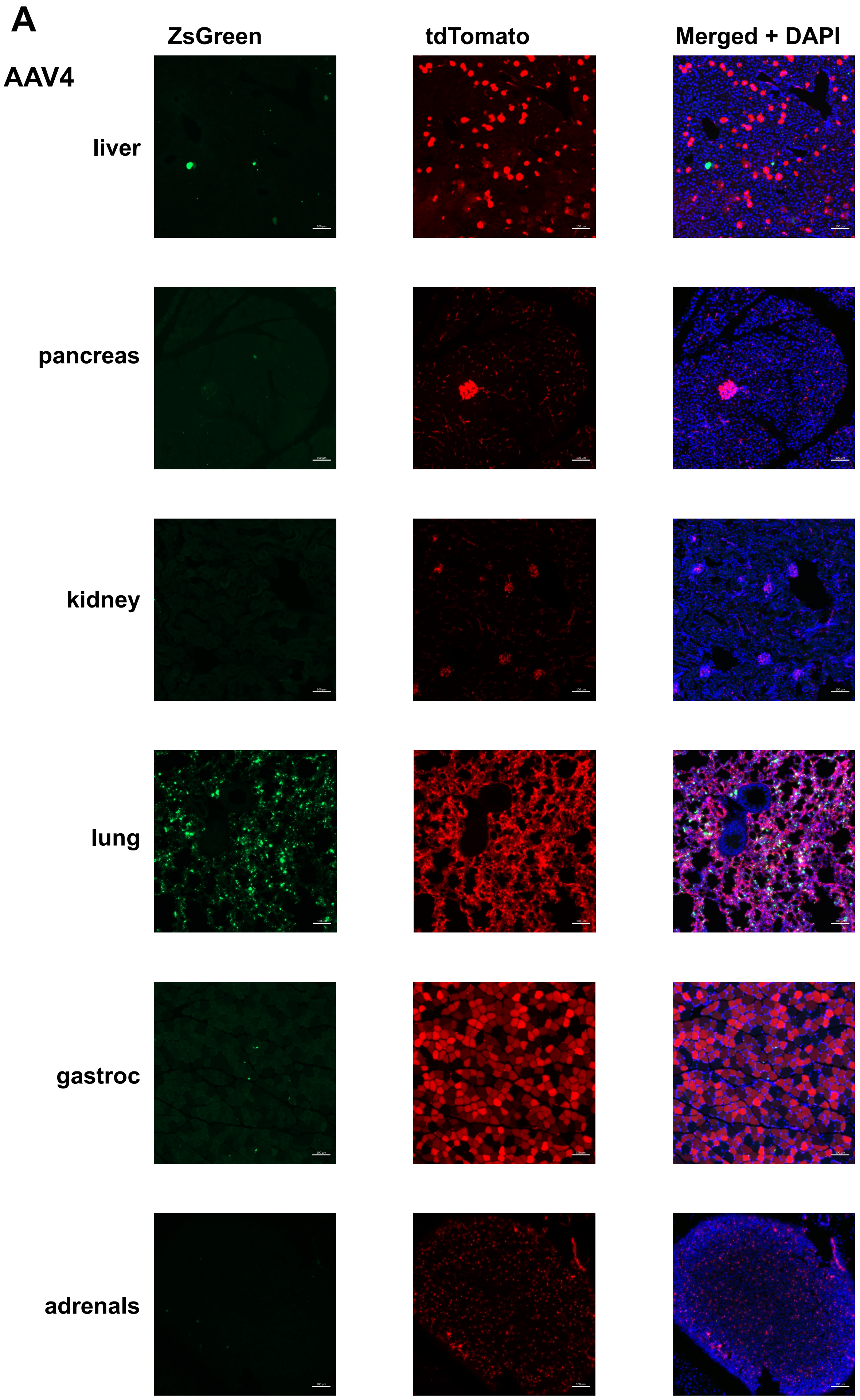


Supplementary Figure 3 - ZsGreen vs. tdTomato

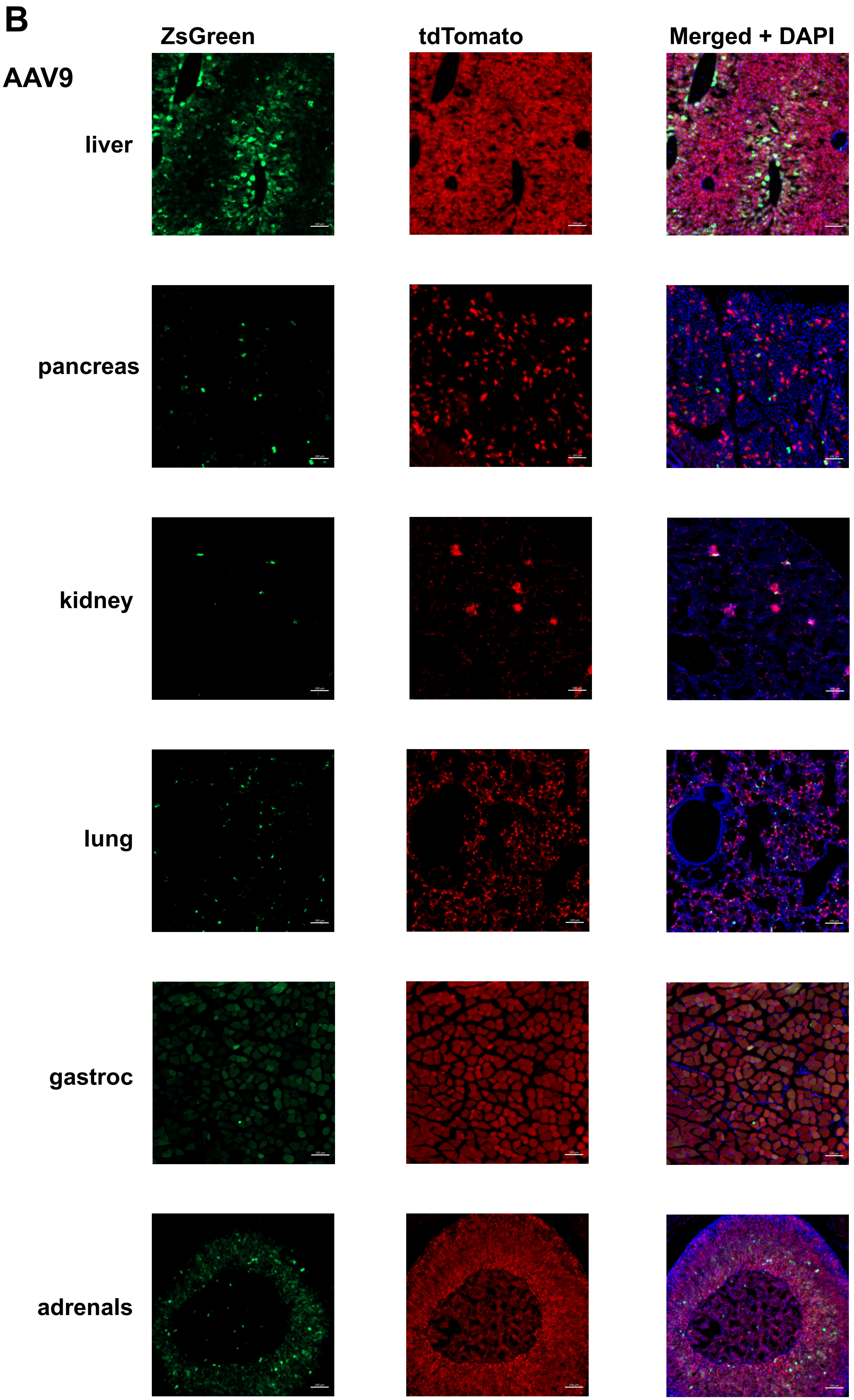


Supplementary Figure 3 - ZsGreen vs. tdTomato

### Supplementary Figure 4 - Images from all organs and serotypes

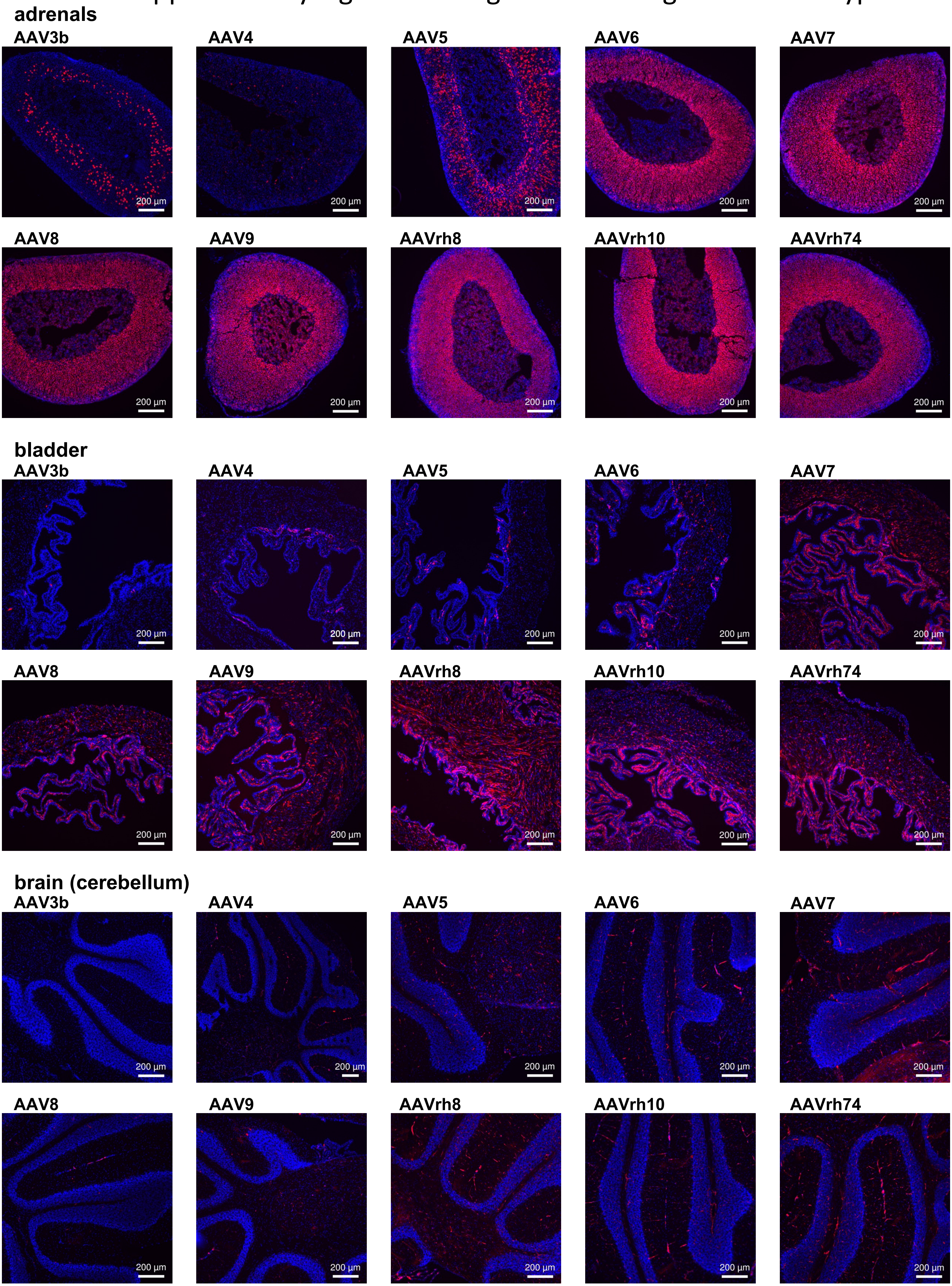

Supplementary Figure 4 - Images from all organs and serotypes

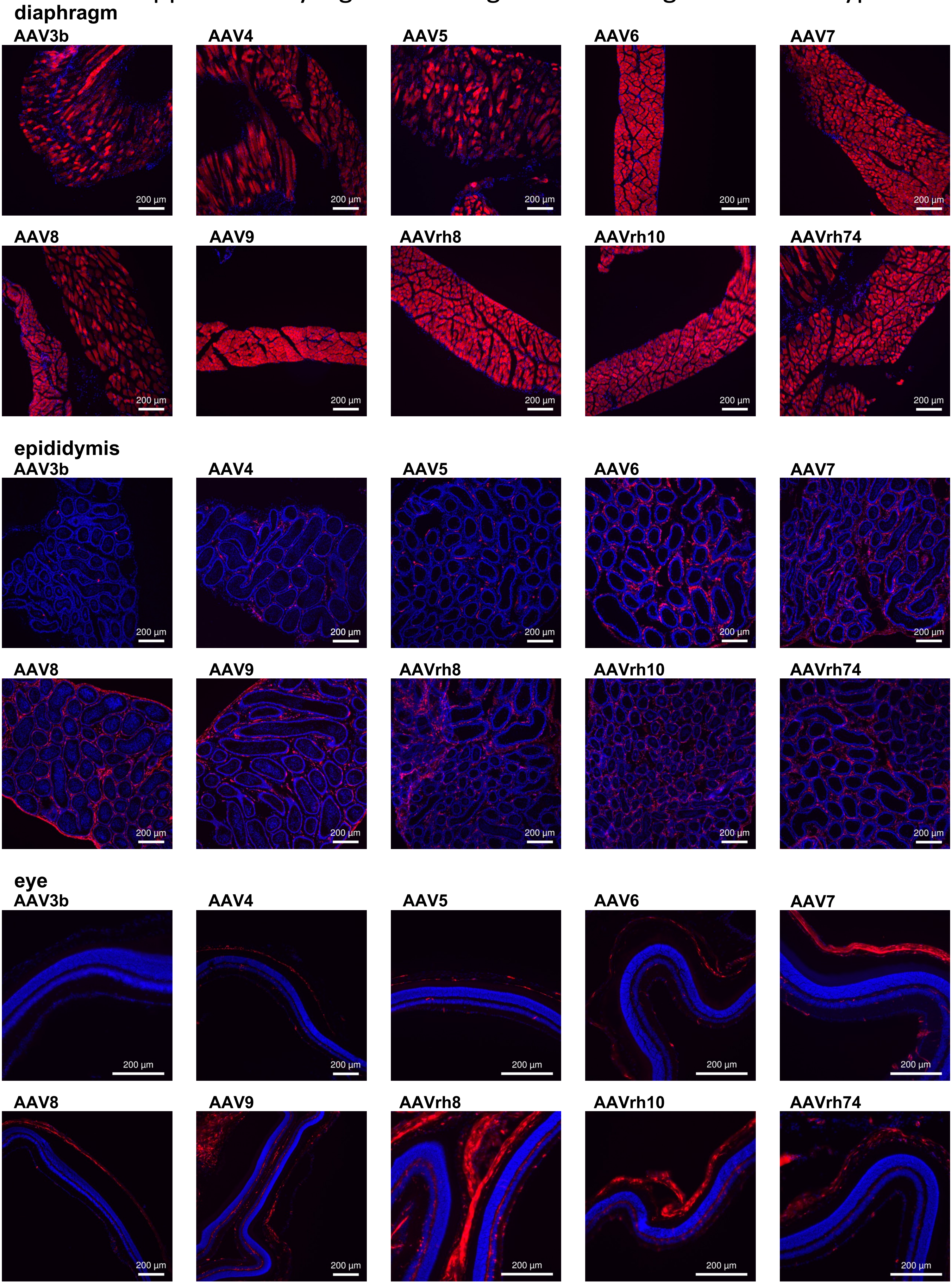

Supplementary Figure 4 - Images from all organs and serotypes

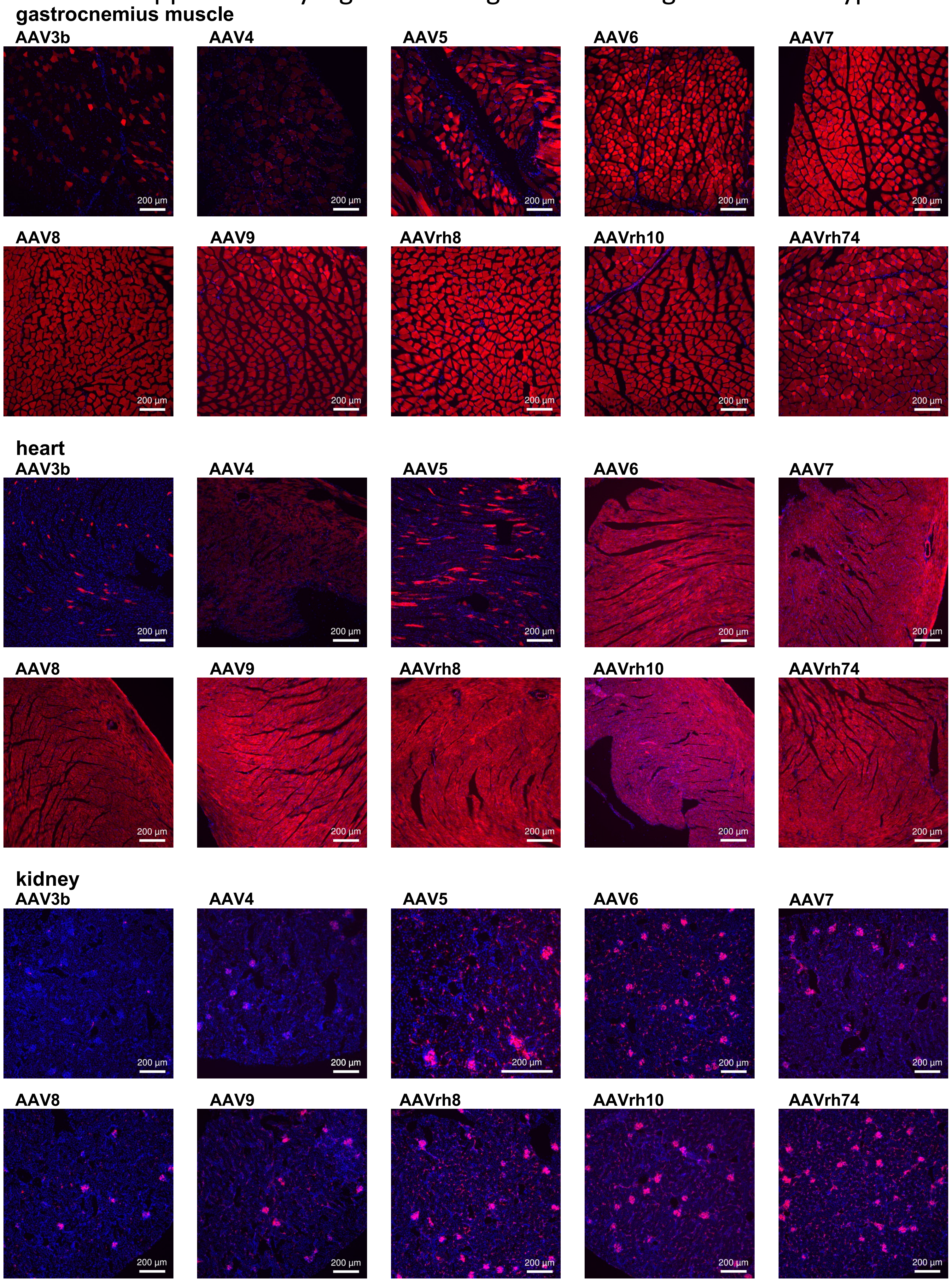

### Supplementary Figure 4 - Images from all organs and serotypes

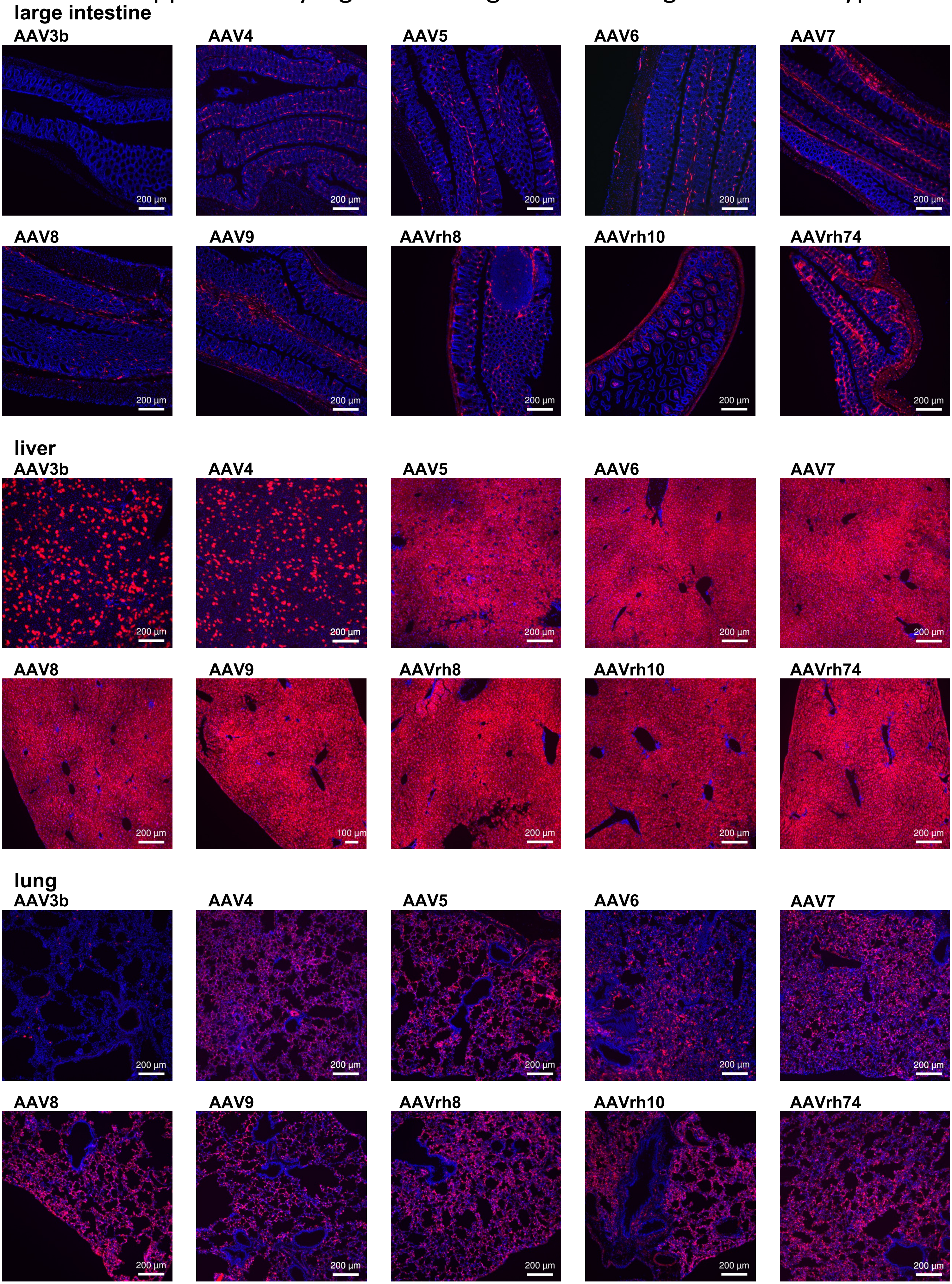

### Supplemntary Figure 4 - Images from all organs and serotypes

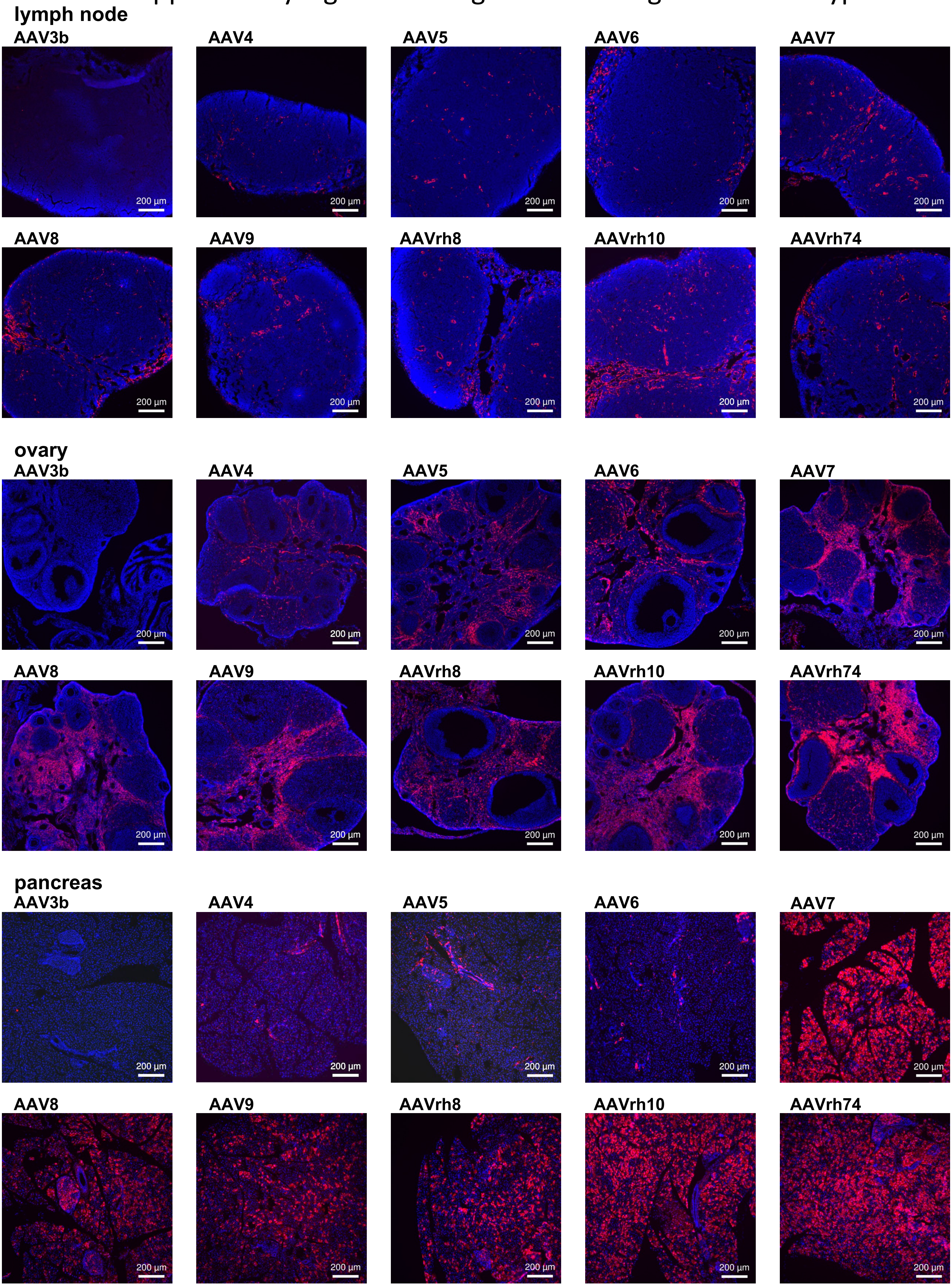

Supplementary Figure 4 - Images from all organs and serotypes

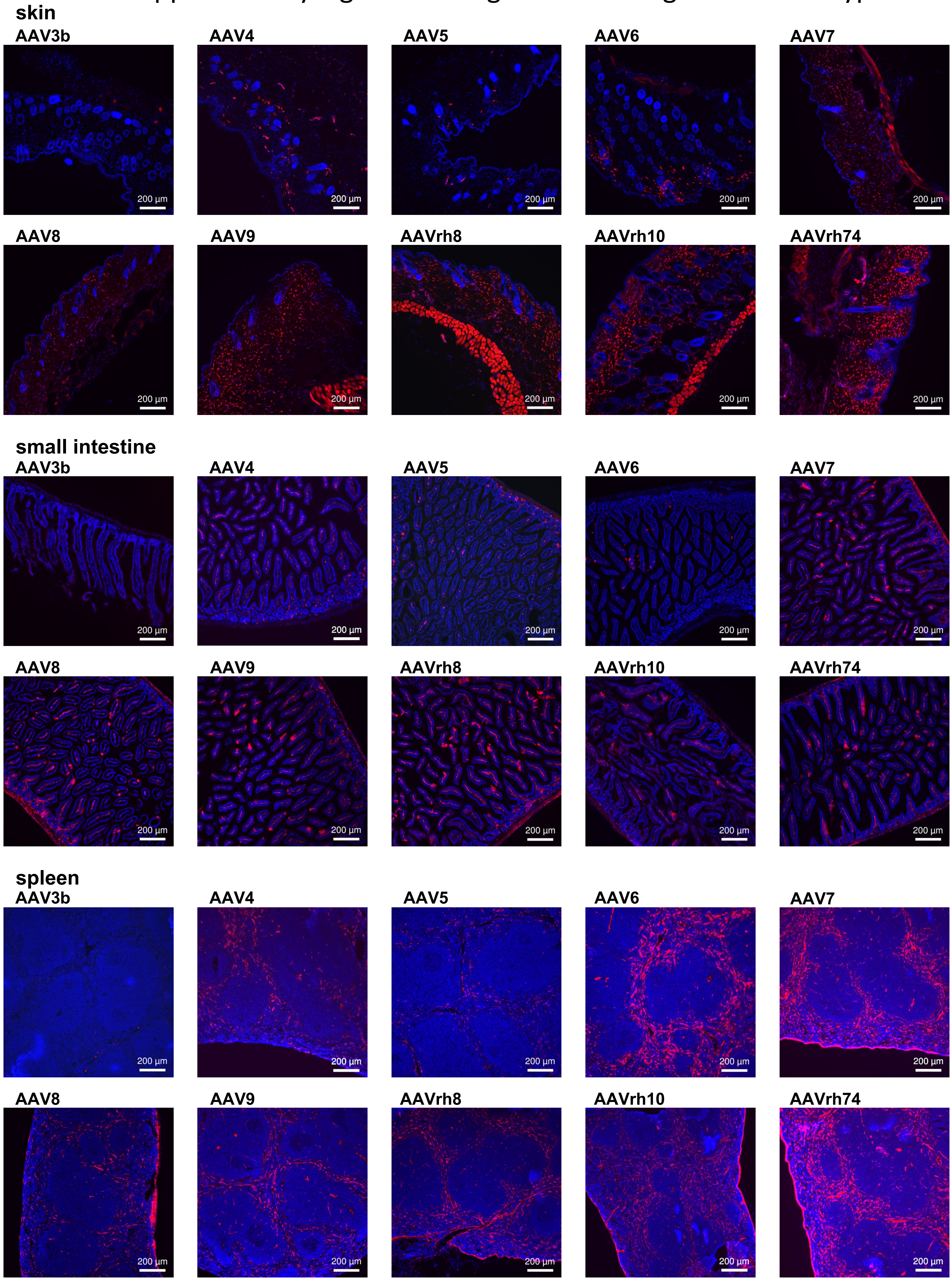

### Supplementary Figure 4 - Images from all organs and serotypes

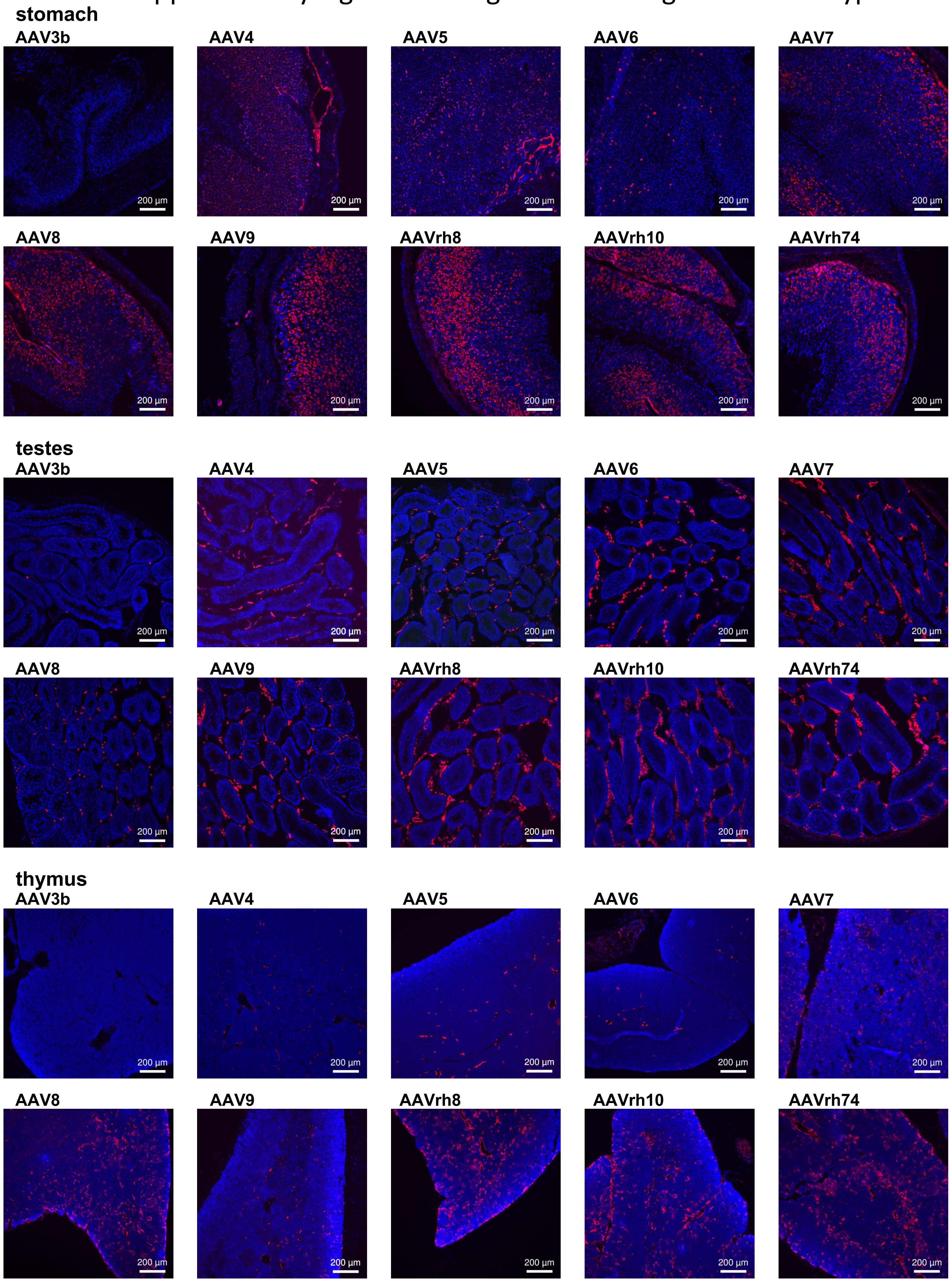

### Supplementary Figure 4 - Images from all organs and serotypes

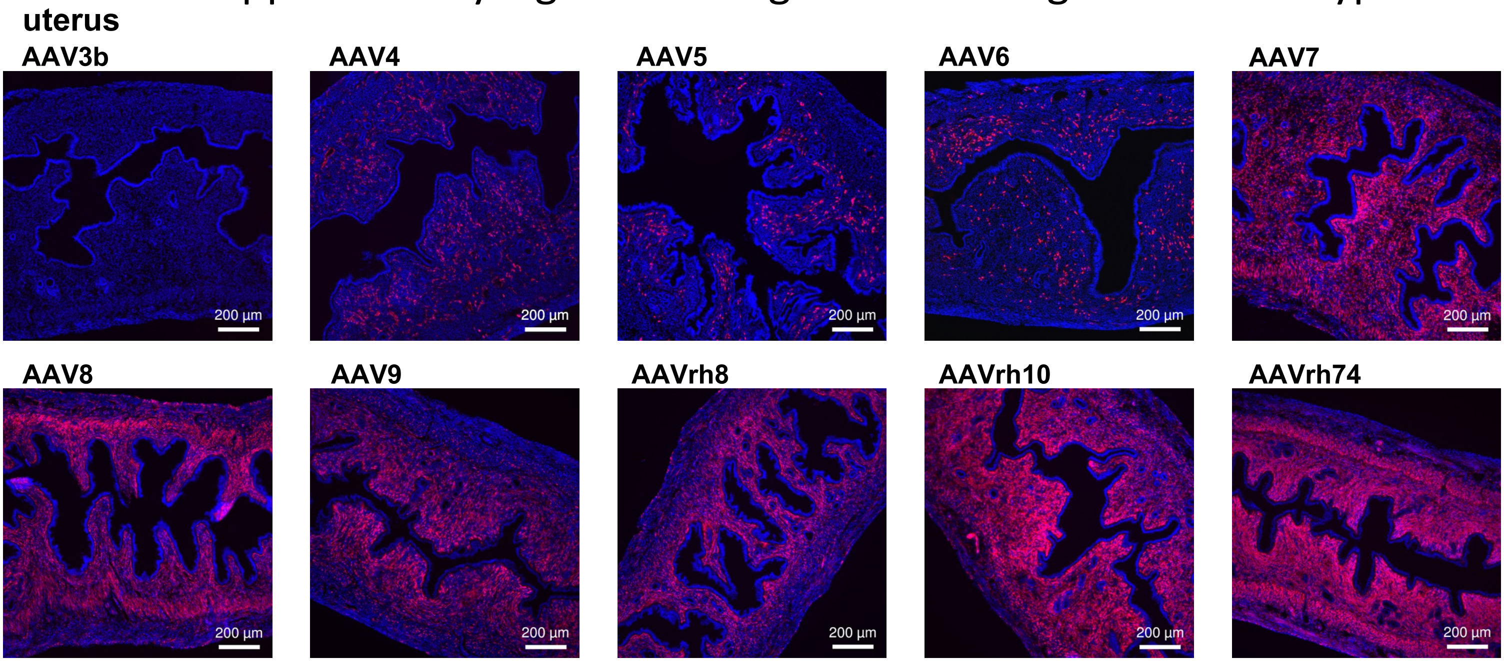

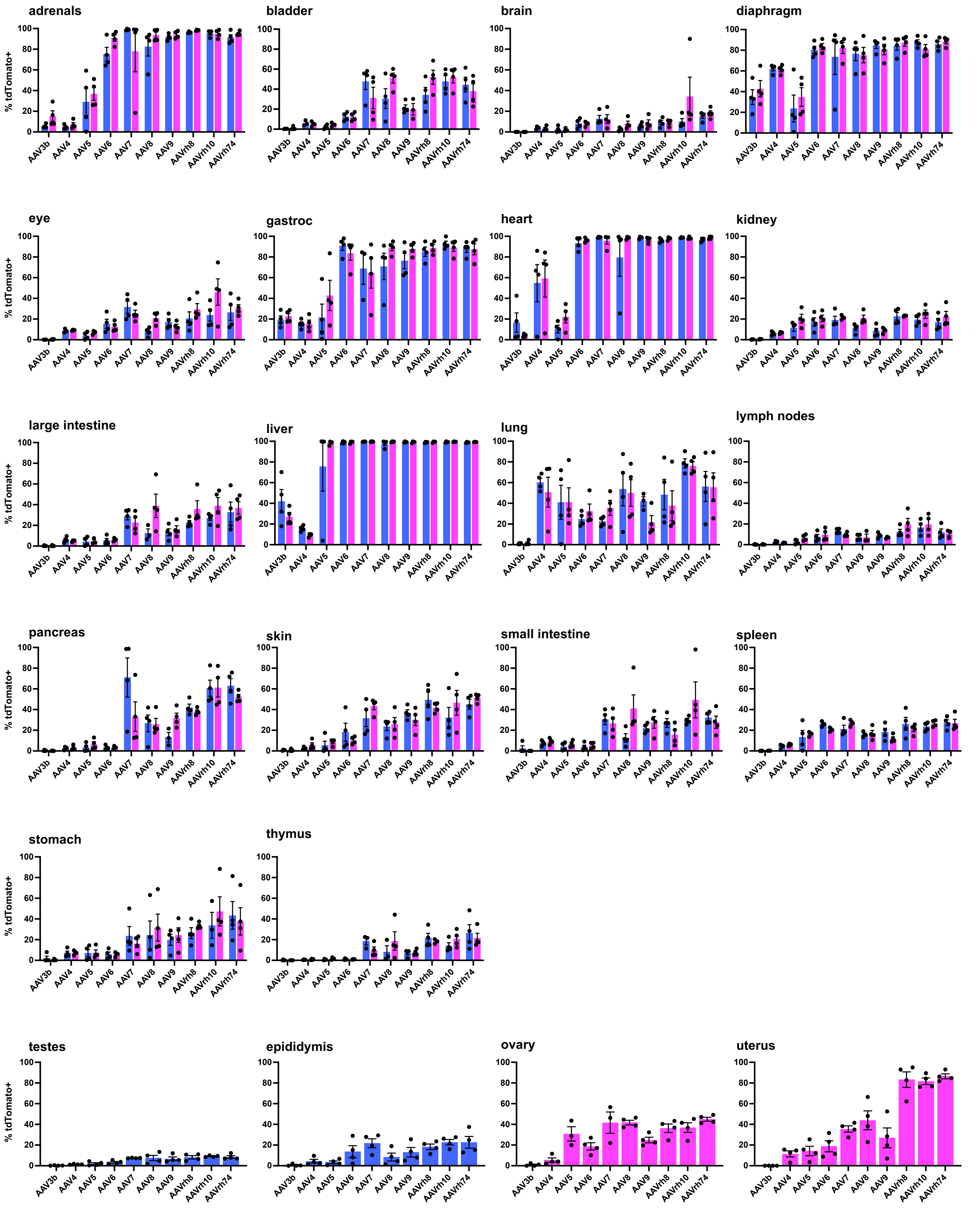


Supplementary Figure 5 - Functional Tropism by individual organs

Males

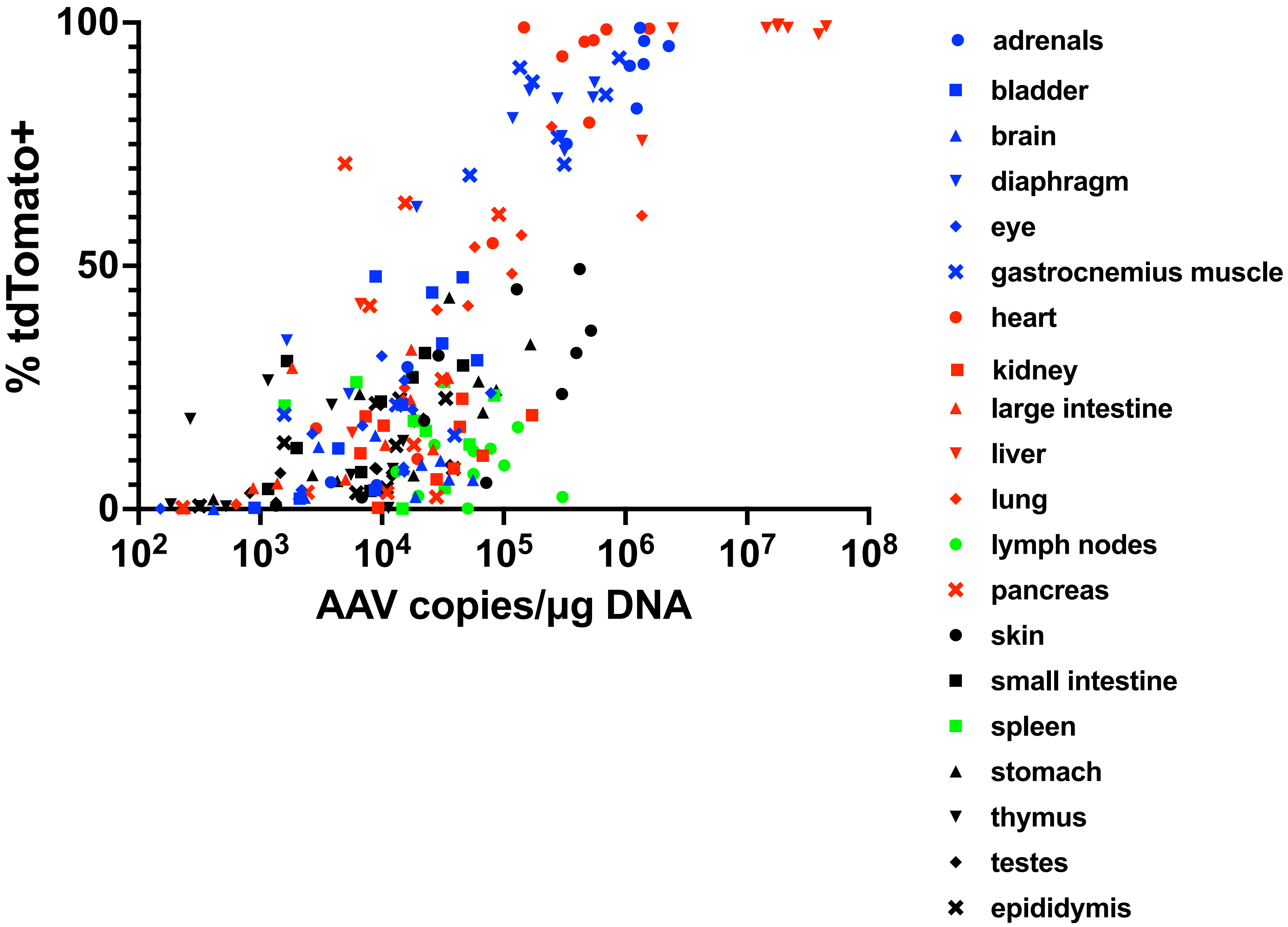


Supplementary Figure 6 - Combined correlations

Females

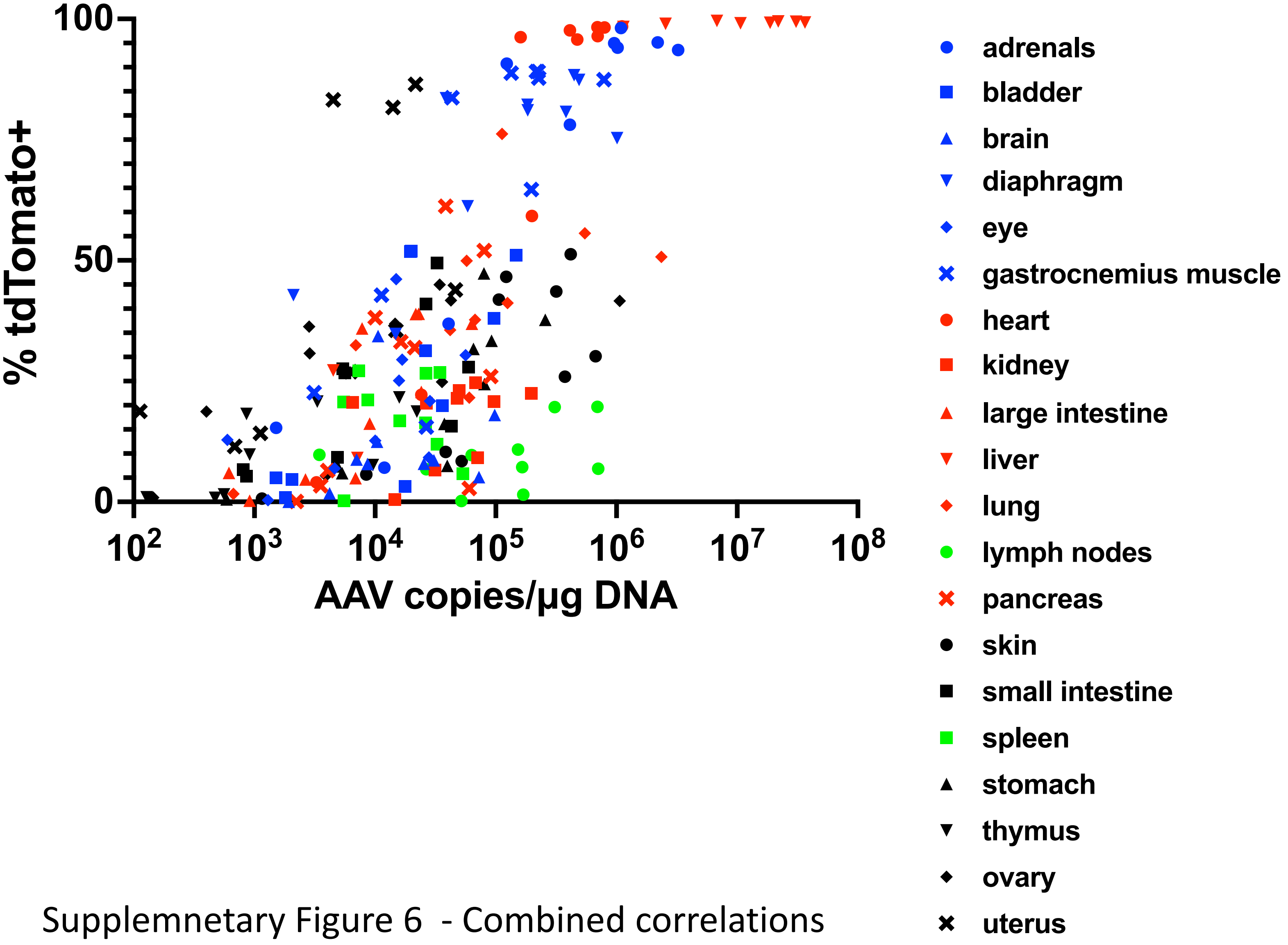


Supplemnetary Figure 6 - Combined correlations

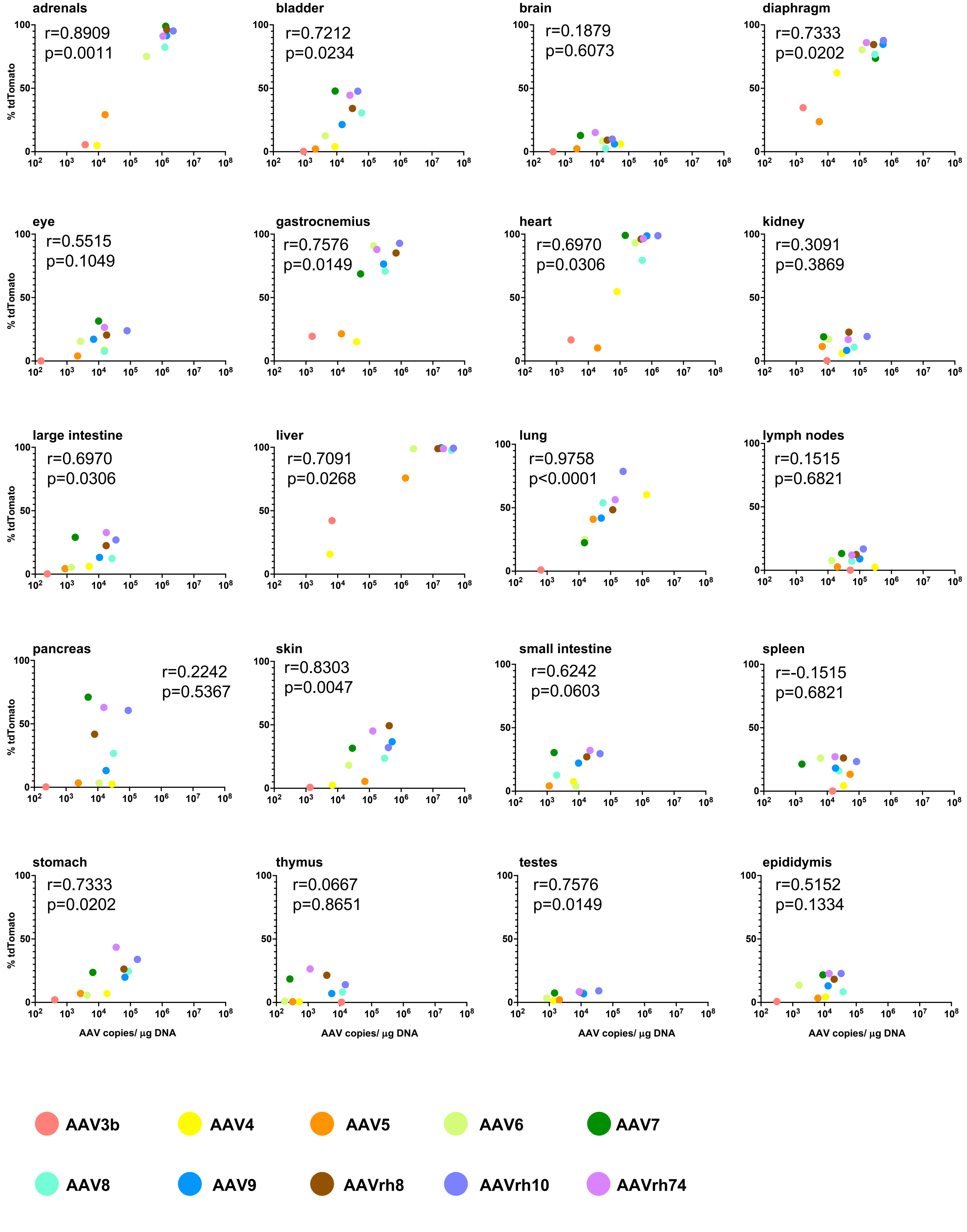


Supplementary Figure 7 - Correlations by individual organs - males

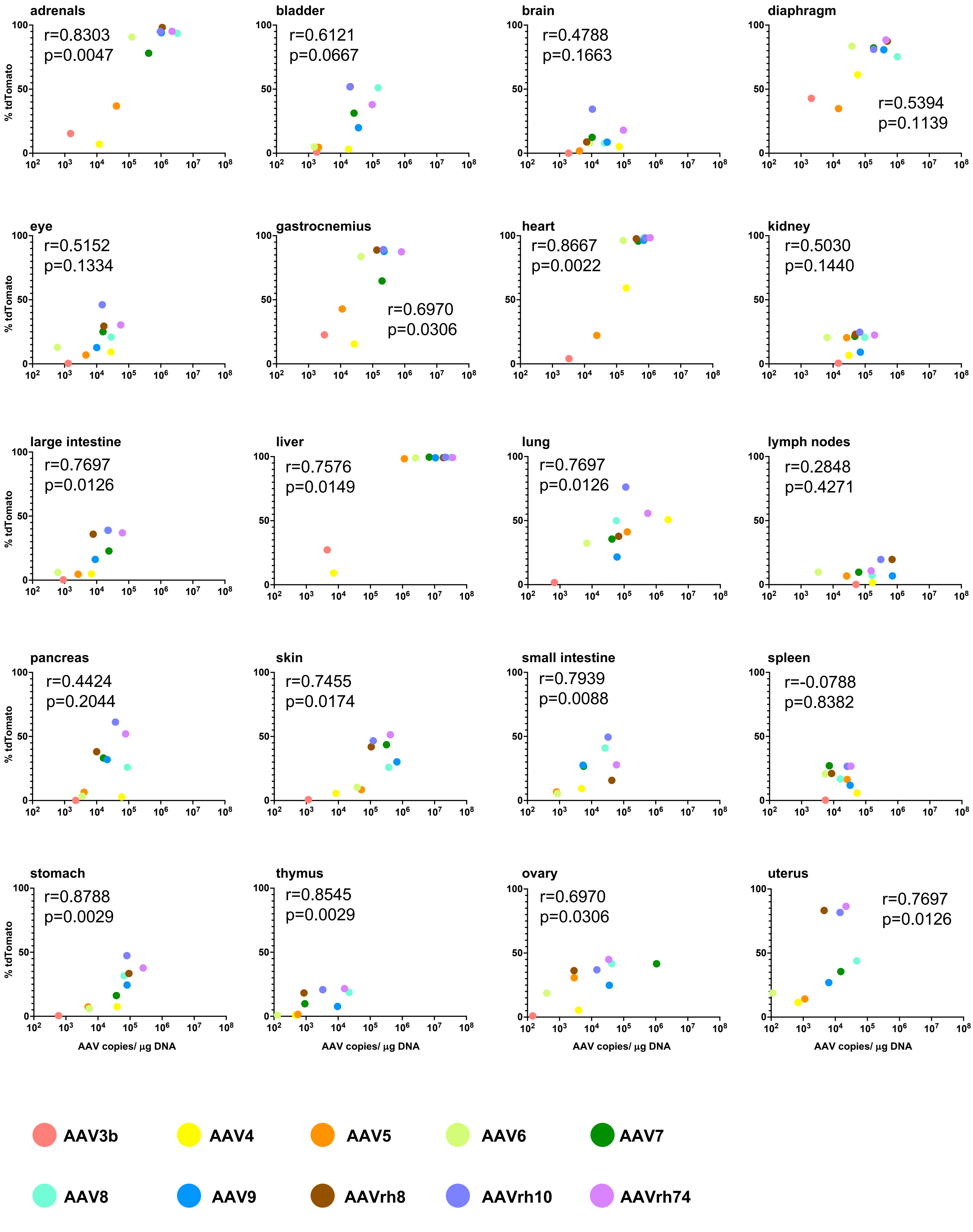


Supplementary Figure 7 - Correlations by individual organs - females

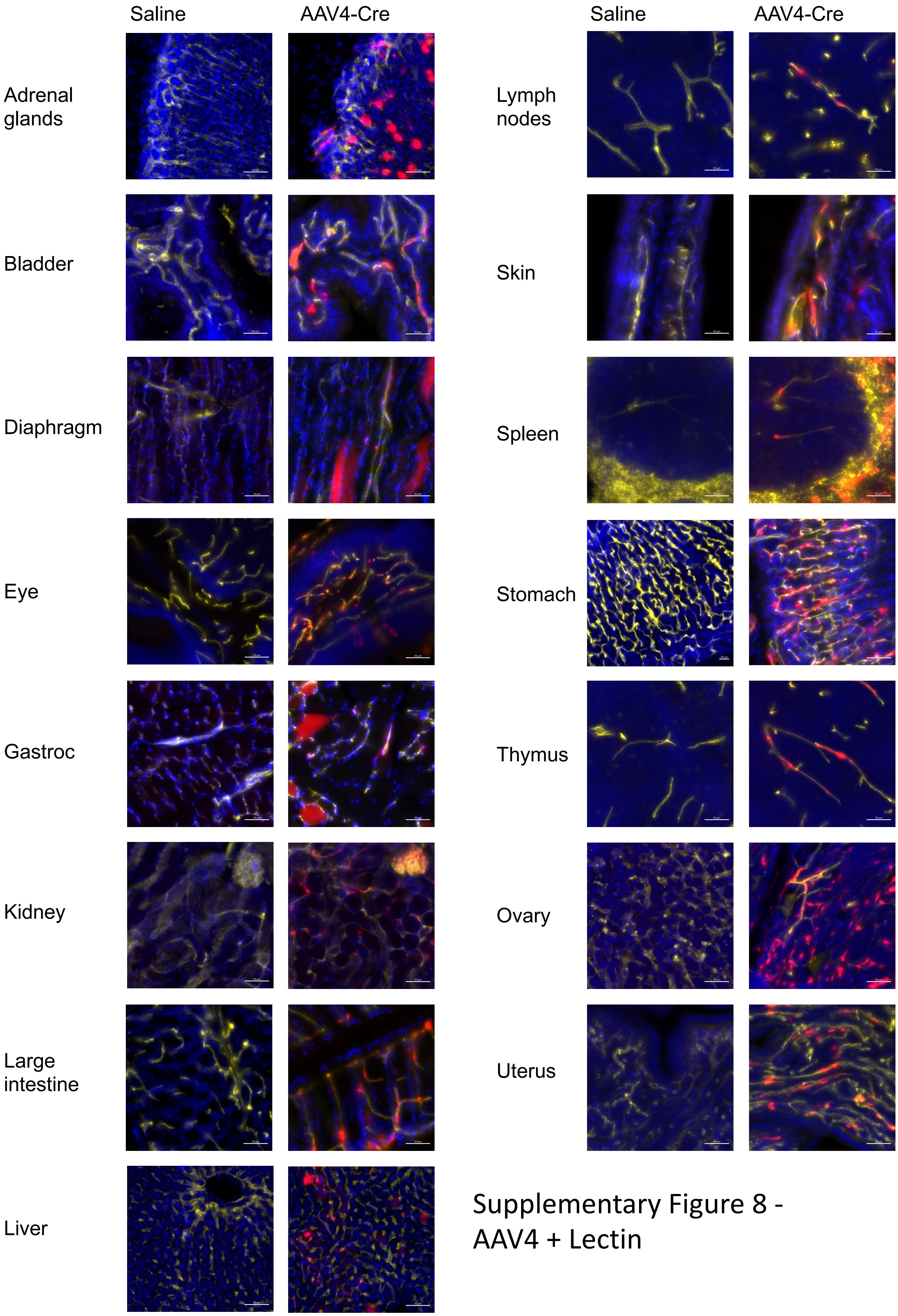


Supplementary Figure 8 - AAV4 + Lectin

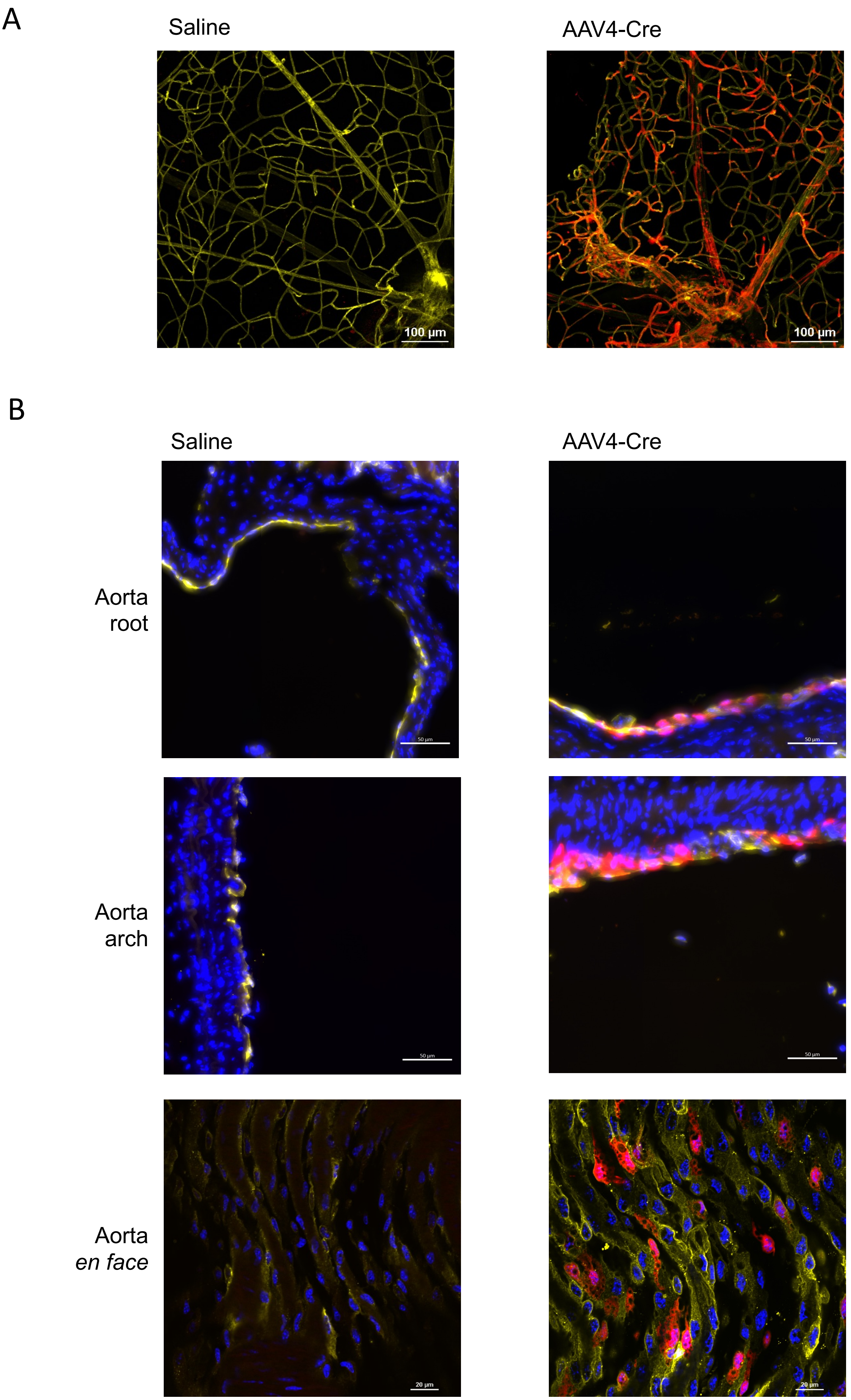


Supplementary Figure 9 - Retina and Aorta

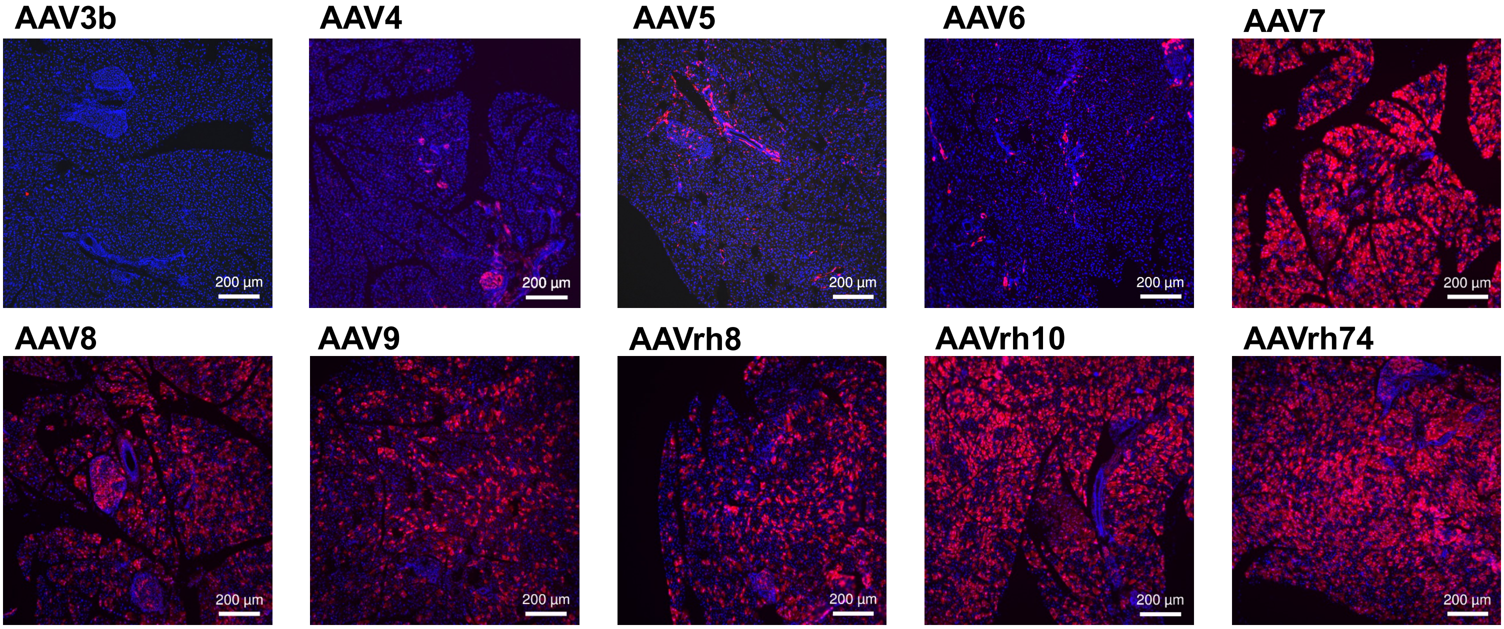


Supplementary Figure 10 – Islet serotype survey

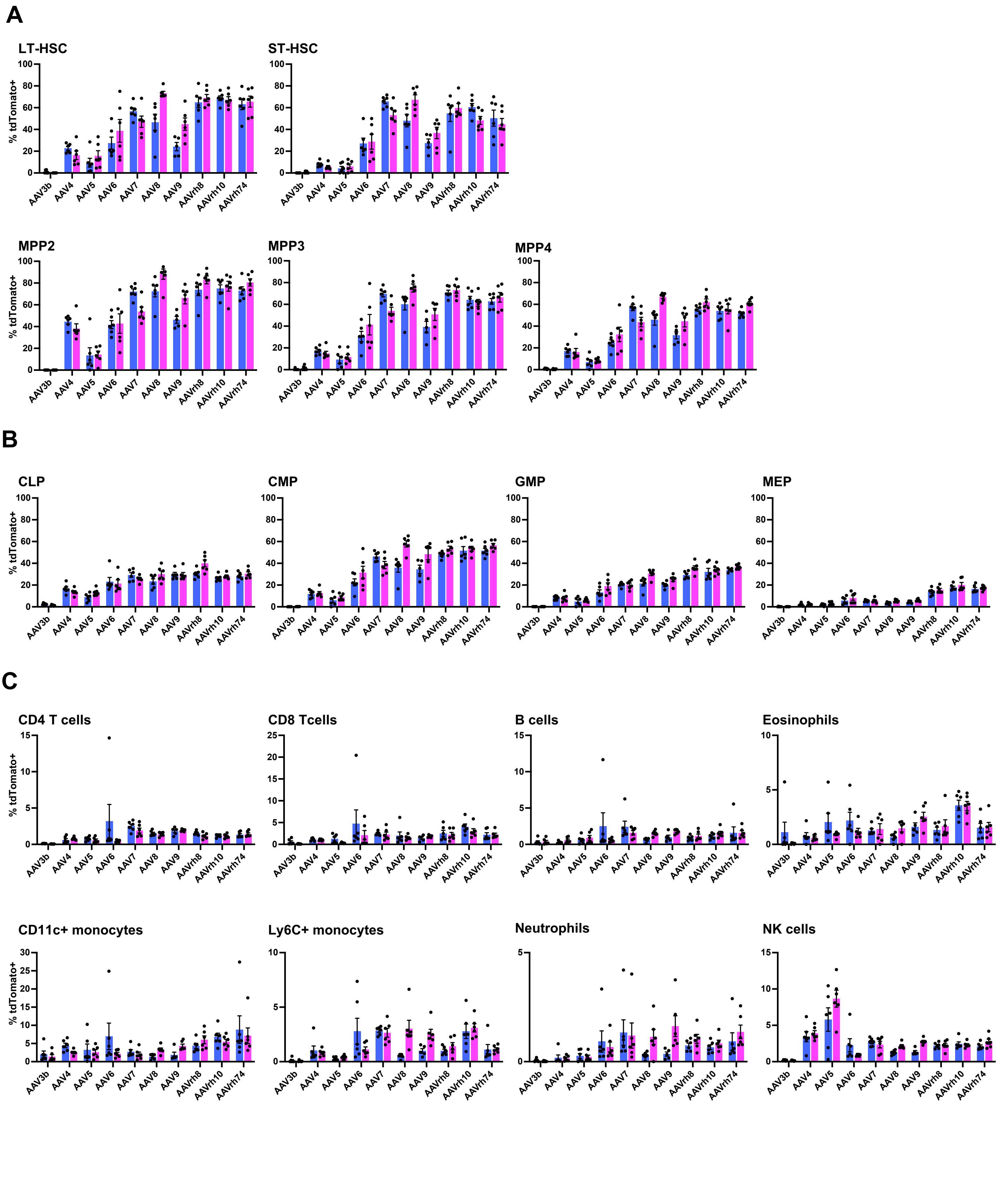


Supplementary Figure 11 - HSC and PBL transduction by cells type

Supplemental Table S1: Capsid sequence identities across serotypes, as percentages

| | AAV3b | AAV4 | AAV5 | AAV6 | AAV7 | AAV8 | AAV9 | AAVrh8 | AAVrh10 | AAVrh74 |
|---|---|---|---|---|---|---|---|---|---|---|
| **AAV3b** | 100 | 64 | 59 | 87 | 85 | 86 | 84 | 85 | 86 | 85 |
| **AAV4** | 64 | 100 | 53 | 63 | 64 | 64 | 63 | 64 | 64 | 64 |
| **AAV5** | 59 | 53 | 100 | 59 | 59 | 58 | 58 | 58 | 58 | 58 |
| **AAV6** | 87 | 63 | 59 | 100 | 85 | 84 | 83 | 86 | 85 | 84 |
| **AAV7** | 85 | 64 | 59 | 85 | 100 | 88 | 82 | 87 | 89 | 88 |
| **AAV8** | 86 | 64 | 58 | 84 | 88 | 100 | 85 | 91 | 94 | 93 |
| **AAV9** | 84 | 63 | 58 | 83 | 82 | 85 | 100 | 87 | 86 | 86 |
| **AAVrh8** | 85 | 64 | 58 | 86 | 87 | 91 | 87 | 100 | 87 | 91 |
| **AAVrh10** | 86 | 64 | 58 | 85 | 89 | 94 | 86 | 87 | 100 | 99 |
| **AAVrh74** | 85 | 64 | 58 | 84 | 88 | 93 | 86 | 91 | 99 | 100 |

| Supplemental Table S2: Organs for biodistribution and functional transduction analysis | | | | |
|---|---|---|---|---|
| Adrenal glands | Bladder | Brain | Diaphragm | Eye |
| Gastrocnemius muscle | Heart | Kidney | Large Intestine | Liver |
| Lung | Lymph nodes | Pancreas | Skin | Small intestine |
| Spleen | Stomach | Thymus | Testes/Ovaries | Epididymis/Uterus |
| Bone Marrow | Blood | | | |

Supplemental Table S3 – Wilcoxon Rank Tests – Biodistribution Male

**Adrenals**

| | AAV3b | AAV4 | AAV5 | AAV6 | AAV7 | AAV8 | AAV9 | AAVrh8 | AAVrh10 |
|---|---|---|---|---|---|---|---|---|---|
| AAV4 | 0.521 | - | - | - | - | - | - | - | - |
| AAV5 | 0.057 | 0.449 | - | - | - | - | - | - | - |
| AAV6 | 0.010 | 0.010 | 0.010 | - | - | - | - | - | - |
| AAV7 | 0.010 | 0.010 | 0.010 | 0.069 | - | - | - | - | - |
| AAV8 | 0.010 | 0.010 | 0.010 | 0.069 | 0.679 | - | - | - | - |
| AAV9 | 0.016 | 0.016 | 0.010 | 0.017 | 0.521 | 1.000 | - | - | - |
| AAVrh8 | 0.010 | 0.010 | 0.010 | 0.192 | 0.959 | 0.767 | 0.959 | - | - |
| AAVrh10 | 0.010 | 0.010 | 0.010 | 0.010 | 0.140 | 0.069 | 0.132 | 0.338 | - |
| AAVrh74 | 0.010 | 0.010 | 0.010 | 0.574 | 0.521 | 0.877 | 0.521 | 0.767 | 0.140 |

**Bladder**

| | AAV3b | AAV4 | AAV5 | AAV6 | AAV7 | AAV8 | AAV9 | AAVrh8 | AAVrh10 |
|---|---|---|---|---|---|---|---|---|---|
| AAV4 | 0.400 | - | - | - | - | - | - | - | - |
| AAV5 | 0.253 | 0.818 | - | - | - | - | - | - | - |
| AAV6 | 0.111 | 0.818 | 0.398 | - | - | - | - | - | - |
| AAV7 | 0.030 | 0.253 | 0.046 | 0.253 | - | - | - | - | - |
| AAV8 | 0.030 | 0.076 | 0.030 | 0.046 | 0.165 | - | - | - | - |
| AAV9 | 0.030 | 0.253 | 0.046 | 0.161 | 0.616 | 0.165 | - | - | - |
| AAVrh8 | 0.028 | 0.220 | 0.030 | 0.165 | 0.616 | 0.400 | 0.818 | - | - |
| AAVrh10 | 0.028 | 0.046 | 0.028 | 0.028 | 0.030 | 0.727 | 0.161 | 0.327 | - |
| AAVrh74 | 0.028 | 0.097 | 0.028 | 0.028 | 0.111 | 0.327 | 0.508 | 0.749 | 0.253 |

**Brain**

| | AAV3b | AAV4 | AAV5 | AAV6 | AAV7 | AAV8 | AAV9 | AAVrh8 | AAVrh10 |
|---|---|---|---|---|---|---|---|---|---|
| AAV4 | 0.238 | - | - | - | - | - | - | - | - |
| AAV5 | 0.030 | 0.537 | - | - | - | - | - | - | - |
| AAV6 | 0.018 | 0.807 | 0.103 | - | - | - | - | - | - |
| AAV7 | 0.018 | 0.537 | 0.537 | 0.139 | - | - | - | - | - |
| AAV8 | 0.018 | 0.937 | 0.103 | 0.898 | 0.139 | - | - | - | - |
| AAV9 | 0.018 | 0.567 | 0.018 | 0.161 | 0.018 | 0.161 | - | - | - |
| AAVrh8 | 0.018 | 0.898 | 0.103 | 0.937 | 0.175 | 0.937 | 0.299 | - | - |
| AAVrh10 | 0.018 | 0.937 | 0.018 | 0.537 | 0.018 | 0.697 | 0.671 | 0.299 | - |
| AAVrh74 | 0.018 | 0.623 | 0.030 | 0.480 | 0.049 | 0.480 | 0.052 | 0.697 | 0.139 |

**Diaphragm**

| | AAV3b | AAV4 | AAV5 | AAV6 | AAV7 | AAV8 | AAV9 | AAVrh8 | AAVrh10 |
|---|---|---|---|---|---|---|---|---|---|
| AAV4 | 0.435 | - | - | - | - | - | - | - | - |
| AAV5 | 0.142 | 0.749 | - | - | - | - | - | - | - |
| AAV6 | 0.011 | 0.028 | 0.011 | - | - | - | - | - | - |
| AAV7 | 0.011 | 0.028 | 0.011 | 0.559 | - | - | - | - | - |
| AAV8 | 0.011 | 0.011 | 0.011 | 0.028 | 0.749 | - | - | - | - |
| AAV9 | 0.020 | 0.011 | 0.011 | 0.142 | 0.508 | 0.937 | - | - | - |
| AAVrh8 | 0.011 | 0.011 | 0.011 | 0.506 | 0.856 | 0.349 | 0.508 | - | - |
| AAVrh10 | 0.011 | 0.011 | 0.011 | 0.028 | 0.422 | 0.349 | 0.508 | 0.349 | - |
| AAVrh74 | 0.011 | 0.021 | 0.011 | 0.662 | 0.937 | 0.289 | 0.209 | 0.422 | 0.028 |

Supplemental Table S3 – Wilcoxon Rank Tests – Biodistribution Male

**Eye**

| | AAV3b | AAV4 | AAV5 | AAV6 | AAV7 | AAV8 | AAV9 | AAVrh8 | AAVrh10 |
|---|---|---|---|---|---|---|---|---|---|
| AAV4 | 0.097 | - | - | - | - | - | - | - | - |
| AAV5 | 0.090 | 0.217 | - | - | - | - | - | - | - |
| AAV6 | 0.062 | 0.217 | 0.944 | - | - | - | - | - | - |
| AAV7 | 0.024 | 0.974 | 0.289 | 0.572 | - | - | - | - | - |
| AAV8 | 0.024 | 0.974 | 0.062 | 0.103 | 0.682 | - | - | - | - |
| AAV9 | 0.024 | 0.874 | 0.289 | 0.217 | 0.974 | 0.494 | - | - | - |
| AAVrh8 | 0.024 | 0.944 | 0.103 | 0.146 | 0.874 | 1.000 | 0.383 | - | - |
| AAVrh10 | 0.024 | 0.123 | 0.024 | 0.024 | 0.062 | 0.103 | 0.065 | 0.182 | - |
| AAVrh74 | 0.024 | 1.000 | 0.182 | 0.182 | 0.874 | 0.803 | 0.974 | 0.944 | 0.103 |

**Gastrocnemius muscle**

| | AAV3b | AAV4 | AAV5 | AAV6 | AAV7 | AAV8 | AAV9 | AAVrh8 | AAVrh10 |
|---|---|---|---|---|---|---|---|---|---|
| AAV4 | 0.482 | - | - | - | - | - | - | - | - |
| AAV5 | 0.047 | 0.616 | - | - | - | - | - | - | - |
| AAV6 | 0.010 | 0.010 | 0.010 | - | - | - | - | - | - |
| AAV7 | 0.010 | 0.616 | 0.010 | 0.010 | - | - | - | - | - |
| AAV8 | 0.010 | 0.010 | 0.010 | 0.089 | 0.010 | - | - | - | - |
| AAV9 | 0.031 | 0.033 | 0.020 | 0.313 | 0.033 | 0.931 | - | - | - |
| AAVrh8 | 0.010 | 0.010 | 0.010 | 0.060 | 0.010 | 0.367 | 0.523 | - | - |
| AAVrh10 | 0.010 | 0.010 | 0.010 | 0.010 | 0.010 | 0.089 | 0.057 | 0.308 | - |
| AAVrh74 | 0.017 | 0.047 | 0.010 | 0.931 | 0.047 | 0.308 | 0.476 | 0.109 | 0.033 |

**Heart**

| | AAV3b | AAV4 | AAV5 | AAV6 | AAV7 | AAV8 | AAV9 | AAVrh8 | AAVrh10 |
|---|---|---|---|---|---|---|---|---|---|
| AAV4 | 0.300 | - | - | - | - | - | - | - | - |
| AAV5 | 0.043 | 0.631 | - | - | - | - | - | - | - |
| AAV6 | 0.008 | 0.064 | 0.023 | - | - | - | - | - | - |
| AAV7 | 0.008 | 0.231 | 0.008 | 0.367 | - | - | - | - | - |
| AAV8 | 0.008 | 0.043 | 0.008 | 0.818 | 0.231 | - | - | - | - |
| AAV9 | 0.043 | 0.043 | 0.043 | 0.043 | 0.043 | 0.319 | - | - | - |
| AAVrh8 | 0.008 | 0.013 | 0.008 | 0.559 | 0.013 | 0.715 | 0.227 | - | - |
| AAVrh10 | 0.008 | 0.008 | 0.008 | 0.043 | 0.008 | 0.097 | 0.227 | 0.064 | - |
| AAVrh74 | 0.008 | 0.023 | 0.008 | 0.192 | 0.038 | 0.631 | 0.616 | 0.715 | 0.043 |

**Kidney**

| | AAV3b | AAV4 | AAV5 | AAV6 | AAV7 | AAV8 | AAV9 | AAVrh8 | AAVrh10 |
|---|---|---|---|---|---|---|---|---|---|
| AAV4 | 0.837 | - | - | - | - | - | - | - | - |
| AAV5 | 0.837 | 0.606 | - | - | - | - | - | - | - |
| AAV6 | 0.716 | 0.807 | 0.373 | - | - | - | - | - | - |
| AAV7 | 0.837 | 0.606 | 0.837 | 0.435 | - | - | - | - | - |
| AAV8 | 0.040 | 0.289 | 0.056 | 0.080 | 0.080 | - | - | - | - |
| AAV9 | 0.097 | 0.449 | 0.016 | 0.016 | 0.016 | 0.289 | - | - | - |
| AAVrh8 | 0.056 | 0.289 | 0.014 | 0.014 | 0.014 | 0.435 | 0.837 | - | - |
| AAVrh10 | 0.014 | 0.026 | 0.014 | 0.014 | 0.014 | 0.238 | 0.026 | 0.026 | - |
| AAVrh74 | 0.056 | 0.435 | 0.016 | 0.040 | 0.016 | 0.521 | 1.000 | 0.807 | 0.056 |

Supplemental Table S3 – Wilcoxon Rank Tests – Biodistribution Male

**Large Intestine**

| | AAV3b | AAV4 | AAV5 | AAV6 | AAV7 | AAV8 | AAV9 | AAVrh8 | AAVrh10 |
|---|---|---|---|---|---|---|---|---|---|
| AAV4 | 0.327 | - | - | - | - | - | - | - | - |
| AAV5 | 0.022 | 0.679 | - | - | - | - | - | - | - |
| AAV6 | 0.010 | 0.877 | 0.679 | - | - | - | - | - | - |
| AAV7 | 0.010 | 0.959 | 0.074 | 0.387 | - | - | - | - | - |
| AAV8 | 0.010 | 0.049 | 0.033 | 0.033 | 0.112 | - | - | - | - |
| AAV9 | 0.015 | 0.327 | 0.015 | 0.022 | 0.022 | 0.261 | - | - | - |
| AAVrh8 | 0.010 | 0.155 | 0.015 | 0.049 | 0.049 | 0.387 | 0.959 | - | - |
| AAVrh10 | 0.010 | 0.033 | 0.010 | 0.010 | 0.010 | 0.787 | 0.202 | 0.327 | - |
| AAVrh74 | 0.010 | 0.261 | 0.010 | 0.022 | 0.022 | 0.679 | 0.870 | 1.000 | 0.261 |

**Liver**

| | AAV3b | AAV4 | AAV5 | AAV6 | AAV7 | AAV8 | AAV9 | AAVrh8 | AAVrh10 |
|---|---|---|---|---|---|---|---|---|---|
| AAV4 | 1.000 | - | - | - | - | - | - | - | - |
| AAV5 | 0.011 | 0.011 | - | - | - | - | - | - | - |
| AAV6 | 0.011 | 0.011 | 0.467 | - | - | - | - | - | - |
| AAV7 | 0.014 | 0.014 | 0.011 | 0.011 | - | - | - | - | - |
| AAV8 | 0.014 | 0.014 | 0.011 | 0.011 | 0.076 | - | - | - | - |
| AAV9 | 0.014 | 0.014 | 0.011 | 0.011 | 0.616 | 0.045 | - | - | - |
| AAVrh8 | 0.014 | 0.014 | 0.011 | 0.014 | 0.860 | 0.024 | 0.616 | - | - |
| AAVrh10 | 0.011 | 0.011 | 0.011 | 0.011 | 0.044 | 0.829 | 0.011 | 0.014 | - |
| AAVrh74 | 0.011 | 0.011 | 0.024 | 0.086 | 0.400 | 0.222 | 0.727 | 0.829 | 0.120 |

**Lung**

| | AAV3b | AAV4 | AAV5 | AAV6 | AAV7 | AAV8 | AAV9 | AAVrh8 | AAVrh10 |
|---|---|---|---|---|---|---|---|---|---|
| AAV4 | 0.011 | - | - | - | - | - | - | - | - |
| AAV5 | 0.011 | 0.360 | - | - | - | - | - | - | - |
| AAV6 | 0.011 | 0.360 | 0.479 | - | - | - | - | - | - |
| AAV7 | 0.011 | 0.479 | 0.715 | 0.646 | - | - | - | - | - |
| AAV8 | 0.011 | 0.479 | 0.168 | 0.065 | 0.065 | - | - | - | - |
| AAV9 | 0.015 | 0.508 | 0.463 | 0.102 | 0.072 | 0.616 | - | - | - |
| AAVrh8 | 0.011 | 0.479 | 0.045 | 0.015 | 0.011 | 0.289 | 0.217 | - | - |
| AAVrh10 | 0.011 | 0.479 | 0.045 | 0.015 | 0.011 | 0.093 | 0.049 | 0.122 | - |
| AAVrh74 | 0.015 | 0.792 | 0.289 | 0.102 | 0.102 | 0.693 | 0.616 | 0.693 | 0.463 |

**Lymph nodes**

| | AAV3b | AAV4 | AAV5 | AAV6 | AAV7 | AAV8 | AAV9 | AAVrh8 | AAVrh10 |
|---|---|---|---|---|---|---|---|---|---|
| AAV4 | 0.383 | - | - | - | - | - | - | - | - |
| AAV5 | 0.435 | 0.071 | - | - | - | - | - | - | - |
| AAV6 | 0.199 | 0.071 | 0.435 | - | - | - | - | - | - |
| AAV7 | 0.767 | 0.071 | 0.435 | 0.248 | - | - | - | - | - |
| AAV8 | 0.590 | 0.679 | 0.083 | 0.071 | 0.154 | - | - | - | - |
| AAV9 | 0.307 | 0.818 | 0.071 | 0.071 | 0.091 | 0.383 | - | - | - |
| AAVrh8 | 0.506 | 0.506 | 0.103 | 0.071 | 0.248 | 0.818 | 0.636 | - | - |
| AAVrh10 | 0.103 | 0.248 | 0.065 | 0.049 | 0.049 | 0.071 | 0.536 | 0.199 | - |
| AAVrh74 | 0.767 | 0.506 | 0.103 | 0.083 | 0.383 | 0.818 | 0.307 | 0.818 | 0.083 |

Supplemental Table S3 – Wilcoxon Rank Tests – Biodistribution Male

**Pancreas**

| | AAV3b | AAV4 | AAV5 | AAV6 | AAV7 | AAV8 | AAV9 | AAVrh8 | AAVrh10 |
|---|---|---|---|---|---|---|---|---|---|
| AAV4 | 0.195 | - | - | - | - | - | - | - | - |
| AAV5 | 0.019 | 0.623 | - | - | - | - | - | - | - |
| AAV6 | 0.069 | 0.697 | 0.537 | - | - | - | - | - | - |
| AAV7 | 0.019 | 0.727 | 0.229 | 0.959 | - | - | - | - | - |
| AAV8 | 0.016 | 0.697 | 0.028 | 0.229 | 0.185 | - | - | - | - |
| AAV9 | 0.019 | 1.000 | 0.028 | 0.478 | 0.195 | 0.727 | - | - | - |
| AAVrh8 | 0.016 | 0.749 | 0.043 | 0.697 | 0.567 | 0.229 | 0.296 | - | - |
| AAVrh10 | 0.016 | 0.229 | 0.016 | 0.028 | 0.019 | 0.154 | 0.076 | 0.016 | - |
| AAVrh74 | 0.016 | 0.959 | 0.043 | 0.383 | 0.383 | 0.537 | 0.727 | 0.464 | 0.028 |

**Skin**

| | AAV3b | AAV4 | AAV5 | AAV6 | AAV7 | AAV8 | AAV9 | AAVrh8 | AAVrh10 |
|---|---|---|---|---|---|---|---|---|---|
| AAV4 | 0.260 | - | - | - | - | - | - | - | - |
| AAV5 | 0.024 | 0.032 | - | - | - | - | - | - | - |
| AAV6 | 0.024 | 0.049 | 0.662 | - | - | - | - | - | - |
| AAV7 | 0.024 | 0.123 | 0.837 | 0.732 | - | - | - | - | - |
| AAV8 | 0.024 | 0.022 | 0.101 | 0.024 | 0.032 | - | - | - | - |
| AAV9 | 0.032 | 0.024 | 0.032 | 0.022 | 0.022 | 0.411 | - | - | - |
| AAVrh8 | 0.024 | 0.022 | 0.045 | 0.022 | 0.022 | 0.937 | 0.710 | - | - |
| AAVrh10 | 0.024 | 0.022 | 0.045 | 0.022 | 0.022 | 0.662 | 0.710 | 0.662 | - |
| AAVrh74 | 0.032 | 0.024 | 0.317 | 0.032 | 0.049 | 0.521 | 0.138 | 0.317 | 0.177 |

**Small Intestine**

| | AAV3b | AAV4 | AAV5 | AAV6 | AAV7 | AAV8 | AAV9 | AAVrh8 | AAVrh10 |
|---|---|---|---|---|---|---|---|---|---|
| AAV4 | 0.506 | - | - | - | - | - | - | - | - |
| AAV5 | 0.380 | 0.380 | - | - | - | - | - | - | - |
| AAV6 | 0.091 | 0.974 | 1.000 | - | - | - | - | - | - |
| AAV7 | 0.019 | 0.506 | 0.974 | 0.764 | - | - | - | - | - |
| AAV8 | 0.380 | 0.784 | 1.000 | 0.695 | 0.685 | - | - | - | - |
| AAV9 | 0.039 | 0.722 | 0.135 | 0.317 | 0.150 | 0.367 | - | - | - |
| AAVrh8 | 0.019 | 0.311 | 0.066 | 0.246 | 0.024 | 0.195 | 0.481 | - | - |
| AAVrh10 | 0.019 | 0.024 | 0.039 | 0.065 | 0.019 | 0.077 | 0.107 | 0.109 | - |
| AAVrh74 | 0.019 | 0.311 | 0.039 | 0.130 | 0.024 | 0.195 | 0.481 | 0.784 | 0.248 |

**Spleen**

| | AAV3b | AAV4 | AAV5 | AAV6 | AAV7 | AAV8 | AAV9 | AAVrh8 | AAVrh10 |
|---|---|---|---|---|---|---|---|---|---|
| AAV4 | 0.828 | - | - | - | - | - | - | - | - |
| AAV5 | 0.073 | 0.464 | - | - | - | - | - | - | - |
| AAV6 | 0.190 | 0.311 | 0.024 | - | - | - | - | - | - |
| AAV7 | 0.024 | 0.030 | 0.024 | 0.030 | - | - | - | - | - |
| AAV8 | 0.567 | 0.849 | 0.311 | 0.076 | 0.024 | - | - | - | - |
| AAV9 | 1.000 | 0.849 | 0.117 | 0.176 | 0.024 | 0.685 | - | - | - |
| AAVrh8 | 0.849 | 0.849 | 0.496 | 0.399 | 0.031 | 0.925 | 0.925 | - | - |
| AAVrh10 | 0.024 | 0.117 | 0.397 | 0.024 | 0.024 | 0.030 | 0.030 | 0.317 | - |
| AAVrh74 | 0.537 | 0.623 | 0.073 | 0.828 | 0.030 | 0.311 | 0.478 | 0.311 | 0.076 |

Supplemental Table S3 – Wilcoxon Rank Tests – Biodistribution Male

**Stomach**

| | AAV3b | AAV4 | AAV5 | AAV6 | AAV7 | AAV8 | AAV9 | AAVrh8 | AAVrh10 |
|---|---|---|---|---|---|---|---|---|---|
| AAV4 | 0.175 | - | - | - | - | - | - | - | - |
| AAV5 | 0.028 | 0.662 | - | - | - | - | - | - | - |
| AAV6 | 0.010 | 0.749 | 0.662 | - | - | - | - | - | - |
| AAV7 | 0.008 | 0.837 | 0.094 | 0.300 | - | - | - | - | - |
| AAV8 | 0.008 | 0.094 | 0.028 | 0.028 | 0.047 | - | - | - | - |
| AAV9 | 0.010 | 0.052 | 0.010 | 0.010 | 0.010 | 0.931 | - | - | - |
| AAVrh8 | 0.008 | 0.131 | 0.008 | 0.008 | 0.020 | 0.837 | 0.727 | - | - |
| AAVrh10 | 0.008 | 0.010 | 0.008 | 0.008 | 0.008 | 0.175 | 0.084 | 0.094 | - |
| AAVrh74 | 0.008 | 0.231 | 0.008 | 0.008 | 0.010 | 0.376 | 0.084 | 0.662 | 0.028 |

**Thymus**

| | AAV3b | AAV4 | AAV5 | AAV6 | AAV7 | AAV8 | AAV9 | AAVrh8 | AAVrh10 |
|---|---|---|---|---|---|---|---|---|---|
| AAV4 | 0.767 | - | - | - | - | - | - | - | - |
| AAV5 | 0.150 | 0.764 | - | - | - | - | - | - | - |
| AAV6 | 0.736 | 0.767 | 0.435 | - | - | - | - | - | - |
| AAV7 | 0.192 | 0.877 | 0.937 | 0.623 | - | - | - | - | - |
| AAV8 | 0.192 | 0.133 | 0.192 | 0.109 | 0.150 | - | - | - | - |
| AAV9 | 0.150 | 0.085 | 0.055 | 0.028 | 0.039 | 0.937 | - | - | - |
| AAVrh8 | 0.133 | 0.055 | 0.056 | 0.028 | 0.039 | 0.937 | 0.764 | - | - |
| AAVrh10 | 0.133 | 0.024 | 0.028 | 0.024 | 0.024 | 0.150 | 0.130 | 0.150 | - |
| AAVrh74 | 0.133 | 0.253 | 0.085 | 0.039 | 0.055 | 0.328 | 0.764 | 0.150 | 0.024 |

**Testes**

| | AAV3b | AAV4 | AAV5 | AAV6 | AAV7 | AAV8 | AAV9 | AAVrh8 | AAVrh10 |
|---|---|---|---|---|---|---|---|---|---|
| AAV4 | 0.008 | - | - | - | - | - | - | - | - |
| AAV5 | 0.011 | 0.662 | - | - | - | - | - | - | - |
| AAV6 | 0.018 | 0.662 | 0.492 | - | - | - | - | - | - |
| AAV7 | 0.008 | 0.309 | 0.959 | 0.662 | - | - | - | - | - |
| AAV8 | 0.008 | 0.043 | 0.180 | 0.180 | 0.180 | - | - | - | - |
| AAV9 | 0.011 | 0.011 | 0.011 | 0.011 | 0.011 | 0.849 | - | - | - |
| AAVrh8 | 0.008 | 0.018 | 0.043 | 0.008 | 0.018 | 0.767 | 0.235 | - | - |
| AAVrh10 | 0.008 | 0.008 | 0.008 | 0.008 | 0.008 | 0.180 | 0.180 | 0.028 | - |
| AAVrh74 | 0.008 | 0.028 | 0.066 | 0.008 | 0.018 | 1.000 | 0.521 | 0.856 | 0.043 |

**Epididymis**

| | AAV3b | AAV4 | AAV5 | AAV6 | AAV7 | AAV8 | AAV9 | AAVrh8 | AAVrh10 |
|---|---|---|---|---|---|---|---|---|---|
| AAV4 | 0.257 | - | - | - | - | - | - | - | - |
| AAV5 | 0.039 | 0.850 | - | - | - | - | - | - | - |
| AAV6 | 0.138 | 0.611 | 0.057 | - | - | - | - | - | - |
| AAV7 | 0.032 | 1.000 | 1.000 | 0.116 | - | - | - | - | - |
| AAV8 | 0.032 | 0.323 | 0.116 | 0.039 | 0.323 | - | - | - | - |
| AAV9 | 0.039 | 0.914 | 0.732 | 0.032 | 0.643 | 0.397 | - | - | - |
| AAVrh8 | 0.032 | 0.682 | 0.682 | 0.039 | 0.397 | 0.779 | 1.000 | - | - |
| AAVrh10 | 0.032 | 0.162 | 0.057 | 0.032 | 0.083 | 0.920 | 0.257 | 0.323 | - |
| AAVrh74 | 0.060 | 0.850 | 1.000 | 0.209 | 0.850 | 0.397 | 0.914 | 1.000 | 0.209 |

Supplemental Table S4 - Wilcoxon Rank Tests - Biodistribution Female

**Adrenals**

| | AAV3b | AAV4 | AAV5 | AAV6 | AAV7 | AAV8 | AAV9 | AAVrh8 | AAVrh10 |
|---|---|---|---|---|---|---|---|---|---|
| AAV4 | 0.170 | - | - | - | - | - | - | - | - |
| AAV5 | 0.007 | 0.024 | - | - | - | - | - | - | - |
| AAV6 | 0.013 | 0.069 | 0.081 | - | - | - | - | - | - |
| AAV7 | 0.007 | 0.007 | 0.007 | 0.007 | - | - | - | - | - |
| AAV8 | 0.007 | 0.007 | 0.007 | 0.007 | 0.007 | - | - | - | - |
| AAV9 | 0.007 | 0.007 | 0.007 | 0.007 | 0.081 | 0.152 | - | - | - |
| AAVrh8 | 0.007 | 0.007 | 0.007 | 0.007 | 0.007 | 0.113 | 0.732 | - | - |
| AAVrh10 | 0.007 | 0.007 | 0.007 | 0.013 | 0.631 | 0.152 | 0.818 | 0.432 | - |
| AAVrh74 | 0.007 | 0.007 | 0.007 | 0.007 | 0.007 | 0.818 | 0.007 | 0.007 | 0.056 |

**Bladder**

| | AAV3b | AAV4 | AAV5 | AAV6 | AAV7 | AAV8 | AAV9 | AAVrh8 | AAVrh10 |
|---|---|---|---|---|---|---|---|---|---|
| AAV4 | 0.202 | - | - | - | - | - | - | - | - |
| AAV5 | 0.642 | 0.202 | - | - | - | - | - | - | - |
| AAV6 | 0.898 | 0.202 | 0.642 | - | - | - | - | - | - |
| AAV7 | 0.019 | 0.898 | 0.015 | 0.014 | - | - | - | - | - |
| AAV8 | 0.014 | 0.015 | 0.014 | 0.014 | 0.015 | - | - | - | - |
| AAV9 | 0.205 | 0.898 | 0.554 | 0.269 | 0.959 | 0.051 | - | - | - |
| AAVrh8 | 0.117 | 1.000 | 0.031 | 0.019 | 0.736 | 0.015 | 0.898 | - | - |
| AAVrh10 | 0.019 | 0.898 | 0.031 | 0.019 | 0.554 | 0.015 | 0.959 | 0.959 | - |
| AAVrh74 | 0.014 | 0.015 | 0.014 | 0.014 | 0.019 | 0.736 | 0.117 | 0.019 | 0.019 |

**Brain**

| | AAV3b | AAV4 | AAV5 | AAV6 | AAV7 | AAV8 | AAV9 | AAVrh8 | AAVrh10 |
|---|---|---|---|---|---|---|---|---|---|
| AAV4 | 0.039 | - | - | - | - | - | - | - | - |
| AAV5 | 0.161 | 0.212 | - | - | - | - | - | - | - |
| AAV6 | 0.109 | 0.188 | 0.536 | - | - | - | - | - | - |
| AAV7 | 0.062 | 0.188 | 0.161 | 0.574 | - | - | - | - | - |
| AAV8 | 0.039 | 0.238 | 0.065 | 0.161 | 0.161 | - | - | - | - |
| AAV9 | 0.039 | 0.188 | 0.039 | 0.083 | 0.146 | 0.877 | - | - | - |
| AAVrh8 | 0.123 | 0.212 | 1.000 | 1.000 | 0.604 | 0.091 | 0.123 | - | - |
| AAVrh10 | 0.083 | 0.188 | 0.161 | 0.574 | 0.767 | 0.238 | 0.109 | 0.604 | - |
| AAVrh74 | 0.039 | 0.974 | 0.039 | 0.039 | 0.039 | 0.161 | 0.398 | 0.039 | 0.039 |

**Diaphragm**

| | AAV3b | AAV4 | AAV5 | AAV6 | AAV7 | AAV8 | AAV9 | AAVrh8 | AAVrh10 |
|---|---|---|---|---|---|---|---|---|---|
| AAV4 | 0.289 | - | - | - | - | - | - | - | - |
| AAV5 | 0.016 | 0.623 | - | - | - | - | - | - | - |
| AAV6 | 0.065 | 0.877 | 0.360 | - | - | - | - | - | - |
| AAV7 | 0.026 | 0.217 | 0.148 | 0.148 | - | - | - | - | - |
| AAV8 | 0.016 | 0.133 | 0.093 | 0.133 | 0.877 | - | - | - | - |
| AAV9 | 0.010 | 0.043 | 0.010 | 0.010 | 0.671 | 1.000 | - | - | - |
| AAVrh8 | 0.010 | 0.360 | 0.026 | 0.220 | 1.000 | 0.877 | 0.697 | - | - |
| AAVrh10 | 0.010 | 0.093 | 0.010 | 0.026 | 1.000 | 0.623 | 0.697 | 0.877 | - |
| AAVrh74 | 0.010 | 0.010 | 0.010 | 0.010 | 0.148 | 0.537 | 0.537 | 0.537 | 0.065 |

Supplemental Table S4 - Wilcoxon Rank Tests - Biodistribution Female

**Eye**

| | AAV3b | AAV4 | AAV5 | AAV6 | AAV7 | AAV8 | AAV9 | AAVrh8 | AAVrh10 |
|---|---|---|---|---|---|---|---|---|---|
| AAV4 | 0.370 | - | - | - | - | - | - | - | - |
| AAV5 | 0.117 | 0.850 | - | - | - | - | - | - | - |
| AAV6 | 0.937 | 0.463 | 0.117 | - | - | - | - | - | - |
| AAV7 | 0.014 | 0.937 | 0.117 | 0.014 | - | - | - | - | - |
| AAV8 | 0.014 | 0.937 | 0.091 | 0.014 | 0.370 | - | - | - | - |
| AAV9 | 0.052 | 0.937 | 0.463 | 0.032 | 0.537 | 0.337 | - | - | - |
| AAVrh8 | 0.032 | 0.937 | 0.370 | 0.019 | 0.850 | 0.370 | 0.937 | - | - |
| AAVrh10 | 0.139 | 0.937 | 0.937 | 0.109 | 0.370 | 0.337 | 0.757 | 0.757 | - |
| AAVrh74 | 0.014 | 0.370 | 0.019 | 0.014 | 0.019 | 0.337 | 0.014 | 0.109 | 0.083 |

**Gastrocnemius muscle**

| | AAV3b | AAV4 | AAV5 | AAV6 | AAV7 | AAV8 | AAV9 | AAVrh8 | AAVrh10 |
|---|---|---|---|---|---|---|---|---|---|
| AAV4 | 0.368 | - | - | - | - | - | - | - | - |
| AAV5 | 0.102 | 0.643 | - | - | - | - | - | - | - |
| AAV6 | 0.019 | 0.505 | 0.051 | - | - | - | - | - | - |
| AAV7 | 0.019 | 0.062 | 0.019 | 0.019 | - | - | - | - | - |
| AAV8 | 0.019 | 0.077 | 0.030 | 0.030 | 0.435 | - | - | - | - |
| AAV9 | 0.159 | 0.368 | 0.521 | 0.521 | 0.856 | 0.632 | - | - | - |
| AAVrh8 | 0.019 | 0.632 | 0.052 | 0.856 | 0.212 | 0.148 | 0.856 | - | - |
| AAVrh10 | 0.019 | 0.222 | 0.051 | 0.159 | 0.931 | 0.422 | 0.931 | 0.368 | - |
| AAVrh74 | 0.019 | 0.062 | 0.019 | 0.019 | 0.062 | 0.068 | 0.122 | 0.062 | 0.102 |

**Heart**

| | AAV3b | AAV4 | AAV5 | AAV6 | AAV7 | AAV8 | AAV9 | AAVrh8 | AAVrh10 |
|---|---|---|---|---|---|---|---|---|---|
| AAV4 | 0.111 | - | - | - | - | - | - | - | - |
| AAV5 | 0.072 | 0.261 | - | - | - | - | - | - | - |
| AAV6 | 0.065 | 0.749 | 0.370 | - | - | - | - | - | - |
| AAV7 | 0.016 | 0.289 | 0.016 | 0.093 | - | - | - | - | - |
| AAV8 | 0.016 | 0.049 | 0.016 | 0.045 | 0.506 | - | - | - | - |
| AAV9 | 0.182 | 0.435 | 0.435 | 0.435 | 0.749 | 0.959 | - | - | - |
| AAVrh8 | 0.016 | 0.536 | 0.016 | 0.289 | 0.697 | 0.435 | 0.749 | - | - |
| AAVrh10 | 0.016 | 0.168 | 0.016 | 0.065 | 0.856 | 0.749 | 1.000 | 0.590 | - |
| AAVrh74 | 0.016 | 0.016 | 0.016 | 0.016 | 0.045 | 0.238 | 0.370 | 0.045 | 0.238 |

**Kidney**

| | AAV3b | AAV4 | AAV5 | AAV6 | AAV7 | AAV8 | AAV9 | AAVrh8 | AAVrh10 |
|---|---|---|---|---|---|---|---|---|---|
| AAV4 | 0.679 | - | - | - | - | - | - | - | - |
| AAV5 | 0.507 | 0.856 | - | - | - | - | - | - | - |
| AAV6 | 0.767 | 0.507 | 0.081 | - | - | - | - | - | - |
| AAV7 | 0.058 | 0.507 | 0.229 | 0.008 | - | - | - | - | - |
| AAV8 | 0.008 | 0.024 | 0.014 | 0.008 | 0.040 | - | - | - | - |
| AAV9 | 0.058 | 0.229 | 0.279 | 0.008 | 0.679 | 0.590 | - | - | - |
| AAVrh8 | 0.081 | 0.590 | 0.360 | 0.014 | 1.000 | 0.175 | 0.435 | - | - |
| AAVrh10 | 0.081 | 0.435 | 0.279 | 0.008 | 0.856 | 0.279 | 0.959 | 0.767 | - |
| AAVrh74 | 0.008 | 0.008 | 0.008 | 0.008 | 0.008 | 0.024 | 0.008 | 0.008 | 0.058 |

**Large Intestine**

| | AAV3b | AAV4 | AAV5 | AAV6 | AAV7 | AAV8 | AAV9 | AAVrh8 | AAVrh10 |
|---|---|---|---|---|---|---|---|---|---|
| AAV4 | 0.321 | - | - | - | - | - | - | - | - |
| AAV5 | 0.120 | 0.574 | - | - | - | - | - | - | - |
| AAV6 | 0.321 | 0.479 | 0.027 | - | - | - | - | - | - |
| AAV7 | 0.027 | 0.120 | 0.012 | 0.012 | - | - | - | - | - |
| AAV8 | 0.042 | 0.111 | 0.018 | 0.018 | 0.869 | - | - | - | - |
| AAV9 | 0.120 | 1.000 | 0.679 | 0.097 | 0.120 | 0.137 | - | - | - |
| AAVrh8 | 0.095 | 0.959 | 0.175 | 0.012 | 0.140 | 0.137 | 0.787 | - | - |
| AAVrh10 | 0.027 | 0.175 | 0.018 | 0.012 | 0.877 | 0.959 | 0.140 | 0.175 | - |
| AAVrh74 | 0.027 | 0.012 | 0.012 | 0.012 | 0.140 | 0.111 | 0.027 | 0.012 | 0.175 |

**Liver**

| | AAV3b | AAV4 | AAV5 | AAV6 | AAV7 | AAV8 | AAV9 | AAVrh8 | AAVrh10 |
|---|---|---|---|---|---|---|---|---|---|
| AAV4 | 0.732 | - | - | - | - | - | - | - | - |
| AAV5 | 0.004 | 0.004 | - | - | - | - | - | - | - |
| AAV6 | 0.004 | 0.004 | 0.084 | - | - | - | - | - | - |
| AAV7 | 0.006 | 0.006 | 0.006 | 0.006 | - | - | - | - | - |
| AAV8 | 0.004 | 0.004 | 0.004 | 0.004 | 0.006 | - | - | - | - |
| AAV9 | 0.004 | 0.004 | 0.004 | 0.012 | 0.216 | 0.116 | - | - | - |
| AAVrh8 | 0.004 | 0.004 | 0.004 | 0.004 | 0.006 | 0.532 | 0.277 | - | - |
| AAVrh10 | 0.004 | 0.004 | 0.004 | 0.004 | 0.040 | 0.631 | 0.348 | 0.818 | - |
| AAVrh74 | 0.004 | 0.004 | 0.004 | 0.004 | 0.006 | 0.818 | 0.004 | 0.021 | 0.277 |

**Lung**

| | AAV3b | AAV4 | AAV5 | AAV6 | AAV7 | AAV8 | AAV9 | AAVrh8 | AAVrh10 |
|---|---|---|---|---|---|---|---|---|---|
| AAV4 | 0.014 | - | - | - | - | - | - | - | - |
| AAV5 | 0.008 | 0.455 | - | - | - | - | - | - | - |
| AAV6 | 0.033 | 0.150 | 0.008 | - | - | - | - | - | - |
| AAV7 | 0.008 | 0.455 | 0.033 | 0.008 | - | - | - | - | - |
| AAV8 | 0.008 | 0.455 | 0.081 | 0.008 | 0.546 | - | - | - | - |
| AAV9 | 0.008 | 0.455 | 0.108 | 0.008 | 0.767 | 0.937 | - | - | - |
| AAVrh8 | 0.008 | 0.435 | 0.205 | 0.023 | 0.856 | 0.856 | 0.937 | - | - |
| AAVrh10 | 0.008 | 0.455 | 0.360 | 0.008 | 0.053 | 0.108 | 0.435 | 0.455 | - |
| AAVrh74 | 0.008 | 0.455 | 0.108 | 0.008 | 0.023 | 0.023 | 0.033 | 0.033 | 0.108 |

**Lymph nodes**

| | AAV3b | AAV4 | AAV5 | AAV6 | AAV7 | AAV8 | AAV9 | AAVrh8 | AAVrh10 |
|---|---|---|---|---|---|---|---|---|---|
| AAV4 | 0.633 | - | - | - | - | - | - | - | - |
| AAV5 | 0.959 | 0.633 | - | - | - | - | - | - | - |
| AAV6 | 0.071 | 0.463 | 0.049 | - | - | - | - | - | - |
| AAV7 | 0.944 | 0.633 | 0.463 | 0.071 | - | - | - | - | - |
| AAV8 | 0.297 | 0.944 | 0.090 | 0.049 | 0.633 | - | - | - | - |
| AAV9 | 0.233 | 0.944 | 0.195 | 0.049 | 0.463 | 0.944 | - | - | - |
| AAVrh8 | 0.297 | 0.931 | 0.097 | 0.054 | 0.463 | 0.805 | 0.959 | - | - |
| AAVrh10 | 0.071 | 0.944 | 0.054 | 0.071 | 0.054 | 0.885 | 0.959 | 0.944 | - |
| AAVrh74 | 0.233 | 0.944 | 0.054 | 0.049 | 0.233 | 1.000 | 0.959 | 0.959 | 0.739 |

Supplemental Table S4 - Wilcoxon Rank Tests - Biodistribution Female

**Pancreas**

| | AAV3b | AAV4 | AAV5 | AAV6 | AAV7 | AAV8 | AAV9 | AAVrh8 | AAVrh10 |
|---|---|---|---|---|---|---|---|---|---|
| AAV4 | 0.176 | - | - | - | - | - | - | - | - |
| AAV5 | 0.261 | 0.220 | - | - | - | - | - | - | - |
| AAV6 | 0.387 | 0.220 | 0.856 | - | - | - | - | - | - |
| AAV7 | 0.028 | 0.261 | 0.028 | 0.103 | - | - | - | - | - |
| AAV8 | 0.012 | 0.937 | 0.012 | 0.012 | 0.069 | - | - | - | - |
| AAV9 | 0.146 | 0.400 | 0.631 | 0.455 | 0.318 | 0.069 | - | - | - |
| AAVrh8 | 0.146 | 0.220 | 0.318 | 0.261 | 0.318 | 0.028 | 0.937 | - | - |
| AAVrh10 | 0.018 | 0.589 | 0.018 | 0.018 | 0.261 | 0.455 | 0.220 | 0.190 | - |
| AAVrh74 | 0.012 | 0.482 | 0.012 | 0.012 | 0.012 | 0.387 | 0.069 | 0.012 | 0.220 |

**Skin**

| | AAV3b | AAV4 | AAV5 | AAV6 | AAV7 | AAV8 | AAV9 | AAVrh8 | AAVrh10 |
|---|---|---|---|---|---|---|---|---|---|
| AAV4 | 0.464 | - | - | - | - | - | - | - | - |
| AAV5 | 0.023 | 0.030 | - | - | - | - | - | - | - |
| AAV6 | 0.023 | 0.367 | 0.376 | - | - | - | - | - | - |
| AAV7 | 0.023 | 0.023 | 0.188 | 0.087 | - | - | - | - | - |
| AAV8 | 0.023 | 0.030 | 0.023 | 0.023 | 0.575 | - | - | - | - |
| AAV9 | 0.023 | 0.023 | 0.023 | 0.023 | 0.245 | 0.470 | - | - | - |
| AAVrh8 | 0.023 | 0.066 | 0.829 | 0.188 | 0.245 | 0.245 | 0.030 | - | - |
| AAVrh10 | 0.023 | 0.023 | 0.380 | 0.188 | 0.376 | 0.055 | 0.030 | 0.959 | - |
| AAVrh74 | 0.023 | 0.030 | 0.023 | 0.023 | 0.380 | 0.294 | 1.000 | 0.023 | 0.023 |

**Small Intestine**

| | AAV3b | AAV4 | AAV5 | AAV6 | AAV7 | AAV8 | AAV9 | AAVrh8 | AAVrh10 |
|---|---|---|---|---|---|---|---|---|---|
| AAV4 | 0.159 | - | - | - | - | - | - | - | - |
| AAV5 | 0.159 | 0.195 | - | - | - | - | - | - | - |
| AAV6 | 0.159 | 0.219 | 0.952 | - | - | - | - | - | - |
| AAV7 | 0.200 | 0.947 | 0.048 | 0.065 | - | - | - | - | - |
| AAV8 | 0.159 | 0.411 | 0.048 | 0.048 | 0.722 | - | - | - | - |
| AAV9 | 0.159 | 1.000 | 0.095 | 0.095 | 0.836 | 0.318 | - | - | - |
| AAVrh8 | 0.159 | 0.159 | 0.048 | 0.048 | 0.179 | 0.521 | 0.159 | - | - |
| AAVrh10 | 0.159 | 0.065 | 0.048 | 0.048 | 0.095 | 0.250 | 0.095 | 0.777 | - |
| AAVrh74 | 0.159 | 0.159 | 0.048 | 0.048 | 0.260 | 0.411 | 0.146 | 0.901 | 0.777 |

**Spleen**

| | AAV3b | AAV4 | AAV5 | AAV6 | AAV7 | AAV8 | AAV9 | AAVrh8 | AAVrh10 |
|---|---|---|---|---|---|---|---|---|---|
| AAV4 | 0.246 | - | - | - | - | - | - | - | - |
| AAV5 | 0.018 | 0.784 | - | - | - | - | - | - | - |
| AAV6 | 0.959 | 0.246 | 0.018 | - | - | - | - | - | - |
| AAV7 | 0.784 | 0.271 | 0.018 | 0.529 | - | - | - | - | - |
| AAV8 | 0.018 | 0.710 | 0.073 | 0.018 | 0.028 | - | - | - | - |
| AAV9 | 0.028 | 0.901 | 0.959 | 0.018 | 0.028 | 0.529 | - | - | - |
| AAVrh8 | 0.661 | 0.307 | 0.018 | 0.661 | 0.784 | 0.146 | 0.052 | - | - |
| AAVrh10 | 0.103 | 0.784 | 0.611 | 0.146 | 0.246 | 0.898 | 0.643 | 0.248 | - |
| AAVrh74 | 0.018 | 1.000 | 0.898 | 0.018 | 0.018 | 0.103 | 0.898 | 0.018 | 0.661 |

Supplemental Table S4 - Wilcoxon Rank Tests - Biodistribution Female

**Stomach**

| | AAV3b | AAV4 | AAV5 | AAV6 | AAV7 | AAV8 | AAV9 | AAVrh8 | AAVrh10 |
|---|---|---|---|---|---|---|---|---|---|
| AAV4 | 0.328 | - | - | - | - | - | - | - | - |
| AAV5 | 0.005 | 0.479 | - | - | - | - | - | - | - |
| AAV6 | 0.016 | 0.479 | 0.856 | - | - | - | - | - | - |
| AAV7 | 0.005 | 0.479 | 0.005 | 0.008 | - | - | - | - | - |
| AAV8 | 0.005 | 0.192 | 0.005 | 0.005 | 0.104 | - | - | - | - |
| AAV9 | 0.005 | 0.192 | 0.005 | 0.005 | 0.253 | 0.959 | - | - | - |
| AAVrh8 | 0.005 | 0.856 | 0.008 | 0.008 | 0.767 | 0.679 | 0.767 | - | - |
| AAVrh10 | 0.005 | 0.192 | 0.005 | 0.008 | 0.104 | 0.679 | 1.000 | 0.479 | - |
| AAVrh74 | 0.005 | 0.005 | 0.005 | 0.005 | 0.005 | 0.005 | 0.005 | 0.071 | 0.005 |

**Thymus**

| | AAV3b | AAV4 | AAV5 | AAV6 | AAV7 | AAV8 | AAV9 | AAVrh8 | AAVrh10 |
|---|---|---|---|---|---|---|---|---|---|
| AAV4 | 0.506 | - | - | - | - | - | - | - | - |
| AAV5 | 0.043 | 0.937 | - | - | - | - | - | - | - |
| AAV6 | 0.449 | 0.506 | 0.043 | - | - | - | - | - | - |
| AAV7 | 0.018 | 0.449 | 0.506 | 0.030 | - | - | - | - | - |
| AAV8 | 0.018 | 0.103 | 0.229 | 0.043 | 0.449 | - | - | - | - |
| AAV9 | 0.133 | 0.787 | 0.697 | 0.182 | 0.937 | 0.606 | - | - | - |
| AAVrh8 | 0.133 | 0.787 | 0.856 | 0.229 | 0.856 | 0.386 | 0.856 | - | - |
| AAVrh10 | 0.011 | 0.011 | 0.030 | 0.011 | 0.133 | 0.697 | 0.506 | 0.133 | - |
| AAVrh74 | 0.011 | 0.011 | 0.011 | 0.011 | 0.011 | 0.229 | 0.299 | 0.011 | 0.103 |

**Ovary**

| | AAV3b | AAV4 | AAV5 | AAV6 | AAV7 | AAV8 | AAV9 | AAVrh8 | AAVrh10 |
|---|---|---|---|---|---|---|---|---|---|
| AAV4 | 0.090 | - | - | - | - | - | - | - | - |
| AAV5 | 0.008 | 0.508 | - | - | - | - | - | - | - |
| AAV6 | 0.270 | 0.137 | 0.008 | - | - | - | - | - | - |
| AAV7 | 0.008 | 0.090 | 0.009 | 0.008 | - | - | - | - | - |
| AAV8 | 0.008 | 0.009 | 0.008 | 0.008 | 0.422 | - | - | - | - |
| AAV9 | 0.008 | 0.358 | 0.492 | 0.009 | 1.000 | 0.662 | - | - | - |
| AAVrh8 | 0.009 | 0.422 | 1.000 | 0.009 | 0.035 | 0.009 | 0.435 | - | - |
| AAVrh10 | 0.008 | 0.508 | 0.150 | 0.008 | 0.492 | 0.077 | 0.767 | 0.195 | - |
| AAVrh74 | 0.008 | 0.009 | 0.008 | 0.008 | 0.559 | 0.856 | 0.856 | 0.009 | 0.077 |

**Uterus**

| | AAV3b | AAV4 | AAV5 | AAV6 | AAV7 | AAV8 | AAV9 | AAVrh8 | AAVrh10 |
|---|---|---|---|---|---|---|---|---|---|
| AAV4 | 0.253 | - | - | - | - | - | - | - | - |
| AAV5 | 0.024 | 0.914 | - | - | - | - | - | - | - |
| AAV6 | 0.376 | 0.253 | 0.056 | - | - | - | - | - | - |
| AAV7 | 0.024 | 0.040 | 0.041 | 0.024 | - | - | - | - | - |
| AAV8 | 0.024 | 0.029 | 0.024 | 0.024 | 0.253 | - | - | - | - |
| AAV9 | 0.024 | 0.253 | 0.253 | 0.024 | 0.327 | 0.040 | - | - | - |
| AAVrh8 | 0.029 | 0.056 | 0.120 | 0.029 | 0.610 | 0.029 | 0.638 | - | - |
| AAVrh10 | 0.040 | 0.638 | 0.467 | 0.056 | 0.327 | 0.220 | 0.715 | 0.536 | - |
| AAVrh74 | 0.029 | 0.056 | 0.029 | 0.029 | 0.357 | 0.536 | 0.198 | 0.107 | 0.331 |

Supplemental Table S5: Serotype with the highest transduction efficiency (GC/μg) DNA for each organ

| | Adrenal glands | Bladder | Brain | Diaphragm | Eye | Gastroc muscle | Heart | Kidney | Large Intestine |
|---|---|---|---|---|---|---|---|---|---|
| Male | AAVrh10 | AAV8 | AAV4 | AAVrh10 | AAVrh10 | AAVrh10 | AAVrh10 | AAVrh10 | AAVrh10 |
| Female | AAV8 | AAV8 | AAVrh74 | AAV8 | AAVrh74 | AAVrh74 | AAVrh74 | AAVrh74 | AAVrh74 |

| | Liver | Lung | Lymph nodes | Pancreas | Skin | Small intestine | Spleen | Stomach | Thymus |
|---|---|---|---|---|---|---|---|---|---|
| Male | AAVrh10 | AAV4 | AAV4 | AAVrh10 | AAV9 | AAVrh10 | AAVrh10 | AAVrh10 | AAVrh10 |
| Female | AAVrh74 | AAV4 | AAV9 | AAV8 | AAV9 | AAVrh74 | AAV4 | AAVrh74 | AAV8 |

| | Testes | Epididymis | Ovary | Uterus |
|---|---|---|---|---|
| Male | AAVrh10 | AAV8 | | |
| Female | | | AAV7 | AAV8 |

Supplemental Table S6 - Wilcoxon Rank Tests - Functional Transduction Male

**Adrenals**

| | AAV3b | AAV4 | AAV5 | AAV6 | AAV7 | AAV8 | AAV9 | AAVrh8 | AAVrh10 |
|---|---|---|---|---|---|---|---|---|---|
| AAV4 | 0.753 | - | - | - | - | - | - | - | - |
| AAV5 | 0.406 | 0.406 | - | - | - | - | - | - | - |
| AAV6 | 0.046 | 0.046 | 0.046 | - | - | - | - | - | - |
| AAV7 | 0.046 | 0.046 | 0.046 | 0.046 | - | - | - | - | - |
| AAV8 | 0.046 | 0.046 | 0.086 | 0.886 | 0.046 | - | - | - | - |
| AAV9 | 0.046 | 0.046 | 0.046 | 0.265 | 0.046 | 0.886 | - | - | - |
| AAVrh8 | 0.046 | 0.046 | 0.046 | 0.046 | 0.046 | 0.046 | 0.046 | - | - |
| AAVrh10 | 0.046 | 0.046 | 0.046 | 0.086 | 0.046 | 0.161 | 0.406 | 0.753 | - |
| AAVrh74 | 0.046 | 0.046 | 0.046 | 0.265 | 0.161 | 0.886 | 0.886 | 0.406 | 0.560 |

**Bladder**

| | AAV3b | AAV4 | AAV5 | AAV6 | AAV7 | AAV8 | AAV9 | AAVrh8 | AAVrh10 |
|---|---|---|---|---|---|---|---|---|---|
| AAV4 | 0.092 | - | - | - | - | - | - | - | - |
| AAV5 | 0.315 | 0.166 | - | - | - | - | - | - | - |
| AAV6 | 0.068 | 0.166 | 0.068 | - | - | - | - | - | - |
| AAV7 | 0.068 | 0.092 | 0.068 | 0.068 | - | - | - | - | - |
| AAV8 | 0.068 | 0.166 | 0.068 | 0.396 | 0.396 | - | - | - | - |
| AAV9 | 0.068 | 0.092 | 0.068 | 0.092 | 0.092 | 0.533 | - | - | - |
| AAVrh8 | 0.068 | 0.092 | 0.068 | 0.068 | 0.265 | 0.927 | 0.396 | - | - |
| AAVrh10 | 0.068 | 0.092 | 0.068 | 0.068 | 1.000 | 0.265 | 0.068 | 0.265 | - |
| AAVrh74 | 0.068 | 0.092 | 0.068 | 0.068 | 0.927 | 0.396 | 0.092 | 0.533 | 1.000 |

**Brain**

| | AAV3b | AAV4 | AAV5 | AAV6 | AAV7 | AAV8 | AAV9 | AAVrh8 | AAVrh10 |
|---|---|---|---|---|---|---|---|---|---|
| AAV4 | 0.083 | - | - | - | - | - | - | - | - |
| AAV5 | 0.281 | 0.575 | - | - | - | - | - | - | - |
| AAV6 | 0.083 | 0.281 | 0.190 | - | - | - | - | - | - |
| AAV7 | 0.083 | 0.083 | 0.083 | 0.735 | - | - | - | - | - |
| AAV8 | 0.083 | 0.575 | 0.575 | 0.112 | 0.083 | - | - | - | - |
| AAV9 | 0.083 | 0.281 | 0.190 | 0.735 | 0.112 | 0.112 | - | - | - |
| AAVrh8 | 0.083 | 0.281 | 0.112 | 0.906 | 0.735 | 0.112 | 0.441 | - | - |
| AAVrh10 | 0.083 | 0.083 | 0.112 | 0.906 | 0.281 | 0.083 | 0.441 | 1.000 | - |
| AAVrh74 | 0.083 | 0.083 | 0.083 | 0.190 | 0.735 | 0.083 | 0.112 | 0.190 | 0.441 |

**Diaphragm**

| | AAV3b | AAV4 | AAV5 | AAV6 | AAV7 | AAV8 | AAV9 | AAVrh8 | AAVrh10 |
|---|---|---|---|---|---|---|---|---|---|
| AAV4 | 0.076 | - | - | - | - | - | - | - | - |
| AAV5 | 0.514 | 0.129 | - | - | - | - | - | - | - |
| AAV6 | 0.076 | 0.076 | 0.076 | - | - | - | - | - | - |
| AAV7 | 0.375 | 0.514 | 0.129 | 0.643 | - | - | - | - | - |
| AAV8 | 0.076 | 0.375 | 0.129 | 0.927 | 0.514 | - | - | - | - |
| AAV9 | 0.076 | 0.076 | 0.076 | 0.857 | 0.643 | 0.643 | - | - | - |
| AAVrh8 | 0.076 | 0.076 | 0.076 | 0.643 | 1.000 | 0.514 | 0.927 | - | - |
| AAVrh10 | 0.076 | 0.076 | 0.076 | 0.375 | 0.927 | 0.245 | 0.857 | 0.927 | - |
| AAVrh74 | 0.076 | 0.076 | 0.076 | 0.514 | 0.927 | 0.514 | 0.927 | 1.000 | 0.927 |

Supplemental Table S6 - Wilcoxon Rank Tests - Functional Transduction Male

**Eye**

| | AAV3b | AAV4 | AAV5 | AAV6 | AAV7 | AAV8 | AAV9 | AAVrh8 | AAVrh10 |
|---|---|---|---|---|---|---|---|---|---|
| AAV4 | 0.066 | - | - | - | - | - | - | - | - |
| AAV5 | 0.184 | 0.107 | - | - | - | - | - | - | - |
| AAV6 | 0.066 | 0.429 | 0.107 | - | - | - | - | - | - |
| AAV7 | 0.066 | 0.066 | 0.066 | 0.184 | - | - | - | - | - |
| AAV8 | 0.066 | 1.000 | 0.290 | 0.290 | 0.066 | - | - | - | - |
| AAV9 | 0.066 | 0.066 | 0.066 | 0.771 | 0.184 | 0.107 | - | - | - |
| AAVrh8 | 0.066 | 0.107 | 0.066 | 0.771 | 0.290 | 0.184 | 1.000 | - | - |
| AAVrh10 | 0.066 | 0.066 | 0.066 | 0.429 | 0.429 | 0.066 | 0.429 | 1.000 | - |
| AAVrh74 | 0.066 | 0.066 | 0.066 | 0.429 | 0.771 | 0.066 | 0.591 | 0.949 | 0.949 |

**Gastrocnemius muscle**

| | AAV3b | AAV4 | AAV5 | AAV6 | AAV7 | AAV8 | AAV9 | AAVrh8 | AAVrh10 |
|---|---|---|---|---|---|---|---|---|---|
| AAV4 | 0.468 | - | - | - | - | - | - | - | - |
| AAV5 | 0.753 | 0.906 | - | - | - | - | - | - | - |
| AAV6 | 0.076 | 0.076 | 0.076 | - | - | - | - | - | - |
| AAV7 | 0.122 | 0.122 | 0.224 | 0.343 | - | - | - | - | - |
| AAV8 | 0.076 | 0.076 | 0.122 | 0.333 | 0.906 | - | - | - | - |
| AAV9 | 0.076 | 0.076 | 0.076 | 0.333 | 0.725 | 0.906 | - | - | - |
| AAVrh8 | 0.076 | 0.076 | 0.076 | 0.575 | 0.343 | 0.575 | 0.468 | - | - |
| AAVrh10 | 0.076 | 0.076 | 0.076 | 1.000 | 0.122 | 0.224 | 0.333 | 0.333 | - |
| AAVrh74 | 0.076 | 0.076 | 0.076 | 0.575 | 0.343 | 0.575 | 0.468 | 0.753 | 0.575 |

**Heart**

| | AAV3b | AAV4 | AAV5 | AAV6 | AAV7 | AAV8 | AAV9 | AAVrh8 | AAVrh10 |
|---|---|---|---|---|---|---|---|---|---|
| AAV4 | 0.265 | - | - | - | - | - | - | - | - |
| AAV5 | 0.972 | 0.265 | - | - | - | - | - | - | - |
| AAV6 | 0.058 | 0.099 | 0.058 | - | - | - | - | - | - |
| AAV7 | 0.058 | 0.058 | 0.058 | 0.058 | - | - | - | - | - |
| AAV8 | 0.099 | 0.265 | 0.058 | 1.000 | 0.265 | - | - | - | - |
| AAV9 | 0.058 | 0.058 | 0.058 | 0.177 | 1.000 | 0.265 | - | - | - |
| AAVrh8 | 0.058 | 0.058 | 0.058 | 0.834 | 0.058 | 1.000 | 0.177 | - | - |
| AAVrh10 | 0.058 | 0.058 | 0.058 | 0.099 | 0.972 | 0.441 | 0.834 | 0.099 | - |
| AAVrh74 | 0.058 | 0.058 | 0.058 | 1.000 | 0.058 | 0.972 | 0.177 | 0.972 | 0.058 |

**Kidney**

| | AAV3b | AAV4 | AAV5 | AAV6 | AAV7 | AAV8 | AAV9 | AAVrh8 | AAVrh10 |
|---|---|---|---|---|---|---|---|---|---|
| AAV4 | 0.086 | - | - | - | - | - | - | - | - |
| AAV5 | 0.086 | 0.482 | - | - | - | - | - | - | - |
| AAV6 | 0.086 | 0.234 | 0.482 | - | - | - | - | - | - |
| AAV7 | 0.086 | 0.086 | 0.333 | 0.906 | - | - | - | - | - |
| AAV8 | 0.086 | 0.333 | 0.753 | 0.333 | 0.135 | - | - | - | - |
| AAV9 | 0.086 | 0.753 | 0.753 | 0.333 | 0.234 | 0.643 | - | - | - |
| AAVrh8 | 0.086 | 0.086 | 0.135 | 0.482 | 0.482 | 0.086 | 0.086 | - | - |
| AAVrh10 | 0.086 | 0.086 | 0.333 | 0.906 | 0.906 | 0.135 | 0.135 | 0.753 | - |
| AAVrh74 | 0.086 | 0.086 | 0.753 | 1.000 | 0.753 | 0.643 | 0.234 | 0.482 | 0.753 |

**Large intestine**

| | AAV3b | AAV4 | AAV5 | AAV6 | AAV7 | AAV8 | AAV9 | AAVrh8 | AAVrh10 |
|---|---|---|---|---|---|---|---|---|---|
| AAV4 | 0.068 | - | - | - | - | - | - | - | - |
| AAV5 | 0.103 | 0.735 | - | - | - | - | - | - | - |
| AAV6 | 0.068 | 0.575 | 0.735 | - | - | - | - | - | - |
| AAV7 | 0.068 | 0.068 | 0.068 | 0.068 | - | - | - | - | - |
| AAV8 | 0.098 | 0.171 | 0.286 | 0.286 | 0.171 | - | - | - | - |
| AAV9 | 0.068 | 0.281 | 0.171 | 0.103 | 0.103 | 0.897 | - | - | - |
| AAVrh8 | 0.068 | 0.068 | 0.068 | 0.068 | 0.417 | 0.286 | 0.171 | - | - |
| AAVrh10 | 0.068 | 0.068 | 0.068 | 0.068 | 0.735 | 0.103 | 0.103 | 0.281 | - |
| AAVrh74 | 0.068 | 0.068 | 0.068 | 0.068 | 0.906 | 0.286 | 0.171 | 0.735 | 1.000 |

**Liver**

| | AAV3b | AAV4 | AAV5 | AAV6 | AAV7 | AAV8 | AAV9 | AAVrh8 | AAVrh10 |
|---|---|---|---|---|---|---|---|---|---|
| AAV4 | 0.151 | - | - | - | - | - | - | - | - |
| AAV5 | 0.643 | 0.643 | - | - | - | - | - | - | - |
| AAV6 | 0.088 | 0.088 | 0.949 | - | - | - | - | - | - |
| AAV7 | 0.088 | 0.088 | 0.949 | 0.643 | - | - | - | - | - |
| AAV8 | 0.088 | 0.088 | 0.949 | 1.000 | 0.691 | - | - | - | - |
| AAV9 | 0.088 | 0.088 | 0.949 | 0.949 | 0.643 | 0.949 | - | - | - |
| AAVrh8 | 0.088 | 0.088 | 0.643 | 0.949 | 0.151 | 0.949 | 0.949 | - | - |
| AAVrh10 | 0.088 | 0.088 | 0.949 | 0.949 | 0.643 | 0.949 | 0.949 | 0.949 | - |
| AAVrh74 | 0.088 | 0.088 | 0.643 | 1.000 | 0.088 | 0.949 | 0.780 | 1.000 | 0.780 |

**Lung**

| | AAV3b | AAV4 | AAV5 | AAV6 | AAV7 | AAV8 | AAV9 | AAVrh8 | AAVrh10 |
|---|---|---|---|---|---|---|---|---|---|
| AAV4 | 0.110 | - | - | - | - | - | - | - | - |
| AAV5 | 0.260 | 0.830 | - | - | - | - | - | - | - |
| AAV6 | 0.110 | 0.110 | 0.680 | - | - | - | - | - | - |
| AAV7 | 0.110 | 0.110 | 0.530 | 0.680 | - | - | - | - | - |
| AAV8 | 0.110 | 1.000 | 0.830 | 0.530 | 0.530 | - | - | - | - |
| AAV9 | 0.140 | 0.140 | 1.000 | 0.140 | 0.140 | 0.830 | - | - | - |
| AAVrh8 | 0.110 | 0.830 | 1.000 | 0.530 | 0.260 | 0.970 | 1.000 | - | - |
| AAVrh10 | 0.110 | 0.110 | 0.140 | 0.110 | 0.110 | 0.530 | 0.140 | 0.380 | - |
| AAVrh74 | 0.110 | 0.970 | 0.830 | 0.380 | 0.380 | 0.970 | 0.600 | 0.970 | 0.380 |

**Lymph nodes**

| | AAV3b | AAV4 | AAV5 | AAV6 | AAV7 | AAV8 | AAV9 | AAVrh8 | AAVrh10 |
|---|---|---|---|---|---|---|---|---|---|
| AAV4 | 0.066 | - | - | - | - | - | - | - | - |
| AAV5 | 0.116 | 0.906 | - | - | - | - | - | - | - |
| AAV6 | 0.066 | 0.116 | 0.177 | - | - | - | - | - | - |
| AAV7 | 0.066 | 0.066 | 0.066 | 0.177 | - | - | - | - | - |
| AAV8 | 0.066 | 0.066 | 0.116 | 1.000 | 0.066 | - | - | - | - |
| AAV9 | 0.066 | 0.066 | 0.066 | 0.591 | 0.177 | 0.735 | - | - | - |
| AAVrh8 | 0.066 | 0.066 | 0.066 | 0.300 | 0.735 | 0.454 | 0.591 | - | - |
| AAVrh10 | 0.066 | 0.066 | 0.066 | 0.177 | 0.735 | 0.177 | 0.177 | 0.591 | - |
| AAVrh74 | 0.066 | 0.066 | 0.066 | 0.454 | 0.735 | 0.454 | 0.735 | 0.906 | 0.454 |

Supplemental Table S6 - Wilcoxon Rank Tests - Functional Transduction Male

**Pancreas**

| | AAV3b | AAV4 | AAV5 | AAV6 | AAV7 | AAV8 | AAV9 | AAVrh8 | AAVrh10 |
|---|---|---|---|---|---|---|---|---|---|
| AAV4 | 0.092 | - | - | - | - | - | - | - | - |
| AAV5 | 0.243 | 1.000 | - | - | - | - | - | - | - |
| AAV6 | 0.092 | 0.735 | 0.927 | - | - | - | - | - | - |
| AAV7 | 0.070 | 0.070 | 0.070 | 0.070 | - | - | - | - | - |
| AAV8 | 0.070 | 0.092 | 0.151 | 0.092 | 0.243 | - | - | - | - |
| AAV9 | 0.092 | 0.092 | 0.151 | 0.151 | 0.151 | 0.462 | - | - | - |
| AAVrh8 | 0.070 | 0.070 | 0.070 | 0.070 | 0.406 | 0.243 | 0.092 | - | - |
| AAVrh10 | 0.070 | 0.070 | 0.070 | 0.070 | 0.546 | 0.070 | 0.092 | 0.151 | - |
| AAVrh74 | 0.070 | 0.070 | 0.070 | 0.070 | 0.735 | 0.070 | 0.092 | 0.092 | 1.000 |

**Skin**

| | AAV3b | AAV4 | AAV5 | AAV6 | AAV7 | AAV8 | AAV9 | AAVrh8 | AAVrh10 |
|---|---|---|---|---|---|---|---|---|---|
| AAV4 | 0.064 | - | - | - | - | - | - | - | - |
| AAV5 | 0.560 | 0.906 | - | - | - | - | - | - | - |
| AAV6 | 0.064 | 0.090 | 0.190 | - | - | - | - | - | - |
| AAV7 | 0.064 | 0.064 | 0.099 | 0.560 | - | - | - | - | - |
| AAV8 | 0.064 | 0.064 | 0.099 | 0.560 | 0.735 | - | - | - | - |
| AAV9 | 0.064 | 0.064 | 0.064 | 0.290 | 0.906 | 0.064 | - | - | - |
| AAVrh8 | 0.064 | 0.064 | 0.064 | 0.099 | 0.290 | 0.064 | 0.429 | - | - |
| AAVrh10 | 0.064 | 0.064 | 0.099 | 0.429 | 1.000 | 0.735 | 0.429 | 0.290 | - |
| AAVrh74 | 0.064 | 0.064 | 0.064 | 0.099 | 0.290 | 0.064 | 0.429 | 0.735 | 0.429 |

**Small intestine**

| | AAV3b | AAV4 | AAV5 | AAV6 | AAV7 | AAV8 | AAV9 | AAVrh8 | AAVrh10 |
|---|---|---|---|---|---|---|---|---|---|
| AAV4 | 0.257 | - | - | - | - | - | - | - | - |
| AAV5 | 0.406 | 0.257 | - | - | - | - | - | - | - |
| AAV6 | 0.406 | 0.257 | 0.906 | - | - | - | - | - | - |
| AAV7 | 0.058 | 0.058 | 0.058 | 0.058 | - | - | - | - | - |
| AAV8 | 0.092 | 0.406 | 0.171 | 0.092 | 0.092 | - | - | - | - |
| AAV9 | 0.058 | 0.058 | 0.058 | 0.058 | 0.257 | 0.171 | - | - | - |
| AAVrh8 | 0.058 | 0.058 | 0.058 | 0.058 | 0.560 | 0.092 | 0.718 | - | - |
| AAVrh10 | 0.058 | 0.058 | 0.058 | 0.058 | 0.718 | 0.058 | 0.092 | 0.718 | - |
| AAVrh74 | 0.058 | 0.058 | 0.058 | 0.058 | 1.000 | 0.058 | 0.092 | 0.257 | 0.718 |

**Spleen**

| | AAV3b | AAV4 | AAV5 | AAV6 | AAV7 | AAV8 | AAV9 | AAVrh8 | AAVrh10 |
|---|---|---|---|---|---|---|---|---|---|
| AAV4 | 0.071 | - | - | - | - | - | - | - | - |
| AAV5 | 0.071 | 0.468 | - | - | - | - | - | - | - |
| AAV6 | 0.071 | 0.071 | 0.468 | - | - | - | - | - | - |
| AAV7 | 0.071 | 0.071 | 0.468 | 0.468 | - | - | - | - | - |
| AAV8 | 0.071 | 0.071 | 0.906 | 0.071 | 0.375 | - | - | - | - |
| AAV9 | 0.071 | 0.071 | 0.753 | 0.375 | 0.753 | 0.906 | - | - | - |
| AAVrh8 | 0.071 | 0.071 | 0.468 | 1.000 | 0.753 | 0.468 | 0.468 | - | - |
| AAVrh10 | 0.071 | 0.071 | 0.468 | 0.753 | 0.624 | 0.135 | 0.468 | 0.753 | - |
| AAVrh74 | 0.071 | 0.071 | 0.375 | 0.753 | 0.375 | 0.071 | 0.375 | 0.906 | 0.624 |

Supplemental Table S6 - Wilcoxon Rank Tests - Functional Transduction Male

**Stomach**

| | AAV3b | AAV4 | AAV5 | AAV6 | AAV7 | AAV8 | AAV9 | AAVrh8 | AAVrh10 |
|---|---|---|---|---|---|---|---|---|---|
| AAV4 | 0.350 | - | - | - | - | - | - | - | - |
| AAV5 | 0.240 | 0.930 | - | - | - | - | - | - | - |
| AAV6 | 0.350 | 0.770 | 1.000 | - | - | - | - | - | - |
| AAV7 | 0.150 | 0.150 | 0.240 | 0.150 | - | - | - | - | - |
| AAV8 | 0.150 | 0.640 | 0.530 | 0.530 | 1.000 | - | - | - | - |
| AAV9 | 0.150 | 0.350 | 0.240 | 0.240 | 0.930 | 0.930 | - | - | - |
| AAVrh8 | 0.150 | 0.150 | 0.150 | 0.150 | 0.770 | 0.640 | 0.640 | - | - |
| AAVrh10 | 0.150 | 0.150 | 0.150 | 0.150 | 0.740 | 0.740 | 0.600 | 0.740 | - |
| AAVrh74 | 0.150 | 0.150 | 0.150 | 0.150 | 0.350 | 0.530 | 0.350 | 0.640 | 0.740 |

**Thymus**

| | AAV3b | AAV4 | AAV5 | AAV6 | AAV7 | AAV8 | AAV9 | AAVrh8 | AAVrh10 |
|---|---|---|---|---|---|---|---|---|---|
| AAV4 | 0.273 | - | - | - | - | - | - | - | - |
| AAV5 | 0.559 | 1.000 | - | - | - | - | - | - | - |
| AAV6 | 0.273 | 0.560 | 0.560 | - | - | - | - | - | - |
| AAV7 | 0.099 | 0.099 | 0.099 | 0.099 | - | - | - | - | - |
| AAV8 | 0.099 | 0.099 | 0.184 | 0.184 | 0.273 | - | - | - | - |
| AAV9 | 0.070 | 0.070 | 0.070 | 0.070 | 0.099 | 0.897 | - | - | - |
| AAVrh8 | 0.070 | 0.070 | 0.070 | 0.070 | 0.897 | 0.294 | 0.070 | - | - |
| AAVrh10 | 0.070 | 0.070 | 0.070 | 0.070 | 0.707 | 0.500 | 0.070 | 0.273 | - |
| AAVrh74 | 0.070 | 0.070 | 0.070 | 0.070 | 0.897 | 0.294 | 0.070 | 0.906 | 0.273 |

**Testes**

| | AAV3b | AAV4 | AAV5 | AAV6 | AAV7 | AAV8 | AAV9 | AAVrh8 | AAVrh10 |
|---|---|---|---|---|---|---|---|---|---|
| AAV4 | 0.058 | - | - | - | - | - | - | - | - |
| AAV5 | 0.099 | 1.000 | - | - | - | - | - | - | - |
| AAV6 | 0.058 | 0.058 | 0.643 | - | - | - | - | - | - |
| AAV7 | 0.058 | 0.058 | 0.058 | 0.058 | - | - | - | - | - |
| AAV8 | 0.058 | 0.099 | 0.190 | 0.643 | 1.000 | - | - | - | - |
| AAV9 | 0.058 | 0.058 | 0.099 | 0.235 | 0.482 | 1.000 | - | - | - |
| AAVrh8 | 0.058 | 0.058 | 0.058 | 0.099 | 1.000 | 0.996 | 0.857 | - | - |
| AAVrh10 | 0.058 | 0.058 | 0.058 | 0.058 | 0.058 | 0.996 | 0.482 | 1.000 | - |
| AAVrh74 | 0.058 | 0.058 | 0.058 | 0.058 | 0.996 | 0.996 | 0.310 | 0.857 | 0.482 |

**Epididymis**

| | AAV3b | AAV4 | AAV5 | AAV6 | AAV7 | AAV8 | AAV9 | AAVrh8 | AAVrh10 |
|---|---|---|---|---|---|---|---|---|---|
| AAV4 | 0.135 | - | - | - | - | - | - | - | - |
| AAV5 | 0.206 | 0.906 | - | - | - | - | - | - | - |
| AAV6 | 0.135 | 0.468 | 0.206 | - | - | - | - | - | - |
| AAV7 | 0.099 | 0.099 | 0.099 | 0.468 | - | - | - | - | - |
| AAV8 | 0.135 | 0.683 | 0.468 | 0.753 | 0.135 | - | - | - | - |
| AAV9 | 0.099 | 0.206 | 0.135 | 0.906 | 0.346 | 0.607 | - | - | - |
| AAVrh8 | 0.099 | 0.099 | 0.099 | 0.607 | 0.753 | 0.206 | 0.607 | - | - |
| AAVrh10 | 0.099 | 0.099 | 0.099 | 0.468 | 0.906 | 0.135 | 0.206 | 0.468 | - |
| AAVrh74 | 0.099 | 0.099 | 0.099 | 0.468 | 1.000 | 0.206 | 0.468 | 0.753 | 0.753 |

Supplemental Table S7 - Wilcoxon Rank Tests - Functional Transduction Female

**Adrenals**

| | AAV3b | AAV4 | AAV5 | AAV6 | AAV7 | AAV8 | AAV9 | AAVrh8 | AAVrh10 |
|---|---|---|---|---|---|---|---|---|---|
| AAV4 | 0.198 | - | - | - | - | - | - | - | - |
| AAV5 | 0.198 | 0.061 | - | - | - | - | - | - | - |
| AAV6 | 0.061 | 0.061 | 0.061 | - | - | - | - | - | - |
| AAV7 | 0.117 | 0.061 | 0.551 | 0.643 | - | - | - | - | - |
| AAV8 | 0.061 | 0.061 | 0.061 | 0.791 | 0.791 | - | - | - | - |
| AAV9 | 0.061 | 0.061 | 0.061 | 0.643 | 0.791 | 1.000 | - | - | - |
| AAVrh8 | 0.061 | 0.061 | 0.061 | 0.061 | 0.949 | 0.643 | 0.198 | - | - |
| AAVrh10 | 0.061 | 0.061 | 0.061 | 0.643 | 0.949 | 0.791 | 0.949 | 0.551 | - |
| AAVrh74 | 0.061 | 0.061 | 0.061 | 0.643 | 0.643 | 1.000 | 0.791 | 0.198 | 1.000 |

**Bladder**

| | AAV3b | AAV4 | AAV5 | AAV6 | AAV7 | AAV8 | AAV9 | AAVrh8 | AAVrh10 |
|---|---|---|---|---|---|---|---|---|---|
| AAV4 | 0.086 | - | - | - | - | - | - | - | - |
| AAV5 | 0.086 | 0.906 | - | - | - | - | - | - | - |
| AAV6 | 0.063 | 0.765 | 1.000 | - | - | - | - | - | - |
| AAV7 | 0.063 | 0.063 | 0.063 | 0.063 | - | - | - | - | - |
| AAV8 | 0.063 | 0.063 | 0.063 | 0.063 | 0.161 | - | - | - | - |
| AAV9 | 0.086 | 0.086 | 0.086 | 0.086 | 0.906 | 0.086 | - | - | - |
| AAVrh8 | 0.063 | 0.063 | 0.063 | 0.063 | 0.250 | 0.906 | 0.086 | - | - |
| AAVrh10 | 0.063 | 0.063 | 0.063 | 0.063 | 0.161 | 0.906 | 0.086 | 0.906 | - |
| AAVrh74 | 0.063 | 0.063 | 0.063 | 0.063 | 0.575 | 0.250 | 0.278 | 0.250 | 0.250 |

**Brain**

| | AAV3b | AAV4 | AAV5 | AAV6 | AAV7 | AAV8 | AAV9 | AAVrh8 | AAVrh10 |
|---|---|---|---|---|---|---|---|---|---|
| AAV4 | 0.068 | - | - | - | - | - | - | - | - |
| AAV5 | 0.068 | 0.454 | - | - | - | - | - | - | - |
| AAV6 | 0.068 | 0.092 | 0.068 | - | - | - | - | - | - |
| AAV7 | 0.068 | 0.092 | 0.092 | 0.591 | - | - | - | - | - |
| AAV8 | 0.068 | 0.171 | 0.092 | 0.718 | 0.591 | - | - | - | - |
| AAV9 | 0.068 | 0.281 | 0.092 | 0.718 | 0.591 | 0.906 | - | - | - |
| AAVrh8 | 0.068 | 0.092 | 0.068 | 0.718 | 0.718 | 0.718 | 0.718 | - | - |
| AAVrh10 | 0.068 | 0.068 | 0.068 | 0.068 | 0.454 | 0.092 | 0.171 | 0.068 | - |
| AAVrh74 | 0.068 | 0.068 | 0.068 | 0.068 | 0.281 | 0.092 | 0.092 | 0.068 | 1.000 |

**Diaphragm**

| | AAV3b | AAV4 | AAV5 | AAV6 | AAV7 | AAV8 | AAV9 | AAVrh8 | AAVrh10 |
|---|---|---|---|---|---|---|---|---|---|
| AAV4 | 0.391 | - | - | - | - | - | - | - | - |
| AAV5 | 0.791 | 0.064 | - | - | - | - | - | - | - |
| AAV6 | 0.064 | 0.064 | 0.064 | - | - | - | - | - | - |
| AAV7 | 0.064 | 0.064 | 0.064 | 1.000 | - | - | - | - | - |
| AAV8 | 0.122 | 0.391 | 0.064 | 0.514 | 0.791 | - | - | - | - |
| AAV9 | 0.064 | 0.064 | 0.064 | 0.972 | 1.000 | 0.791 | - | - | - |
| AAVrh8 | 0.064 | 0.064 | 0.064 | 0.514 | 0.624 | 0.514 | 0.624 | - | - |
| AAVrh10 | 0.064 | 0.064 | 0.064 | 0.624 | 0.972 | 0.791 | 1.000 | 0.624 | - |
| AAVrh74 | 0.064 | 0.064 | 0.064 | 0.514 | 0.624 | 0.514 | 0.514 | 1.000 | 0.514 |

Supplemental Table S7 - Wilcoxon Rank Tests - Functional Transduction Female

**Eye**

| | AAV3b | AAV4 | AAV5 | AAV6 | AAV7 | AAV8 | AAV9 | AAVrh8 | AAVrh10 |
|---|---|---|---|---|---|---|---|---|---|
| AAV4 | 0.051 | - | - | - | - | - | - | - | - |
| AAV5 | 0.051 | 0.051 | - | - | - | - | - | - | - |
| AAV6 | 0.051 | 0.432 | 0.051 | - | - | - | - | - | - |
| AAV7 | 0.051 | 0.051 | 0.051 | 0.092 | - | - | - | - | - |
| AAV8 | 0.051 | 0.051 | 0.051 | 0.156 | 0.886 | - | - | - | - |
| AAV9 | 0.051 | 0.396 | 0.156 | 0.886 | 0.092 | 0.092 | - | - | - |
| AAVrh8 | 0.051 | 0.051 | 0.051 | 0.051 | 0.520 | 0.396 | 0.051 | - | - |
| AAVrh10 | 0.051 | 0.051 | 0.051 | 0.156 | 0.396 | 0.396 | 0.156 | 0.396 | - |
| AAVrh74 | 0.051 | 0.051 | 0.051 | 0.051 | 0.520 | 0.156 | 0.051 | 0.718 | 0.396 |

**Gastrocnemius muscle**

| | AAV3b | AAV4 | AAV5 | AAV6 | AAV7 | AAV8 | AAV9 | AAVrh8 | AAVrh10 |
|---|---|---|---|---|---|---|---|---|---|
| AAV4 | 0.230 | - | - | - | - | - | - | - | - |
| AAV5 | 0.550 | 0.230 | - | - | - | - | - | - | - |
| AAV6 | 0.110 | 0.110 | 0.140 | - | - | - | - | - | - |
| AAV7 | 0.230 | 0.140 | 0.710 | 0.860 | - | - | - | - | - |
| AAV8 | 0.110 | 0.110 | 0.140 | 0.710 | 0.330 | - | - | - | - |
| AAV9 | 0.110 | 0.110 | 0.140 | 0.860 | 0.330 | 0.970 | - | - | - |
| AAVrh8 | 0.110 | 0.110 | 0.140 | 0.860 | 0.330 | 0.970 | 0.970 | - | - |
| AAVrh10 | 0.110 | 0.110 | 0.140 | 0.710 | 0.330 | 0.970 | 0.860 | 1.000 | - |
| AAVrh74 | 0.110 | 0.110 | 0.140 | 0.860 | 0.330 | 1.000 | 0.970 | 1.000 | 1.000 |

**Heart**

| | AAV3b | AAV4 | AAV5 | AAV6 | AAV7 | AAV8 | AAV9 | AAVrh8 | AAVrh10 |
|---|---|---|---|---|---|---|---|---|---|
| AAV4 | 0.056 | - | - | - | - | - | - | - | - |
| AAV5 | 0.056 | 0.498 | - | - | - | - | - | - | - |
| AAV6 | 0.056 | 0.056 | 0.056 | - | - | - | - | - | - |
| AAV7 | 0.056 | 0.056 | 0.056 | 0.498 | - | - | - | - | - |
| AAV8 | 0.056 | 0.056 | 0.056 | 0.206 | 0.886 | - | - | - | - |
| AAV9 | 0.056 | 0.056 | 0.056 | 0.886 | 0.498 | 0.333 | - | - | - |
| AAVrh8 | 0.056 | 0.056 | 0.056 | 0.662 | 0.771 | 0.662 | 0.771 | - | - |
| AAVrh10 | 0.056 | 0.056 | 0.056 | 0.333 | 0.886 | 0.886 | 0.771 | 0.886 | - |
| AAVrh74 | 0.056 | 0.056 | 0.056 | 0.206 | 0.771 | 0.771 | 0.498 | 0.771 | 0.771 |

**Kidney**

| | AAV3b | AAV4 | AAV5 | AAV6 | AAV7 | AAV8 | AAV9 | AAVrh8 | AAVrh10 |
|---|---|---|---|---|---|---|---|---|---|
| AAV4 | 0.058 | - | - | - | - | - | - | - | - |
| AAV5 | 0.058 | 0.058 | - | - | - | - | - | - | - |
| AAV6 | 0.058 | 0.058 | 0.906 | - | - | - | - | - | - |
| AAV7 | 0.058 | 0.058 | 0.624 | 1.000 | - | - | - | - | - |
| AAV8 | 0.058 | 0.058 | 0.906 | 0.906 | 0.624 | - | - | - | - |
| AAV9 | 0.058 | 0.514 | 0.112 | 0.058 | 0.058 | 0.058 | - | - | - |
| AAVrh8 | 0.058 | 0.058 | 0.514 | 0.514 | 0.514 | 0.514 | 0.058 | - | - |
| AAVrh10 | 0.058 | 0.058 | 0.624 | 0.624 | 0.514 | 0.857 | 0.058 | 0.514 | - |
| AAVrh74 | 0.058 | 0.058 | 0.906 | 0.906 | 0.906 | 0.906 | 0.058 | 0.624 | 0.906 |

Supplemental Table S7 - Wilcoxon Rank Tests - Functional Transduction Female

**Large Intestine**

| | AAV3b | AAV4 | AAV5 | AAV6 | AAV7 | AAV8 | AAV9 | AAVrh8 | AAVrh10 |
|---|---|---|---|---|---|---|---|---|---|
| AAV4 | 0.061 | - | - | - | - | - | - | - | - |
| AAV5 | 0.086 | 0.725 | - | - | - | - | - | - | - |
| AAV6 | 0.061 | 0.265 | 0.486 | - | - | - | - | - | - |
| AAV7 | 0.061 | 0.061 | 0.086 | 0.061 | - | - | - | - | - |
| AAV8 | 0.061 | 0.061 | 0.086 | 0.061 | 0.265 | - | - | - | - |
| AAV9 | 0.061 | 0.061 | 0.086 | 0.061 | 0.429 | 0.166 | - | - | - |
| AAVrh8 | 0.061 | 0.061 | 0.086 | 0.061 | 0.575 | 0.753 | 0.061 | - | - |
| AAVrh10 | 0.061 | 0.061 | 0.086 | 0.061 | 0.265 | 1.000 | 0.086 | 0.927 | - |
| AAVrh74 | 0.061 | 0.061 | 0.086 | 0.061 | 0.429 | 1.000 | 0.086 | 0.927 | 0.753 |

**Liver**

| | AAV3b | AAV4 | AAV5 | AAV6 | AAV7 | AAV8 | AAV9 | AAVrh8 | AAVrh10 |
|---|---|---|---|---|---|---|---|---|---|
| AAV4 | 0.076 | - | - | - | - | - | - | - | - |
| AAV5 | 0.076 | 0.076 | - | - | - | - | - | - | - |
| AAV6 | 0.076 | 0.076 | 0.729 | - | - | - | - | - | - |
| AAV7 | 0.076 | 0.076 | 0.257 | 0.834 | - | - | - | - | - |
| AAV8 | 0.076 | 0.076 | 0.409 | 0.834 | 0.834 | - | - | - | - |
| AAV9 | 0.076 | 0.076 | 0.729 | 0.949 | 0.643 | 0.949 | - | - | - |
| AAVrh8 | 0.076 | 0.076 | 0.729 | 1.000 | 0.729 | 0.890 | 1.000 | - | - |
| AAVrh10 | 0.076 | 0.076 | 0.409 | 1.000 | 0.890 | 0.949 | 0.789 | 0.729 | - |
| AAVrh74 | 0.076 | 0.076 | 0.643 | 0.729 | 0.257 | 0.834 | 0.834 | 0.789 | 0.257 |

**Lung**

| | AAV3b | AAV4 | AAV5 | AAV6 | AAV7 | AAV8 | AAV9 | AAVrh8 | AAVrh10 |
|---|---|---|---|---|---|---|---|---|---|
| AAV4 | 0.110 | - | - | - | - | - | - | - | - |
| AAV5 | 0.110 | 0.970 | - | - | - | - | - | - | - |
| AAV6 | 0.110 | 0.710 | 0.970 | - | - | - | - | - | - |
| AAV7 | 0.110 | 0.710 | 1.000 | 0.970 | - | - | - | - | - |
| AAV8 | 0.110 | 1.000 | 0.970 | 0.550 | 0.910 | - | - | - | - |
| AAV9 | 0.110 | 0.430 | 0.430 | 0.430 | 0.550 | 0.320 | - | - | - |
| AAVrh8 | 0.110 | 0.970 | 0.910 | 0.970 | 0.970 | 0.910 | 0.550 | - | - |
| AAVrh10 | 0.110 | 0.550 | 0.430 | 0.110 | 0.110 | 0.320 | 0.110 | 0.320 | - |
| AAVrh74 | 0.110 | 1.000 | 0.710 | 0.430 | 0.550 | 1.000 | 0.200 | 0.550 | 0.550 |

**Lymph nodes**

| | AAV3b | AAV4 | AAV5 | AAV6 | AAV7 | AAV8 | AAV9 | AAVrh8 | AAVrh10 |
|---|---|---|---|---|---|---|---|---|---|
| AAV4 | 0.120 | - | - | - | - | - | - | - | - |
| AAV5 | 0.120 | 0.120 | - | - | - | - | - | - | - |
| AAV6 | 0.120 | 0.120 | 0.610 | - | - | - | - | - | - |
| AAV7 | 0.120 | 0.120 | 0.610 | 0.950 | - | - | - | - | - |
| AAV8 | 0.120 | 0.120 | 1.000 | 0.530 | 0.730 | - | - | - | - |
| AAV9 | 0.120 | 0.120 | 1.000 | 1.000 | 0.120 | 0.790 | - | - | - |
| AAVrh8 | 0.120 | 0.120 | 0.210 | 0.320 | 0.320 | 0.210 | 0.120 | - | - |
| AAVrh10 | 0.120 | 0.120 | 0.120 | 0.320 | 0.210 | 0.210 | 0.120 | 0.950 | - |
| AAVrh74 | 0.120 | 0.120 | 0.330 | 0.730 | 0.730 | 0.530 | 0.530 | 0.330 | 0.330 |

Supplemental Table S7 - Wilcoxon Rank Tests - Functional Transduction Female

**Pancreas**

| | AAV3b | AAV4 | AAV5 | AAV6 | AAV7 | AAV8 | AAV9 | AAVrh8 | AAVrh10 |
|---|---|---|---|---|---|---|---|---|---|
| AAV4 | 0.041 | - | - | - | - | - | - | - | - |
| AAV5 | 0.041 | 0.243 | - | - | - | - | - | - | - |
| AAV6 | 0.041 | 0.520 | 0.376 | - | - | - | - | - | - |
| AAV7 | 0.041 | 0.041 | 0.078 | 0.041 | - | - | - | - | - |
| AAV8 | 0.041 | 0.041 | 0.041 | 0.041 | 0.906 | - | - | - | - |
| AAV9 | 0.041 | 0.041 | 0.041 | 0.041 | 0.718 | 0.243 | - | - | - |
| AAVrh8 | 0.041 | 0.041 | 0.041 | 0.041 | 0.376 | 0.243 | 0.376 | - | - |
| AAVrh10 | 0.041 | 0.041 | 0.041 | 0.041 | 0.243 | 0.041 | 0.041 | 0.041 | - |
| AAVrh74 | 0.041 | 0.041 | 0.041 | 0.041 | 0.376 | 0.041 | 0.041 | 0.041 | 1.000 |

**Skin**

| | AAV3b | AAV4 | AAV5 | AAV6 | AAV7 | AAV8 | AAV9 | AAVrh8 | AAVrh10 |
|---|---|---|---|---|---|---|---|---|---|
| AAV4 | 0.046 | - | - | - | - | - | - | - | - |
| AAV5 | 0.046 | 0.546 | - | - | - | - | - | - | - |
| AAV6 | 0.046 | 0.147 | 0.718 | - | - | - | - | - | - |
| AAV7 | 0.046 | 0.046 | 0.046 | 0.046 | - | - | - | - | - |
| AAV8 | 0.046 | 0.046 | 0.046 | 0.086 | 0.086 | - | - | - | - |
| AAV9 | 0.046 | 0.046 | 0.046 | 0.046 | 0.147 | 0.718 | - | - | - |
| AAVrh8 | 0.046 | 0.046 | 0.046 | 0.046 | 0.546 | 0.147 | 0.147 | - | - |
| AAVrh10 | 0.046 | 0.046 | 0.046 | 0.046 | 0.886 | 0.406 | 0.406 | 0.718 | - |
| AAVrh74 | 0.046 | 0.046 | 0.046 | 0.046 | 0.250 | 0.046 | 0.046 | 0.147 | 0.886 |

**Small Intestine**

| | AAV3b | AAV4 | AAV5 | AAV6 | AAV7 | AAV8 | AAV9 | AAVrh8 | AAVrh10 |
|---|---|---|---|---|---|---|---|---|---|
| AAV4 | 0.056 | - | - | - | - | - | - | - | - |
| AAV5 | 0.056 | 0.417 | - | - | - | - | - | - | - |
| AAV6 | 0.056 | 0.177 | 0.753 | - | - | - | - | - | - |
| AAV7 | 0.056 | 0.056 | 0.056 | 0.056 | - | - | - | - | - |
| AAV8 | 0.056 | 0.056 | 0.056 | 0.056 | 0.753 | - | - | - | - |
| AAV9 | 0.056 | 0.103 | 0.056 | 0.056 | 0.906 | 0.906 | - | - | - |
| AAVrh8 | 0.056 | 0.753 | 0.273 | 0.177 | 0.273 | 0.103 | 0.177 | - | - |
| AAVrh10 | 0.056 | 0.056 | 0.056 | 0.056 | 0.417 | 0.753 | 0.273 | 0.177 | - |
| AAVrh74 | 0.056 | 0.056 | 0.056 | 0.056 | 0.906 | 0.417 | 1.000 | 0.273 | 0.417 |

**Spleen**

| | AAV3b | AAV4 | AAV5 | AAV6 | AAV7 | AAV8 | AAV9 | AAVrh8 | AAVrh10 |
|---|---|---|---|---|---|---|---|---|---|
| AAV4 | 0.048 | - | - | - | - | - | - | - | - |
| AAV5 | 0.048 | 0.048 | - | - | - | - | - | - | - |
| AAV6 | 0.048 | 0.048 | 0.048 | - | - | - | - | - | - |
| AAV7 | 0.048 | 0.048 | 0.048 | 0.048 | - | - | - | - | - |
| AAV8 | 0.048 | 0.048 | 0.701 | 0.406 | 0.086 | - | - | - | - |
| AAV9 | 0.048 | 0.048 | 0.250 | 0.048 | 0.048 | 0.406 | - | - | - |
| AAVrh8 | 0.048 | 0.048 | 0.250 | 0.533 | 0.086 | 0.533 | 0.086 | - | - |
| AAVrh10 | 0.048 | 0.048 | 0.048 | 0.048 | 0.886 | 0.156 | 0.048 | 0.156 | - |
| AAVrh74 | 0.048 | 0.048 | 0.048 | 0.156 | 0.533 | 0.250 | 0.048 | 0.601 | 0.701 |

Supplemental Table S7 - Wilcoxon Rank Tests - Functional Transduction Female

**Stomach**

| | AAV3b | AAV4 | AAV5 | AAV6 | AAV7 | AAV8 | AAV9 | AAVrh8 | AAVrh10 |
|---|---|---|---|---|---|---|---|---|---|
| AAV4 | 0.064 | - | - | - | - | - | - | - | - |
| AAV5 | 0.064 | 0.735 | - | - | - | - | - | - | - |
| AAV6 | 0.064 | 0.441 | 0.886 | - | - | - | - | - | - |
| AAV7 | 0.064 | 0.300 | 0.190 | 0.300 | - | - | - | - | - |
| AAV8 | 0.064 | 0.064 | 0.190 | 0.064 | 0.735 | - | - | - | - |
| AAV9 | 0.064 | 0.190 | 0.107 | 0.107 | 0.575 | 0.886 | - | - | - |
| AAVrh8 | 0.064 | 0.064 | 0.064 | 0.064 | 0.064 | 0.441 | 0.441 | - | - |
| AAVrh10 | 0.064 | 0.064 | 0.064 | 0.064 | 0.064 | 0.441 | 0.441 | 0.575 | - |
| AAVrh74 | 0.064 | 0.107 | 0.107 | 0.064 | 0.300 | 0.735 | 0.575 | 0.886 | 0.735 |

**Thymus**

| | AAV3b | AAV4 | AAV5 | AAV6 | AAV7 | AAV8 | AAV9 | AAVrh8 | AAVrh10 |
|---|---|---|---|---|---|---|---|---|---|
| AAV4 | 0.055 | - | - | - | - | - | - | - | - |
| AAV5 | 0.055 | 0.273 | - | - | - | - | - | - | - |
| AAV6 | 0.055 | 1.000 | 0.417 | - | - | - | - | - | - |
| AAV7 | 0.055 | 0.055 | 0.055 | 0.055 | - | - | - | - | - |
| AAV8 | 0.055 | 0.055 | 0.086 | 0.055 | 0.771 | - | - | - | - |
| AAV9 | 0.055 | 0.055 | 0.055 | 0.055 | 1.000 | 0.417 | - | - | - |
| AAVrh8 | 0.086 | 0.086 | 0.086 | 0.086 | 0.303 | 0.474 | 0.086 | - | - |
| AAVrh10 | 0.055 | 0.055 | 0.055 | 0.055 | 0.166 | 0.771 | 0.055 | 0.941 | - |
| AAVrh74 | 0.055 | 0.055 | 0.055 | 0.055 | 0.273 | 0.417 | 0.055 | 1.000 | 0.949 |

**Ovary**

| | AAV3b | AAV4 | AAV5 | AAV6 | AAV7 | AAV8 | AAV9 | AAVrh8 | AAVrh10 |
|---|---|---|---|---|---|---|---|---|---|
| AAV4 | 0.112 | - | - | - | - | - | - | - | - |
| AAV5 | 0.112 | 0.112 | - | - | - | - | - | - | - |
| AAV6 | 0.086 | 0.112 | 0.332 | - | - | - | - | - | - |
| AAV7 | 0.112 | 0.112 | 0.500 | 0.190 | - | - | - | - | - |
| AAV8 | 0.086 | 0.086 | 0.332 | 0.086 | 0.897 | - | - | - | - |
| AAV9 | 0.086 | 0.086 | 0.690 | 0.454 | 0.500 | 0.086 | - | - | - |
| AAVrh8 | 0.086 | 0.086 | 0.897 | 0.112 | 0.690 | 0.454 | 0.190 | - | - |
| AAVrh10 | 0.086 | 0.086 | 0.690 | 0.112 | 0.690 | 0.591 | 0.190 | 1.000 | - |
| AAVrh74 | 0.086 | 0.086 | 0.332 | 0.086 | 1.000 | 0.454 | 0.086 | 0.190 | 0.321 |

**Uterus**

| | AAV3b | AAV4 | AAV5 | AAV6 | AAV7 | AAV8 | AAV9 | AAVrh8 | AAVrh10 |
|---|---|---|---|---|---|---|---|---|---|
| AAV4 | 0.044 | - | - | - | - | - | - | - | - |
| AAV5 | 0.044 | 0.735 | - | - | - | - | - | - | - |
| AAV6 | 0.044 | 0.735 | 0.906 | - | - | - | - | - | - |
| AAV7 | 0.044 | 0.044 | 0.044 | 0.080 | - | - | - | - | - |
| AAV8 | 0.044 | 0.044 | 0.080 | 0.156 | 0.560 | - | - | - | - |
| AAV9 | 0.044 | 0.265 | 0.417 | 0.906 | 0.560 | 0.417 | - | - | - |
| AAVrh8 | 0.044 | 0.044 | 0.044 | 0.044 | 0.044 | 0.080 | 0.044 | - | - |
| AAVrh10 | 0.044 | 0.044 | 0.044 | 0.044 | 0.044 | 0.044 | 0.044 | 0.735 | - |
| AAVrh74 | 0.044 | 0.044 | 0.044 | 0.044 | 0.044 | 0.044 | 0.044 | 1.000 | 0.417 |

Supplemental Table S8 - Wilcoxon Rank Tests - Flow Cytometry Male

**CLP**

| | AAV3b | AAV4 | AAV5 | AAV6 | AAV7 | AAV8 | AAV9 | AAVrh8 | AAVrh10 |
|---|---|---|---|---|---|---|---|---|---|
| AAV4 | 0.007 | - | - | - | - | - | - | - | - |
| AAV5 | 0.010 | 0.018 | - | - | - | - | - | - | - |
| AAV6 | 0.007 | 0.135 | 0.007 | - | - | - | - | - | - |
| AAV7 | 0.007 | 0.018 | 0.007 | 0.108 | - | - | - | - | - |
| AAV8 | 0.007 | 0.135 | 0.007 | 0.749 | 0.180 | - | - | - | - |
| AAV9 | 0.010 | 0.010 | 0.010 | 0.132 | 0.931 | 0.177 | - | - | - |
| AAVrh8 | 0.007 | 0.007 | 0.007 | 0.108 | 0.837 | 0.081 | 0.619 | - | - |
| AAVrh10 | 0.007 | 0.010 | 0.007 | 0.108 | 0.574 | 0.749 | 0.317 | 0.135 | - |
| AAVrh74 | 0.007 | 0.010 | 0.007 | 0.108 | 0.574 | 0.317 | 0.829 | 0.662 | 0.492 |

**CMP**

| | AAV3b | AAV4 | AAV5 | AAV6 | AAV7 | AAV8 | AAV9 | AAVrh8 | AAVrh10 |
|---|---|---|---|---|---|---|---|---|---|
| AAV4 | 0.004 | - | - | - | - | - | - | - | - |
| AAV5 | 0.034 | 0.110 | - | - | - | - | - | - | - |
| AAV6 | 0.004 | 0.034 | 0.007 | - | - | - | - | - | - |
| AAV7 | 0.004 | 0.004 | 0.004 | 0.004 | - | - | - | - | - |
| AAV8 | 0.004 | 0.004 | 0.004 | 0.053 | 0.013 | - | - | - | - |
| AAV9 | 0.007 | 0.007 | 0.007 | 0.063 | 0.063 | 0.829 | - | - | - |
| AAVrh8 | 0.004 | 0.004 | 0.004 | 0.004 | 0.837 | 0.007 | 0.013 | - | - |
| AAVrh10 | 0.004 | 0.004 | 0.004 | 0.004 | 0.646 | 0.007 | 0.024 | 0.937 | - |
| AAVrh74 | 0.004 | 0.004 | 0.004 | 0.004 | 0.270 | 0.004 | 0.007 | 0.270 | 0.749 |

**GMP**

| | AAV3b | AAV4 | AAV5 | AAV6 | AAV7 | AAV8 | AAV9 | AAVrh8 | AAVrh10 |
|---|---|---|---|---|---|---|---|---|---|
| AAV4 | 0.004 | - | - | - | - | - | - | - | - |
| AAV5 | 0.048 | 0.149 | - | - | - | - | - | - | - |
| AAV6 | 0.004 | 0.077 | 0.020 | - | - | - | - | - | - |
| AAV7 | 0.004 | 0.004 | 0.004 | 0.050 | - | - | - | - | - |
| AAV8 | 0.004 | 0.004 | 0.004 | 0.050 | 0.412 | - | - | - | - |
| AAV9 | 0.006 | 0.006 | 0.006 | 0.095 | 0.810 | 0.361 | - | - | - |
| AAVrh8 | 0.004 | 0.004 | 0.004 | 0.004 | 0.004 | 0.006 | 0.006 | - | - |
| AAVrh10 | 0.004 | 0.004 | 0.004 | 0.004 | 0.004 | 0.012 | 0.006 | 0.818 | - |
| AAVrh74 | 0.004 | 0.004 | 0.004 | 0.004 | 0.004 | 0.004 | 0.006 | 0.020 | 0.412 |

**MEP**

| | AAV3b | AAV4 | AAV5 | AAV6 | AAV7 | AAV8 | AAV9 | AAVrh8 | AAVrh10 |
|---|---|---|---|---|---|---|---|---|---|
| AAV4 | 0.012 | - | - | - | - | - | - | - | - |
| AAV5 | 0.022 | 1.000 | - | - | - | - | - | - | - |
| AAV6 | 0.008 | 0.051 | 0.022 | - | - | - | - | - | - |
| AAV7 | 0.008 | 0.006 | 0.006 | 1.000 | - | - | - | - | - |
| AAV8 | 0.008 | 0.113 | 0.051 | 0.340 | 0.008 | - | - | - | - |
| AAV9 | 0.012 | 0.024 | 0.008 | 0.895 | 0.149 | 0.202 | - | - | - |
| AAVrh8 | 0.008 | 0.006 | 0.006 | 0.008 | 0.006 | 0.006 | 0.008 | - | - |
| AAVrh10 | 0.008 | 0.006 | 0.006 | 0.006 | 0.006 | 0.006 | 0.008 | 0.034 | - |
| AAVrh74 | 0.008 | 0.006 | 0.006 | 0.006 | 0.006 | 0.006 | 0.008 | 0.422 | 0.202 |

Supplemental Table S8 - Wilcoxon Rank Tests - Flow Cytometry Male

**LT-HSC**

| | AAV3b | AAV4 | AAV5 | AAV6 | AAV7 | AAV8 | AAV9 | AAVrh8 | AAVrh10 |
|---|---|---|---|---|---|---|---|---|---|
| AAV4 | 0.008 | - | - | - | - | - | - | - | - |
| AAV5 | 0.043 | 0.091 | - | - | - | - | - | - | - |
| AAV6 | 0.008 | 0.837 | 0.042 | - | - | - | - | - | - |
| AAV7 | 0.008 | 0.008 | 0.008 | 0.008 | - | - | - | - | - |
| AAV8 | 0.008 | 0.091 | 0.008 | 0.165 | 0.443 | - | - | - | - |
| AAV9 | 0.012 | 1.000 | 0.046 | 0.710 | 0.008 | 0.112 | - | - | - |
| AAVrh8 | 0.008 | 0.008 | 0.008 | 0.008 | 0.219 | 0.120 | 0.008 | - | - |
| AAVrh10 | 0.008 | 0.008 | 0.008 | 0.008 | 0.041 | 0.015 | 0.008 | 0.646 | - |
| AAVrh74 | 0.008 | 0.008 | 0.008 | 0.008 | 0.357 | 0.120 | 0.008 | 0.732 | 0.357 |

**ST-HSC**

| | AAV3b | AAV4 | AAV5 | AAV6 | AAV7 | AAV8 | AAV9 | AAVrh8 | AAVrh10 |
|---|---|---|---|---|---|---|---|---|---|
| AAV4 | 0.009 | - | - | - | - | - | - | - | - |
| AAV5 | 0.043 | 0.277 | - | - | - | - | - | - | - |
| AAV6 | 0.009 | 0.009 | 0.009 | - | - | - | - | - | - |
| AAV7 | 0.009 | 0.009 | 0.009 | 0.009 | - | - | - | - | - |
| AAV8 | 0.009 | 0.009 | 0.009 | 0.040 | 0.025 | - | - | - | - |
| AAV9 | 0.009 | 0.009 | 0.014 | 0.693 | 0.009 | 0.044 | - | - | - |
| AAVrh8 | 0.009 | 0.009 | 0.009 | 0.058 | 0.277 | 0.348 | 0.071 | - | - |
| AAVrh10 | 0.009 | 0.009 | 0.009 | 0.009 | 0.277 | 0.170 | 0.009 | 0.837 | - |
| AAVrh74 | 0.009 | 0.009 | 0.009 | 0.040 | 0.225 | 0.520 | 0.109 | 1.000 | 0.520 |

**MPP2**

| | AAV3b | AAV4 | AAV5 | AAV6 | AAV7 | AAV8 | AAV9 | AAVrh8 | AAVrh10 |
|---|---|---|---|---|---|---|---|---|---|
| AAV4 | 0.007 | - | - | - | - | - | - | - | - |
| AAV5 | 0.016 | 0.024 | - | - | - | - | - | - | - |
| AAV6 | 0.007 | 0.697 | 0.038 | - | - | - | - | - | - |
| AAV7 | 0.007 | 0.007 | 0.007 | 0.007 | - | - | - | - | - |
| AAV8 | 0.007 | 0.008 | 0.007 | 0.008 | 0.697 | - | - | - | - |
| AAV9 | 0.008 | 0.837 | 0.043 | 0.567 | 0.008 | 0.026 | - | - | - |
| AAVrh8 | 0.007 | 0.007 | 0.007 | 0.008 | 0.623 | 0.767 | 0.026 | - | - |
| AAVrh10 | 0.007 | 0.007 | 0.007 | 0.007 | 0.422 | 0.837 | 0.008 | 0.837 | - |
| AAVrh74 | 0.007 | 0.007 | 0.007 | 0.007 | 0.937 | 0.767 | 0.008 | 0.651 | 0.767 |

**MPP3**

| | AAV3b | AAV4 | AAV5 | AAV6 | AAV7 | AAV8 | AAV9 | AAVrh8 | AAVrh10 |
|---|---|---|---|---|---|---|---|---|---|
| AAV4 | 0.008 | - | - | - | - | - | - | - | - |
| AAV5 | 0.024 | 0.116 | - | - | - | - | - | - | - |
| AAV6 | 0.008 | 0.086 | 0.013 | - | - | - | - | - | - |
| AAV7 | 0.008 | 0.008 | 0.008 | 0.008 | - | - | - | - | - |
| AAV8 | 0.008 | 0.008 | 0.008 | 0.019 | 0.205 | - | - | - | - |
| AAV9 | 0.009 | 0.008 | 0.008 | 0.361 | 0.008 | 0.075 | - | - | - |
| AAVrh8 | 0.008 | 0.008 | 0.008 | 0.008 | 0.856 | 0.043 | 0.008 | - | - |
| AAVrh10 | 0.008 | 0.008 | 0.008 | 0.008 | 0.348 | 1.000 | 0.008 | 0.207 | - |
| AAVrh74 | 0.008 | 0.008 | 0.008 | 0.008 | 0.161 | 0.957 | 0.008 | 0.100 | 0.856 |

Supplemental Table S8 - Wilcoxon Rank Tests - Flow Cytometry Male

**MPP4**

| | AAV3b | AAV4 | AAV5 | AAV6 | AAV7 | AAV8 | AAV9 | AAVrh8 | AAVrh10 |
|---|---|---|---|---|---|---|---|---|---|
| AAV4 | 0.005 | - | - | - | - | - | - | - | - |
| AAV5 | 0.072 | 0.013 | - | - | - | - | - | - | - |
| AAV6 | 0.005 | 0.056 | 0.007 | - | - | - | - | - | - |
| AAV7 | 0.005 | 0.005 | 0.005 | 0.005 | - | - | - | - | - |
| AAV8 | 0.005 | 0.007 | 0.005 | 0.056 | 0.056 | - | - | - | - |
| AAV9 | 0.007 | 0.013 | 0.007 | 0.278 | 0.007 | 0.103 | - | - | - |
| AAVrh8 | 0.005 | 0.005 | 0.005 | 0.005 | 0.631 | 0.084 | 0.007 | - | - |
| AAVrh10 | 0.005 | 0.005 | 0.005 | 0.005 | 0.715 | 0.277 | 0.007 | 0.937 | - |
| AAVrh74 | 0.005 | 0.005 | 0.005 | 0.005 | 0.213 | 0.572 | 0.007 | 0.213 | 0.660 |

**CD4+ T cells**

| | AAV3b | AAV4 | AAV5 | AAV6 | AAV7 | AAV8 | AAV9 | AAVrh8 | AAVrh10 |
|---|---|---|---|---|---|---|---|---|---|
| AAV4 | 0.059 | - | - | - | - | - | - | - | - |
| AAV5 | 0.017 | 0.519 | - | - | - | - | - | - | - |
| AAV6 | 0.017 | 0.245 | 0.892 | - | - | - | - | - | - |
| AAV7 | 0.017 | 0.017 | 0.017 | 0.140 | - | - | - | - | - |
| AAV8 | 0.017 | 0.022 | 0.017 | 0.376 | 0.031 | - | - | - | - |
| AAV9 | 0.022 | 0.022 | 0.022 | 0.245 | 0.182 | 0.470 | - | - | - |
| AAVrh8 | 0.017 | 0.017 | 0.017 | 0.376 | 0.031 | 0.856 | 0.390 | - | - |
| AAVrh10 | 0.017 | 0.071 | 0.071 | 0.443 | 0.017 | 0.104 | 0.087 | 0.031 | - |
| AAVrh74 | 0.017 | 0.071 | 0.031 | 0.443 | 0.022 | 0.309 | 0.128 | 0.309 | 1.000 |

**CD8+ T cells**

| | AAV3b | AAV4 | AAV5 | AAV6 | AAV7 | AAV8 | AAV9 | AAVrh8 | AAVrh10 |
|---|---|---|---|---|---|---|---|---|---|
| AAV4 | 0.270 | - | - | - | - | - | - | - | - |
| AAV5 | 0.337 | 1.000 | - | - | - | - | - | - | - |
| AAV6 | 0.076 | 0.154 | 0.516 | - | - | - | - | - | - |
| AAV7 | 0.024 | 0.024 | 0.152 | 0.416 | - | - | - | - | - |
| AAV8 | 0.154 | 0.807 | 0.716 | 0.554 | 0.154 | - | - | - | - |
| AAV9 | 0.154 | 0.567 | 0.981 | 0.671 | 0.056 | 0.981 | - | - | - |
| AAVrh8 | 0.117 | 0.270 | 0.416 | 1.000 | 0.807 | 0.554 | 0.529 | - | - |
| AAVrh10 | 0.024 | 0.024 | 0.056 | 0.209 | 0.154 | 0.154 | 0.056 | 0.337 | - |
| AAVrh74 | 0.076 | 0.154 | 0.554 | 0.981 | 0.554 | 0.623 | 0.567 | 0.981 | 0.154 |

**B cells**

| | AAV3b | AAV4 | AAV5 | AAV6 | AAV7 | AAV8 | AAV9 | AAVrh8 | AAVrh10 |
|---|---|---|---|---|---|---|---|---|---|
| AAV4 | 0.606 | - | - | - | - | - | - | - | - |
| AAV5 | 0.487 | 0.606 | - | - | - | - | - | - | - |
| AAV6 | 0.097 | 0.062 | 0.716 | - | - | - | - | - | - |
| AAV7 | 0.019 | 0.019 | 0.062 | 0.268 | - | - | - | - | - |
| AAV8 | 0.248 | 0.199 | 0.606 | 0.959 | 0.019 | - | - | - | - |
| AAV9 | 0.130 | 0.056 | 0.487 | 0.767 | 0.130 | 0.487 | - | - | - |
| AAVrh8 | 0.116 | 0.032 | 0.487 | 0.767 | 0.058 | 0.606 | 1.000 | - | - |
| AAVrh10 | 0.062 | 0.019 | 0.199 | 0.487 | 0.153 | 0.116 | 0.829 | 0.487 | - |
| AAVrh74 | 0.116 | 0.019 | 0.248 | 0.606 | 0.116 | 0.248 | 0.829 | 0.767 | 0.767 |

Supplemental Table S8 - Wilcoxon Rank Tests - Flow Cytometry Male

**Eosinophils**

| | AAV3b | AAV4 | AAV5 | AAV6 | AAV7 | AAV8 | AAV9 | AAVrh8 | AAVrh10 |
|---|---|---|---|---|---|---|---|---|---|
| AAV4 | 0.563 | - | - | - | - | - | - | - | - |
| AAV5 | 0.233 | 0.270 | - | - | - | - | - | - | - |
| AAV6 | 0.408 | 0.279 | 0.779 | - | - | - | - | - | - |
| AAV7 | 0.209 | 0.209 | 0.613 | 0.497 | - | - | - | - | - |
| AAV8 | 0.491 | 0.842 | 0.209 | 0.209 | 0.167 | - | - | - | - |
| AAV9 | 0.233 | 0.209 | 0.937 | 0.828 | 0.755 | 0.209 | - | - | - |
| AAVrh8 | 0.233 | 0.416 | 0.828 | 0.779 | 0.842 | 0.233 | 0.869 | - | - |
| AAVrh10 | 0.209 | 0.049 | 0.270 | 0.270 | 0.049 | 0.049 | 0.130 | 0.049 | - |
| AAVrh74 | 0.270 | 0.521 | 0.937 | 0.828 | 0.828 | 0.279 | 0.895 | 0.877 | 0.130 |

**CD11c+ monocytes**

| | AAV3b | AAV4 | AAV5 | AAV6 | AAV7 | AAV8 | AAV9 | AAVrh8 | AAVrh10 |
|---|---|---|---|---|---|---|---|---|---|
| AAV4 | 0.139 | - | - | - | - | - | - | - | - |
| AAV5 | 0.877 | 0.373 | - | - | - | - | - | - | - |
| AAV6 | 0.139 | 0.716 | 0.311 | - | - | - | - | - | - |
| AAV7 | 0.373 | 0.139 | 0.521 | 0.521 | - | - | - | - | - |
| AAV8 | 1.000 | 0.024 | 0.877 | 0.106 | 0.139 | - | - | - | - |
| AAV9 | 0.877 | 0.139 | 0.877 | 0.139 | 0.168 | 0.895 | - | - | - |
| AAVrh8 | 0.139 | 1.000 | 0.435 | 0.716 | 0.139 | 0.024 | 0.139 | - | - |
| AAVrh10 | 0.085 | 0.238 | 0.182 | 0.373 | 0.039 | 0.024 | 0.065 | 0.139 | - |
| AAVrh74 | 0.106 | 0.623 | 0.238 | 0.435 | 0.085 | 0.024 | 0.087 | 0.435 | 0.877 |

**Ly6C+ monocytes**

| | AAV3b | AAV4 | AAV5 | AAV6 | AAV7 | AAV8 | AAV9 | AAVrh8 | AAVrh10 |
|---|---|---|---|---|---|---|---|---|---|
| AAV4 | 0.036 | - | - | - | - | - | - | - | - |
| AAV5 | 0.188 | 0.049 | - | - | - | - | - | - | - |
| AAV6 | 0.027 | 0.192 | 0.028 | - | - | - | - | - | - |
| AAV7 | 0.02 | 0.049 | 0.016 | 0.492 | - | - | - | - | - |
| AAV8 | 0.044 | 0.532 | 0.067 | 0.069 | 0.016 | - | - | - | - |
| AAV9 | 0.036 | 0.575 | 0.041 | 0.508 | 0.02 | 0.128 | - | - | - |
| AAVrh8 | 0.02 | 0.41 | 0.016 | 0.492 | 0.016 | 0.049 | 1 | - | - |
| AAVrh10 | 0.02 | 0.049 | 0.016 | 0.532 | 0.837 | 0.016 | 0.041 | 0.04 | - |
| AAVrh74 | 0.027 | 0.532 | 0.02 | 0.41 | 0.104 | 0.41 | 0.693 | 0.508 | 0.069 |

**Neutrophils**

| | AAV3b | AAV4 | AAV5 | AAV6 | AAV7 | AAV8 | AAV9 | AAVrh8 | AAVrh10 |
|---|---|---|---|---|---|---|---|---|---|
| AAV4 | 0.708 | - | - | - | - | - | - | - | - |
| AAV5 | 0.442 | 0.937 | - | - | - | - | - | - | - |
| AAV6 | 0.037 | 0.103 | 0.32 | - | - | - | - | - | - |
| AAV7 | 0.037 | 0.062 | 0.073 | 0.406 | - | - | - | - | - |
| AAV8 | 0.073 | 0.299 | 0.937 | 0.238 | 0.037 | - | - | - | - |
| AAV9 | 0.073 | 0.154 | 0.636 | 0.472 | 0.117 | 0.727 | - | - | - |
| AAVrh8 | 0.037 | 0.062 | 0.073 | 0.59 | 0.559 | 0.056 | 0.117 | - | - |
| AAVrh10 | 0.037 | 0.073 | 0.133 | 0.679 | 0.537 | 0.062 | 0.154 | 0.59 | - |
| AAVrh74 | 0.037 | 0.103 | 0.133 | 0.802 | 0.472 | 0.062 | 0.299 | 0.59 | 0.937 |

Supplemental Table S8 - Wilcoxon Rank Tests - Flow Cytometry Male

**NK cells**

| | AAV3b | AAV4 | AAV5 | AAV6 | AAV7 | AAV8 | AAV9 | AAVrh8 | AAVrh10 |
|---|---|---|---|---|---|---|---|---|---|
| AAV4 | 0.010 | - | - | - | - | - | - | - | - |
| AAV5 | 0.010 | 0.410 | - | - | - | - | - | - | - |
| AAV6 | 0.010 | 0.311 | 0.338 | - | - | - | - | - | - |
| AAV7 | 0.017 | 0.338 | 0.486 | 0.338 | - | - | - | - | - |
| AAV8 | 0.010 | 0.133 | 0.133 | 0.856 | 0.017 | - | - | - | - |
| AAV9 | 0.017 | 0.161 | 0.235 | 1.000 | 0.041 | 0.952 | - | - | - |
| AAVrh8 | 0.010 | 0.133 | 0.338 | 0.574 | 0.321 | 0.010 | 0.049 | - | - |
| AAVrh10 | 0.010 | 0.311 | 0.410 | 0.574 | 0.600 | 0.010 | 0.028 | 0.802 | - |
| AAVrh74 | 0.010 | 0.133 | 0.321 | 0.662 | 0.133 | 0.010 | 0.133 | 0.767 | 0.492 |

Supplemental Table S9 - Wilcoxon Rank Tests - Flow Cytometry Female

**CLP**

| | AAV3b | AAV4 | AAV5 | AAV6 | AAV7 | AAV8 | AAV9 | AAVrh8 | AAVrh10 |
|---|---|---|---|---|---|---|---|---|---|
| AAV4 | 0.005 | - | - | - | - | - | - | - | - |
| AAV5 | 0.005 | 0.856 | - | - | - | - | - | - | - |
| AAV6 | 0.005 | 0.026 | 0.026 | - | - | - | - | - | - |
| AAV7 | 0.005 | 0.005 | 0.005 | 0.165 | - | - | - | - | - |
| AAV8 | 0.005 | 0.005 | 0.005 | 0.086 | 0.546 | - | - | - | - |
| AAV9 | 0.010 | 0.010 | 0.010 | 0.063 | 0.361 | 1.000 | - | - | - |
| AAVrh8 | 0.005 | 0.005 | 0.005 | 0.026 | 0.005 | 0.060 | 0.033 | - | - |
| AAVrh10 | 0.005 | 0.005 | 0.005 | 0.086 | 0.165 | 0.959 | 0.749 | 0.010 | - |
| AAVrh74 | 0.005 | 0.005 | 0.005 | 0.060 | 0.040 | 0.749 | 0.543 | 0.040 | 0.467 |

**CMP**

| | AAV3b | AAV4 | AAV5 | AAV6 | AAV7 | AAV8 | AAV9 | AAVrh8 | AAVrh10 |
|---|---|---|---|---|---|---|---|---|---|
| AAV4 | 0.004 | - | - | - | - | - | - | - | - |
| AAV5 | 0.004 | 0.367 | - | - | - | - | - | - | - |
| AAV6 | 0.004 | 0.013 | 0.004 | - | - | - | - | - | - |
| AAV7 | 0.004 | 0.004 | 0.004 | 0.455 | - | - | - | - | - |
| AAV8 | 0.004 | 0.004 | 0.004 | 0.007 | 0.007 | - | - | - | - |
| AAV9 | 0.004 | 0.004 | 0.004 | 0.058 | 0.170 | 0.089 | - | - | - |
| AAVrh8 | 0.004 | 0.004 | 0.004 | 0.022 | 0.013 | 0.300 | 0.856 | - | - |
| AAVrh10 | 0.004 | 0.004 | 0.004 | 0.022 | 0.007 | 0.170 | 1.000 | 0.959 | - |
| AAVrh74 | 0.004 | 0.004 | 0.004 | 0.007 | 0.004 | 0.520 | 0.520 | 0.367 | 0.475 |

**GMP**

| | AAV3b | AAV4 | AAV5 | AAV6 | AAV7 | AAV8 | AAV9 | AAVrh8 | AAVrh10 |
|---|---|---|---|---|---|---|---|---|---|
| AAV4 | 0.007 | - | - | - | - | - | - | - | - |
| AAV5 | 0.007 | 0.412 | - | - | - | - | - | - | - |
| AAV6 | 0.007 | 0.011 | 0.005 | - | - | - | - | - | - |
| AAV7 | 0.007 | 0.007 | 0.005 | 0.699 | - | - | - | - | - |
| AAV8 | 0.007 | 0.005 | 0.005 | 0.011 | 0.011 | - | - | - | - |
| AAV9 | 0.007 | 0.005 | 0.005 | 0.152 | 0.077 | 0.077 | - | - | - |
| AAVrh8 | 0.007 | 0.005 | 0.005 | 0.007 | 0.005 | 0.051 | 0.007 | - | - |
| AAVrh10 | 0.007 | 0.005 | 0.005 | 0.007 | 0.005 | 0.412 | 0.011 | 0.412 | - |
| AAVrh74 | 0.007 | 0.005 | 0.005 | 0.005 | 0.005 | 0.005 | 0.005 | 0.602 | 0.202 |

**MEP**

| | AAV3b | AAV4 | AAV5 | AAV6 | AAV7 | AAV8 | AAV9 | AAVrh8 | AAVrh10 |
|---|---|---|---|---|---|---|---|---|---|
| AAV4 | 0.004 | - | - | - | - | - | - | - | - |
| AAV5 | 0.004 | 0.285 | - | - | - | - | - | - | - |
| AAV6 | 0.004 | 0.021 | 0.084 | - | - | - | - | - | - |
| AAV7 | 0.004 | 0.007 | 0.035 | 0.749 | - | - | - | - | - |
| AAV8 | 0.004 | 0.004 | 0.054 | 1.000 | 0.856 | - | - | - | - |
| AAV9 | 0.004 | 0.004 | 0.007 | 1.000 | 0.749 | 0.219 | - | - | - |
| AAVrh8 | 0.004 | 0.004 | 0.004 | 0.021 | 0.004 | 0.004 | 0.004 | - | - |
| AAVrh10 | 0.004 | 0.004 | 0.004 | 0.004 | 0.004 | 0.004 | 0.004 | 0.165 | - |
| AAVrh74 | 0.004 | 0.004 | 0.004 | 0.007 | 0.004 | 0.004 | 0.004 | 0.357 | 0.662 |

Supplemental Table S9 - Wilcoxon Rank Tests - Flow Cytometry Female

**LT-HSC**

| | AAV3b | AAV4 | AAV5 | AAV6 | AAV7 | AAV8 | AAV9 | AAVrh8 | AAVrh10 |
|---|---|---|---|---|---|---|---|---|---|
| AAV4 | 0.008 | - | - | - | - | - | - | - | - |
| AAV5 | 0.008 | 0.913 | - | - | - | - | - | - | - |
| AAV6 | 0.008 | 0.123 | 0.170 | - | - | - | - | - | - |
| AAV7 | 0.008 | 0.010 | 0.010 | 0.559 | - | - | - | - | - |
| AAV8 | 0.008 | 0.010 | 0.010 | 0.065 | 0.010 | - | - | - | - |
| AAV9 | 0.008 | 0.010 | 0.010 | 0.662 | 0.749 | 0.010 | - | - | - |
| AAVrh8 | 0.008 | 0.008 | 0.008 | 0.039 | 0.039 | 0.271 | 0.016 | - | - |
| AAVrh10 | 0.008 | 0.008 | 0.008 | 0.089 | 0.039 | 0.216 | 0.026 | 0.749 | - |
| AAVrh74 | 0.008 | 0.008 | 0.008 | 0.089 | 0.026 | 0.271 | 0.039 | 0.937 | 0.937 |

**ST-HSC**

| | AAV3b | AAV4 | AAV5 | AAV6 | AAV7 | AAV8 | AAV9 | AAVrh8 | AAVrh10 |
|---|---|---|---|---|---|---|---|---|---|
| AAV4 | 0.008 | - | - | - | - | - | - | - | - |
| AAV5 | 0.082 | 0.937 | - | - | - | - | - | - | - |
| AAV6 | 0.008 | 0.008 | 0.018 | - | - | - | - | - | - |
| AAV7 | 0.008 | 0.006 | 0.006 | 0.040 | - | - | - | - | - |
| AAV8 | 0.008 | 0.006 | 0.006 | 0.006 | 0.091 | - | - | - | - |
| AAV9 | 0.008 | 0.006 | 0.006 | 0.443 | 0.062 | 0.006 | - | - | - |
| AAVrh8 | 0.008 | 0.006 | 0.006 | 0.006 | 0.532 | 0.357 | 0.006 | - | - |
| AAVrh10 | 0.008 | 0.006 | 0.006 | 0.127 | 0.732 | 0.024 | 0.357 | 0.170 | - |
| AAVrh74 | 0.008 | 0.006 | 0.006 | 0.186 | 0.357 | 0.024 | 0.631 | 0.170 | 0.937 |

**MPP2**

| | AAV3b | AAV4 | AAV5 | AAV6 | AAV7 | AAV8 | AAV9 | AAVrh8 | AAVrh10 |
|---|---|---|---|---|---|---|---|---|---|
| AAV4 | 0.007 | - | - | - | - | - | - | - | - |
| AAV5 | 0.007 | 0.007 | - | - | - | - | - | - | - |
| AAV6 | 0.007 | 0.937 | 0.023 | - | - | - | - | - | - |
| AAV7 | 0.007 | 0.054 | 0.007 | 0.520 | - | - | - | - | - |
| AAV8 | 0.007 | 0.007 | 0.007 | 0.007 | 0.007 | - | - | - | - |
| AAV9 | 0.007 | 0.007 | 0.007 | 0.113 | 0.156 | 0.035 | - | - | - |
| AAVrh8 | 0.007 | 0.007 | 0.007 | 0.007 | 0.007 | 0.270 | 0.035 | - | - |
| AAVrh10 | 0.007 | 0.007 | 0.007 | 0.007 | 0.013 | 0.070 | 0.113 | 0.340 | - |
| AAVrh74 | 0.007 | 0.007 | 0.007 | 0.013 | 0.007 | 0.270 | 0.035 | 0.715 | 0.715 |

**MPP3**

| | AAV3b | AAV4 | AAV5 | AAV6 | AAV7 | AAV8 | AAV9 | AAVrh8 | AAVrh10 |
|---|---|---|---|---|---|---|---|---|---|
| AAV4 | 0.009 | - | - | - | - | - | - | - | - |
| AAV5 | 0.009 | 0.213 | - | - | - | - | - | - | - |
| AAV6 | 0.009 | 0.009 | 0.009 | - | - | - | - | - | - |
| AAV7 | 0.009 | 0.008 | 0.008 | 0.270 | - | - | - | - | - |
| AAV8 | 0.009 | 0.008 | 0.008 | 0.040 | 0.015 | - | - | - | - |
| AAV9 | 0.009 | 0.008 | 0.008 | 0.340 | 0.937 | 0.024 | - | - | - |
| AAVrh8 | 0.009 | 0.008 | 0.008 | 0.060 | 0.015 | 0.837 | 0.008 | - | - |
| AAVrh10 | 0.009 | 0.008 | 0.008 | 0.165 | 0.127 | 0.076 | 0.213 | 0.024 | - |
| AAVrh74 | 0.009 | 0.008 | 0.008 | 0.165 | 0.060 | 0.270 | 0.165 | 0.412 | 0.412 |

Supplemental Table S9 - Wilcoxon Rank Tests - Flow Cytometry Female

**MPP4**

| | AAV3b | AAV4 | AAV5 | AAV6 | AAV7 | AAV8 | AAV9 | AAVrh8 | AAVrh10 |
|---|---|---|---|---|---|---|---|---|---|
| AAV4 | 0.007 | - | - | - | - | - | - | - | - |
| AAV5 | 0.007 | 0.007 | - | - | - | - | - | - | - |
| AAV6 | 0.007 | 0.033 | 0.007 | - | - | - | - | - | - |
| AAV7 | 0.007 | 0.007 | 0.007 | 0.264 | - | - | - | - | - |
| AAV8 | 0.007 | 0.007 | 0.007 | 0.007 | 0.007 | - | - | - | - |
| AAV9 | 0.007 | 0.007 | 0.007 | 0.264 | 0.937 | 0.007 | - | - | - |
| AAVrh8 | 0.007 | 0.007 | 0.007 | 0.021 | 0.021 | 0.207 | 0.007 | - | - |
| AAVrh10 | 0.007 | 0.007 | 0.007 | 0.033 | 0.110 | 0.033 | 0.110 | 0.520 | - |
| AAVrh74 | 0.007 | 0.007 | 0.007 | 0.013 | 0.007 | 0.033 | 0.007 | 0.937 | 0.616 |

**CD4+ T cells**

| | AAV3b | AAV4 | AAV5 | AAV6 | AAV7 | AAV8 | AAV9 | AAVrh8 | AAVrh10 |
|---|---|---|---|---|---|---|---|---|---|
| AAV4 | 0.005 | - | - | - | - | - | - | - | - |
| AAV5 | 0.005 | 0.455 | - | - | - | - | - | - | - |
| AAV6 | 0.005 | 0.025 | 0.732 | - | - | - | - | - | - |
| AAV7 | 0.005 | 0.008 | 0.015 | 0.005 | - | - | - | - | - |
| AAV8 | 0.005 | 0.008 | 0.008 | 0.005 | 0.546 | - | - | - | - |
| AAV9 | 0.005 | 0.005 | 0.005 | 0.005 | 0.732 | 0.005 | - | - | - |
| AAVrh8 | 0.005 | 0.127 | 0.062 | 0.008 | 0.175 | 0.367 | 0.008 | - | - |
| AAVrh10 | 0.005 | 0.062 | 0.062 | 0.005 | 0.127 | 0.231 | 0.005 | 1.000 | - |
| AAVrh74 | 0.005 | 0.008 | 0.015 | 0.005 | 0.646 | 0.959 | 0.094 | 0.292 | 0.292 |

**CD8+ T cells**

| | AAV3b | AAV4 | AAV5 | AAV6 | AAV7 | AAV8 | AAV9 | AAVrh8 | AAVrh10 |
|---|---|---|---|---|---|---|---|---|---|
| AAV4 | 0.005 | - | - | - | - | - | - | - | - |
| AAV5 | 0.010 | 0.005 | - | - | - | - | - | - | - |
| AAV6 | 0.005 | 0.877 | 0.386 | - | - | - | - | - | - |
| AAV7 | 0.005 | 0.031 | 0.005 | 0.492 | - | - | - | - | - |
| AAV8 | 0.005 | 0.031 | 0.005 | 0.574 | 0.464 | - | - | - | - |
| AAV9 | 0.005 | 0.005 | 0.005 | 0.492 | 0.959 | 0.229 | - | - | - |
| AAVrh8 | 0.005 | 0.122 | 0.005 | 0.574 | 0.959 | 0.492 | 1.000 | - | - |
| AAVrh10 | 0.005 | 0.005 | 0.005 | 0.464 | 0.492 | 0.031 | 0.081 | 0.492 | - |
| AAVrh74 | 0.005 | 0.005 | 0.005 | 0.492 | 0.787 | 0.168 | 0.877 | 0.787 | 0.386 |

**B cells**

| | AAV3b | AAV4 | AAV5 | AAV6 | AAV7 | AAV8 | AAV9 | AAVrh8 | AAVrh10 |
|---|---|---|---|---|---|---|---|---|---|
| AAV4 | 0.435 | - | - | - | - | - | - | - | - |
| AAV5 | 0.373 | 0.944 | - | - | - | - | - | - | - |
| AAV6 | 0.373 | 1.000 | 1.000 | - | - | - | - | - | - |
| AAV7 | 0.052 | 0.078 | 0.283 | 0.052 | - | - | - | - | - |
| AAV8 | 0.024 | 0.035 | 0.337 | 0.024 | 0.850 | - | - | - | - |
| AAV9 | 0.024 | 0.024 | 0.233 | 0.024 | 1.000 | 0.642 | - | - | - |
| AAVrh8 | 0.116 | 0.283 | 0.642 | 0.172 | 0.757 | 0.373 | 0.283 | - | - |
| AAVrh10 | 0.024 | 0.035 | 0.435 | 0.024 | 1.000 | 0.850 | 0.337 | 0.373 | - |
| AAVrh74 | 0.035 | 0.078 | 0.373 | 0.024 | 1.000 | 0.944 | 0.435 | 0.337 | 1.000 |

Supplemental Table S9 - Wilcoxon Rank Tests - Flow Cytometry Female

**Eosinophils**

| | AAV3b | AAV4 | AAV5 | AAV6 | AAV7 | AAV8 | AAV9 | AAVrh8 | AAVrh10 |
|---|---|---|---|---|---|---|---|---|---|
| AAV4 | 0.019 | - | - | - | - | - | - | - | - |
| AAV5 | 0.019 | 0.086 | - | - | - | - | - | - | - |
| AAV6 | 0.019 | 0.065 | 0.679 | - | - | - | - | - | - |
| AAV7 | 0.086 | 0.623 | 0.959 | 0.856 | - | - | - | - | - |
| AAV8 | 0.086 | 0.117 | 0.161 | 0.537 | 0.802 | - | - | - | - |
| AAV9 | 0.019 | 0.019 | 0.019 | 0.117 | 0.299 | 0.086 | - | - | - |
| AAVrh8 | 0.019 | 0.045 | 0.537 | 0.679 | 0.767 | 0.623 | 0.279 | - | - |
| AAVrh10 | 0.019 | 0.019 | 0.019 | 0.019 | 0.045 | 0.028 | 0.279 | 0.086 | - |
| AAVrh74 | 0.019 | 0.065 | 0.380 | 0.679 | 0.767 | 0.679 | 0.220 | 1.000 | 0.028 |

**CD11c+ monocytes**

| | AAV3b | AAV4 | AAV5 | AAV6 | AAV7 | AAV8 | AAV9 | AAVrh8 | AAVrh10 |
|---|---|---|---|---|---|---|---|---|---|
| AAV4 | 0.161 | - | - | - | - | - | - | - | - |
| AAV5 | 0.328 | 1.000 | - | - | - | - | - | - | - |
| AAV6 | 0.269 | 0.492 | 0.767 | - | - | - | - | - | - |
| AAV7 | 0.398 | 0.574 | 0.574 | 0.662 | - | - | - | - | - |
| AAV8 | 0.122 | 0.269 | 0.662 | 0.220 | 0.161 | - | - | - | - |
| AAV9 | 0.065 | 0.065 | 0.269 | 0.032 | 0.065 | 0.122 | - | - | - |
| AAVrh8 | 0.032 | 0.032 | 0.122 | 0.032 | 0.052 | 0.093 | 0.328 | - | - |
| AAVrh10 | 0.032 | 0.032 | 0.065 | 0.032 | 0.032 | 0.093 | 0.328 | 0.981 | - |
| AAVrh74 | 0.032 | 0.032 | 0.065 | 0.032 | 0.032 | 0.122 | 0.398 | 1.000 | 0.981 |

**Ly6C+ monocytes**

| | AAV3b | AAV4 | AAV5 | AAV6 | AAV7 | AAV8 | AAV9 | AAVrh8 | AAVrh10 |
|---|---|---|---|---|---|---|---|---|---|
| AAV4 | 0.011 | - | - | - | - | - | - | - | - |
| AAV5 | 0.016 | 0.024 | - | - | - | - | - | - | - |
| AAV6 | 0.011 | 0.500 | 0.016 | - | - | - | - | - | - |
| AAV7 | 0.011 | 0.010 | 0.010 | 0.024 | - | - | - | - | - |
| AAV8 | 0.011 | 0.011 | 0.010 | 0.016 | 1.000 | - | - | - | - |
| AAV9 | 0.011 | 0.010 | 0.010 | 0.016 | 0.959 | 0.646 | - | - | - |
| AAVrh8 | 0.011 | 0.231 | 0.039 | 0.559 | 0.091 | 0.060 | 0.127 | - | - |
| AAVrh10 | 0.011 | 0.010 | 0.010 | 0.010 | 0.479 | 0.749 | 0.300 | 0.024 | - |
| AAVrh74 | 0.011 | 0.175 | 0.010 | 0.856 | 0.011 | 0.024 | 0.011 | 0.646 | 0.010 |

**Neutrophils**

| | AAV3b | AAV4 | AAV5 | AAV6 | AAV7 | AAV8 | AAV9 | AAVrh8 | AAVrh10 |
|---|---|---|---|---|---|---|---|---|---|
| AAV4 | 0.046 | - | - | - | - | - | - | - | - |
| AAV5 | 0.014 | 0.492 | - | - | - | - | - | - | - |
| AAV6 | 0.014 | 0.023 | 0.122 | - | - | - | - | - | - |
| AAV7 | 0.014 | 0.023 | 0.038 | 0.662 | - | - | - | - | - |
| AAV8 | 0.077 | 0.122 | 0.122 | 0.449 | 0.662 | - | - | - | - |
| AAV9 | 0.014 | 0.014 | 0.014 | 0.122 | 0.373 | 0.749 | - | - | - |
| AAVrh8 | 0.014 | 0.014 | 0.014 | 0.220 | 0.590 | 1.000 | 0.492 | - | - |
| AAVrh10 | 0.014 | 0.014 | 0.014 | 0.492 | 0.783 | 0.492 | 0.161 | 0.492 | - |
| AAVrh74 | 0.014 | 0.014 | 0.014 | 0.161 | 0.449 | 0.749 | 0.837 | 0.662 | 0.289 |

Supplemental Table S9 - Wilcoxon Rank Tests - Flow Cytometry Female

**NK cells**

| | AAV3b | AAV4 | AAV5 | AAV6 | AAV7 | AAV8 | AAV9 | AAVrh8 | AAVrh10 |
|---|---|---|---|---|---|---|---|---|---|
| AAV4 | 0.005 | - | - | - | - | - | - | - | - |
| AAV5 | 0.005 | 0.015 | - | - | - | - | - | - | - |
| AAV6 | 0.005 | 0.005 | 0.005 | - | - | - | - | - | - |
| AAV7 | 0.005 | 0.040 | 0.005 | 0.024 | - | - | - | - | - |
| AAV8 | 0.005 | 0.008 | 0.005 | 0.005 | 0.559 | - | - | - | - |
| AAV9 | 0.005 | 0.024 | 0.005 | 0.005 | 0.818 | 0.060 | - | - | - |
| AAVrh8 | 0.005 | 0.008 | 0.005 | 0.005 | 0.631 | 0.528 | 0.357 | - | - |
| AAVrh10 | 0.005 | 0.008 | 0.005 | 0.005 | 0.559 | 0.507 | 0.131 | 0.732 | - |
| AAVrh74 | 0.005 | 0.060 | 0.008 | 0.005 | 0.818 | 0.410 | 0.631 | 0.631 | 0.559 |
